# Comparison of sparse biclustering algorithms for gene expression datasets

**DOI:** 10.1101/2020.12.15.422852

**Authors:** Kath Nicholls, Chris Wallace

## Abstract

Gene clustering and sample clustering are commonly used to find patterns in gene expression datasets. However, in heterogeneous samples (e.g. different tissues or disease states), genes may cluster differently. Biclustering algorithms aim to solve this issue by performing sample clustering and gene clustering simultaneously. Existing reviews of biclustering algorithms have yet to include a number of more recent algorithms and have based comparisons on simplistic simulated datasets without specific evaluation of biclusters in real datasets, using less robust metrics.

In this study we compared four classes of sparse biclustering algorithms on a range of simulated and real datasets. In particular we use a knockout mouse RNA-seq dataset to evaluate each algorithm’s ability to simultaneously cluster genes and cluster samples across multiple tissues. We found that Bayesian algorithms with strict sparsity constraints had high accuracy on the simulated datasets and didn’t require any post-processing, but were considerably slower than other algorithm classes. We assessed whether non-negative matrix factorisation algorithms can be repurposed for biclustering and found that, although the raw output was poor, after using a sparsity-inducing post-processing procedure we introduce, one such algorithm was one of the most highly ranked on real datasets. We also exhibit the limitations of biclustering algorithms by varying the complexity of simulated datasets. The algorithms generally struggled on simulated datasets with a large number of implanted factors, or with a large number of genes. In real datasets, the algorithms rarely returned clusters containing samples from multiple tissues, which highlights the need for further thought in the design and analysis of multi-tissue studies to avoid differences between tissues dominating the analysis.

Code to run the analysis is available at https://github.com/nichollskc/biclust_comp, including wrappers for each algorithm, implementations of evaluation metrics, and code to simulate datasets and perform pre- and post-processing. The full tables of results are available at https://doi.org/10.5281/zenodo.4317556

## 1 Introduction

Clustering can be used in two main ways to analyse gene expression datasets [1]. The first is to cluster the samples, finding groups of samples that have similar expression in all genes. This can be used, for example, to find subgroups of disease [2]. The second is to cluster the genes, finding groups of genes that have similar expression across all samples. Finding such groups of genes has many useful applications such as inferring function using guilt by association and inferring regulatory relationships [3].

Instead of clustering only samples or only genes, biclustering algorithms find groups of samples that have similar expression in some subset of the genes, effectively clustering both genes and samples simultaneously. Such a group is called a *bicluster* and we say that the bicluster consists of a set of samples and a set of genes. Biclustering has three main advantages over normal clustering. Firstly, it can discover meaningful groups that would not be detected using normal clustering; in complex datasets, many interesting groupings of genes will not hold across all samples. For example, we might expect genes to cluster differently in different cell types. Secondly, it provides a link between sets of genes and sample traits such as disease or sex. For example, if a biclustering algorithm returns a bicluster consisting of all the samples from patients with a given disease and a small set of genes then we can hypothesise that the set of genes might have biological importance for the disease. Finally, biclustering algorithms are additive, allowing the algorithm to learn biclusters corresponding to confounders, such as batch or sex, and adjust for these confounders whilst simultaneously extracting biologically interesting biclusters. In this study we have focused on identifying algorithms that should be able to identify sparse biclusters in a complex bulk RNA-seq dataset, such as one including samples from multiple cell types.

There exist previous reviews of biclustering algorithms [4–7], but we hope to improve on them in the following ways. First, we include new classes of algorithm yet to be considered in independent comparison studies. In particular, we include non-negative matrix factorisation algorithms, which we believe can be repurposed for biclustering, tensor factorisation algorithms, which aim to improve performance by sharing information across tissues, and two Bayesian algorithms which allow for a mixture of sparse and dense biclusters. Second, we use more robust metrics. Horta and Campello investigated metrics used to evaluate similarity between biclusterings, and found problems with many of the metrics used by previous comparison papers [8]. In this study we use one of the two metrics recommended by Horta and Campello, which was shown to satisfy all but one of their criteria. Third, we narrow the gap between real and simulated datasets. Previous reviews have often used unrealistically simplistic simulated datasets, such as using only *K* = 1, 2, 3, 4, 5 biclusters, leading to discrepancies between the conclusions they draw on simulated and on real datasets [6]. In this study we simulate datasets from a wider range of complexities, including datasets closer in complexity to real datasets than those included in previous reviews. A final key flaw of existing comparison studies is the lack of evaluation of biclustering ability on real datasets. In the absence of known structure in the real gene expression datasets used for evaluation, previous reviews have evaluated sample clustering ability and gene clustering ability separately. We carefully chose a knockout mouse RNA-seq dataset that allows linked analysis of sample clustering and gene clustering, thus allowing direct evaluation of biclustering on real datasets.

## 2 Methods

Here we discuss the algorithms compared, the datasets they are tested on and the evaluation metrics used to score their performance. Similar to previous reviews, we use a mixture of simulated and real datasets. Simulated data is important as it allows more precise evaluation of performance, since the true structure of the data is known. However, it is difficult to exactly mimic the noise and structure of real gene expression datasets, so it is also important to see whether the algorithms can handle the noise structure of real datasets.

### 2.1 Algorithms compared

We chose most promising algorithms from four classes of algorithm, focusing on sparse algorithms (Table 1).

**Table 1:**
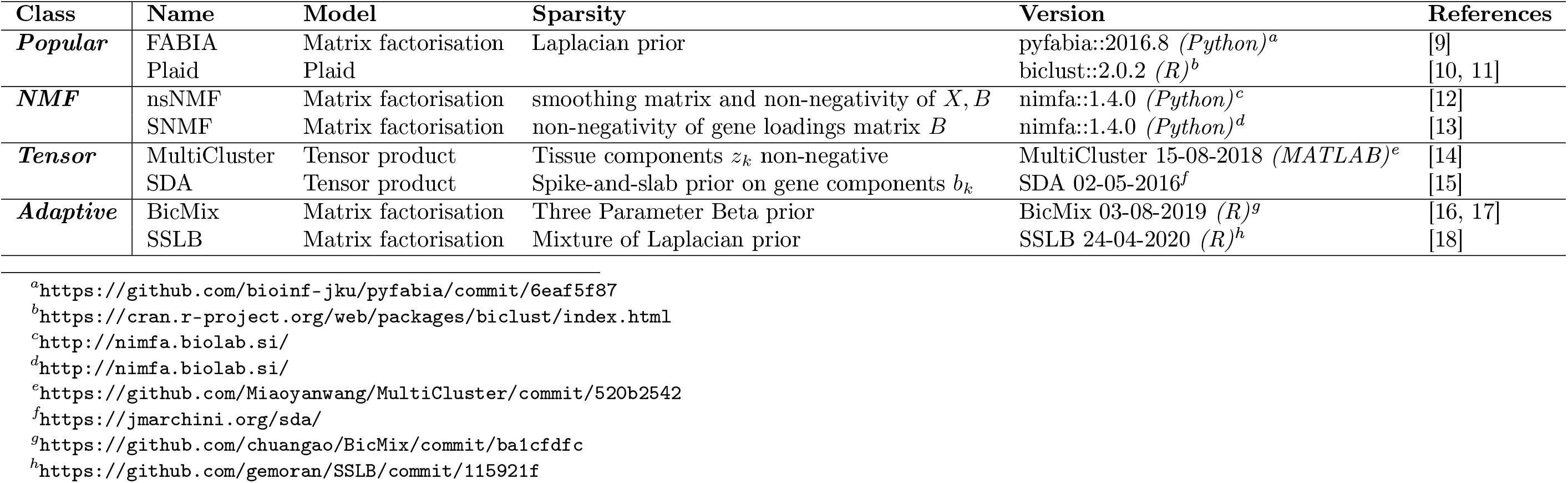
Summary of algorithms included in comparison. Algorithms are listed in groups: popular algorithms which have been included in previous reviews (*Popular*), non-negative matrix factorisations algorithms (*NMF*), tensor factorisation algorithms (*Tensor*) and Bayesian algorithms allowing for a mixture of sparse and dense biclusters, with strength of sparsity constraints adapting to the bicluster (*Adaptive*). With the exception of Plaid (Section SI.1), algorithms either factorise the gene expression matrix *Y* as *Y* = *XB^T^*+*ε* (matrix factorisation), where *X* is the samples loading matrix, and *B* is the gene loading matrix, or write it as a tensor product 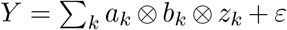 (tensor product), where *a_k_* gives the loadings for bicluster *k* for the individuals, *b_k_* gives the loadings for genes and *z_k_* gives the loadings for the tissues. Some algorithms use one language to implement the algorithm, and provide a ‘wrapper’ in another language. Where this occurs, the language given in the ‘Version’ column is the language used to interact with the algorithm (the wrapper), rather than the language that the implementation uses.

We define a matrix 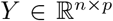 where entry *Y_ij_* gives the expression of gene *j* in sample *i*. This can either be the raw read count from an RNA-seq experiment, or a normalised count which has been adjusted for sample-specific effects such as library size, or gene-specific effects such as mean expression level. The typical approach to biclustering is to factorise this matrix as a product of two sparse matrices 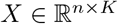, which we call the sample loadings matrix, and 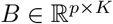, which we call the gene loadings matrix, with error matrix *ε*:

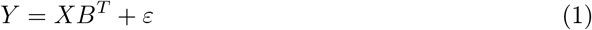

The individual algorithms are described in detail in Section S1. Here we discuss why each algorithm was chosen for inclusion in this study and group the algorithms as *Popular, Adaptive, NMF* and *Tensor*.

#### 2.1.1 Popular algorithms

We include two algorithms which have been included in previous reviews, which we use as a baseline to allow relative performance to be related to other comparison studies. FABIA [9] is a Bayesian algorithm using sparsity-inducing priors, included in a number of previous comparisons [3, 6, 7, 19]. Although our study focuses on sparse biclustering algorithms, we chose to also include Plaid [10, 11] even though it does not enforce sparsity, as it has often appeared as one of the better performing algorithms in other studies [6, 7] and its inclusion thus provides a helpful link to these studies.

#### 2.1.2 Adaptive Bayesian algorithms

Like FABIA, BicMix [16, 17] and SSLB [18] use sparsity-inducing priors. The key difference with BicMix and SSLB is that they allow for both sparse and dense biclusters, and adapt the sparsity constraints to each bicluster. Neither has been included in previous comparisons but they have been compared against each other and against FABIA in the paper introducing SSLB, where both achieved much greater sparsity and accuracy than FABIA.

#### 2.1.3 Non-negative matrix factorisation

Non-negative matrix factorisation (NMF) algorithms in general are not designed for biclustering, but since biclustering can be described as sparse matrix factorisation, NMF algorithms can recover biclusters if they use sufficiently strong sparsity constraints. We chose to include two examples of such algorithms: SNMF [13] and nsNMF [12]. The main advantage we expect these algorithms will have is speed, as they are computationally much simpler than many of the others included in this study.

#### 2.1.4 Tensor algorithms

When applying an algorithm to data from multiple cell types, a natural extension to the twodimensional algorithms presented so far is a three-dimensional algorithm which exploits similarity between corresponding samples in different cell types. We chose to include two algorithms which attempt this: SDA [15] and MultiCluster [14].

### 2.2 Algorithm parameters

The algorithms evaluated here, outlined in Table 1, have many parameters which can be tuned. Before running the full analysis, we conducted a parameter sweep (Section S2) to see if there were any parameter values that consistently improved the score relative to that when the default values were used. For most algorithm parameters, there was either no clear optimal value, or the default value was optimal. Thus for most algorithms we used the default parameters throughout this study. One key exception was BicMix, which has a parameter determining whether or not each gene gets transformed to a Gaussian distribution before the algorithm runs. Changing this parameter had a dramatic but inconsistent effect, so we decided to use two versions of BicMix: BicMix, using default behaviour of not transforming genes, and BicMix-Q, which does apply the Gaussian transformation before analysis. Full discussion of our investigation of parameter sensitivity are given in Section S2.

### 2.3 Simulated datasets

We simulated individual gene expression data for each gene as a sum across biclusters of negative binomial counts. Our base model for generating a gene expression dataset with *p* genes, *m* individuals and *t* tissues and with *K* potentially overlapping biclusters is illustrated in Figure 1 and described below:

1. For each bicluster *k* = 1,…, *K*:

a. Select genes to include in bicluster *k*: first draw number of genes *g_k_* uniformly from the set 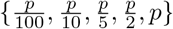 and then pick a random sample of *g_k_* genes.
b. Select individuals to include in bicluster *k*: first draw number of individuals *m_k_* uniformly from the set 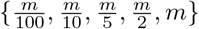 and then pick a random sample of *m_k_* individuals.
c. Select tissues to include in bicluster *k*: first draw number of tissues *t_k_* uniformly from the set {1, 2,…, *t*} and then pick a random sample of *t_k_* tissues.
d. Sample bicluster-specific mean *μ_k_* ~ Gamma(*α, β*) using *α* = 2, 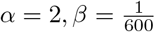.
e. Sample values in bicluster using negative binomial distribution with mean *μ_k_*, shared parameter *p* = 0.3.
2. Add together values from all biclusters
3. Add background noise using negative binomial distribution

**Figure 1:**
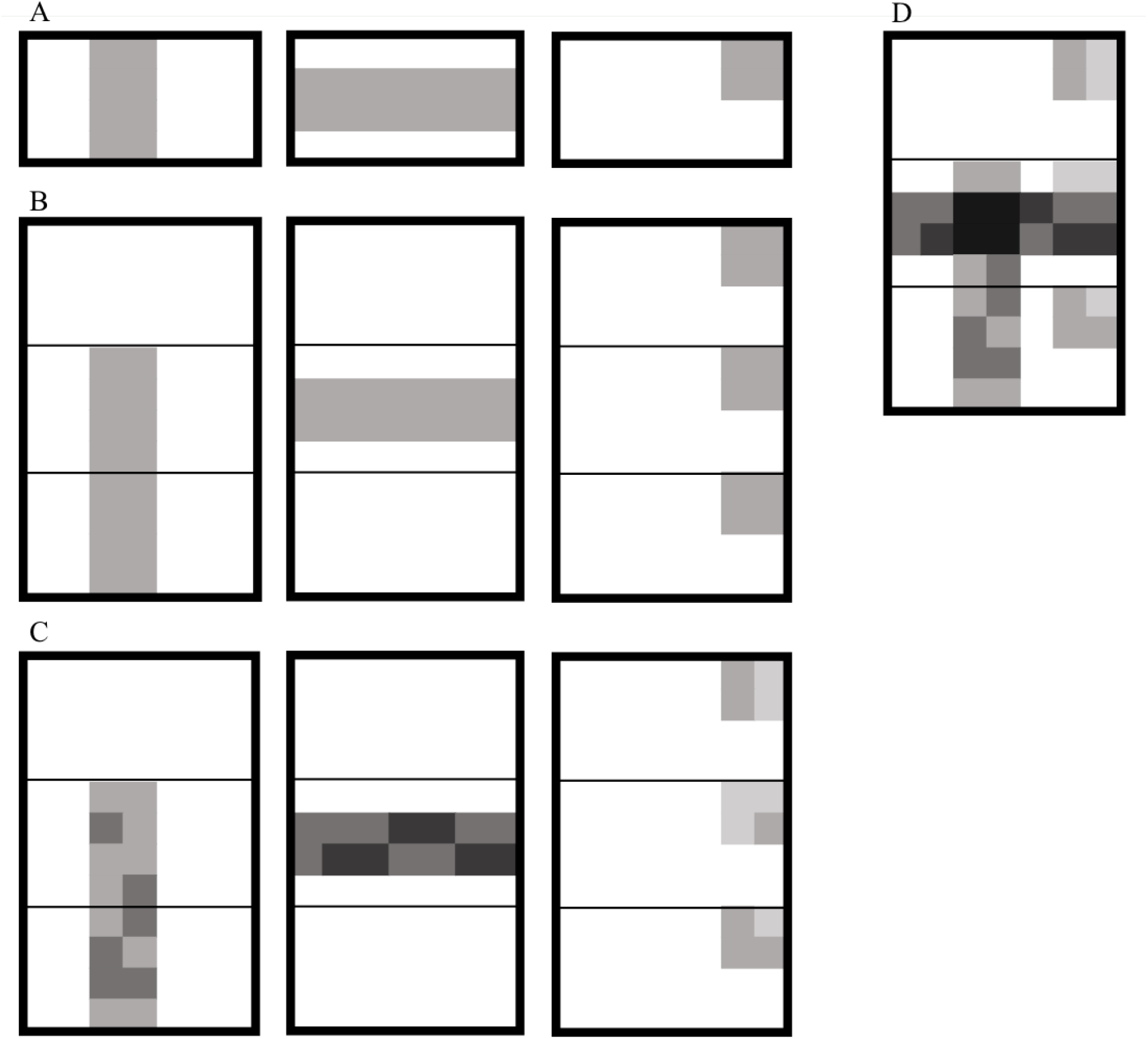
Illustration of process for simulating gene expression datasets with implanted biclusters. In this diagram (A) shows steps 1.a-1.b where membership for genes (columns) and individuals (rows) are sampled for 3 biclusters, (B) shows step 1.c for the 3 biclusters, where we extend the biclusters from size (*m_k_, g_k_*) to size (*m_k_t_k_,g_k_*) by sampling membership for tissues, (C) shows steps 1.d and 1.e where values for the bicluster members are sampled, with bicluster-specific means *μ_k_*, (D) shows step 2 where the effects from the 3 biclusters are added together.

Formally the base model is:

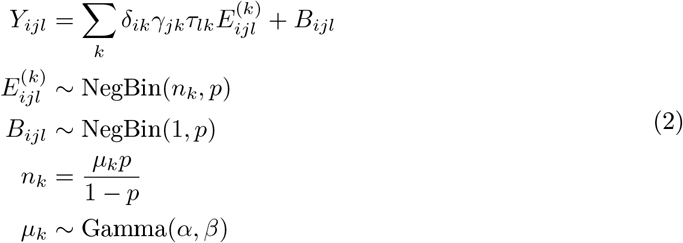

where *δ_ik_, γ_jk_* and *τ_lk_* are binary indicators of membership of individual *i*, gene *j* and tissue *l* to bicluster *k* respectively, 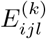 is the increase in expression of gene *i* in tissue *l* in individual *i* due to bicluster *k* and *B_ijl_* is background noise. We chose to force the genes chosen in a bicluster to belong to a contiguous block rather than allowing genes from a bicluster to be scattered freely throughout the matrix, and did the same for the tissues and individuals chosen in a bicluster. This arrangement has little impact on the generality of the data but makes it easier to visualise the datasets.

We vary the size of the dataset, the number of biclusters (which also naturally changes the amount of overlap between biclusters) and the size of biclusters. We also introduce diversity by using different noise distributions. For Gaussian noise we use 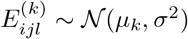 and for noiseless datasets we use 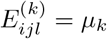, *B_jil_* = 0. A summary of the properties of all the simulated datasets is given in Table 2.

**Table 2:** Summary of simulated datasets. The attributes of the datasets are displayed in bold if they differ from the base dataset. N is the number of samples, T the number of tissues and G the number of genes in the dataset.

| Name | N | T | G | Bicluster sizes | K | Noise |
| --- | --- | --- | --- | --- | --- | --- |
| base | 10 | 10 | 1000 | mixed | 20 | $NB(n_k, 0.3)$ |
| N50-T2 | **50** | **2** | 1000 | mixed | 20 | $NB(n_k, 0.3)$ |
| N10-T20 | 10 | **20** | 1000 | mixed | 20 | $NB(n_k, 0.3)$ |
| N100-T10 | **100** | 10 | 1000 | mixed | 20 | $NB(n_k, 0.3)$ |
| N500-T10 | **500** | 10 | 1000 | mixed | 20 | $NB(n_k, 0.3)$ |
| G100 | 10 | 10 | **100** | mixed | 20 | $NB(n_k, 0.3)$ |
| G5000 | 10 | 10 | **5000** | mixed | 20 | $NB(n_k, 0.3)$ |
| large-K20 | **300** | **20** | **10000** | mixed | 20 | $NB(n_k, 0.3)$ |
| negbin-medium | 10 | 10 | 1000 | mixed | 20 | $NB(n_k, \mathbf{0.1})$ |
| negbin-high | 10 | 10 | 1000 | mixed | 20 | $NB(n_k, \mathbf{0.01})$ |
| Gaussian | 10 | 10 | 1000 | mixed | 20 | $\mathcal{N}(\mu_k, \mathbf{20^2})$ |
| Gaussian-medium | 10 | 10 | 1000 | mixed | 20 | $\mathcal{N}(\mu_k, \mathbf{100^2})$ |
| Gaussian-high | 10 | 10 | 1000 | mixed | 20 | $\mathcal{N}(\mu_k, \mathbf{300^2})$ |
| noiseless | 10 | 10 | 1000 | mixed | 20 | **No noise** |
| sparse | 10 | 10 | 1000 | **small** | 20 | $NB(n_k, 0.3)$ |
| dense | 10 | 10 | 1000 | **large** | 20 | $NB(n_k, 0.3)$ |
| sparse-square | 10 | 10 | 1000 | **small, square** | 20 | $NB(n_k, 0.3)$ |
| dense-square | 10 | 10 | 1000 | **large, square** | 20 | $NB(n_k, 0.3)$ |
| K5 | 10 | 10 | 1000 | mixed | **5** | $NB(n_k, 0.3)$ |
| K10 | 10 | 10 | 1000 | mixed | **10** | $NB(n_k, 0.3)$ |
| K50 | 10 | 10 | 1000 | mixed | **50** | $NB(n_k, 0.3)$ |
| K70 | 10 | 10 | 1000 | mixed | **70** | $NB(n_k, 0.3)$ |
| large-K100 | **300** | **20** | **10000** | mixed | **100** | $NB(n_k, 0.3)$ |
| large-K400 | **300** | **20** | **10000** | mixed | **400** | $NB(n_k, 0.3)$ |

Previous reviews have also varied simulation parameters but have often used very small ranges such as *K* = 1, 2, 3, 4, 5 [6]. Real gene expression datasets are likely to be more complex than this, so we have used larger values of *K*: most of our simulated datasets have *K* = 20 but we consider values from *K* = 5 to *K* = 400.

The *Tensor* algorithms require an explicit breakdown of the samples into tissues. By listing the samples from each tissue in turn, with individuals in the same order within each tissue, we are able to use the *Tensor* algorithms on the same datasets as the remaining algorithms, allowing direct comparison between the classes of algorithm.

### 2.4 Real datasets

A key limitation of the existing reviews of biclustering algorithms is their inability to assess *simultaneous* clustering of samples and genes on real datasets, due to the absence of known biclusters in the data. In order to have predictable bicluster structure in a real dataset, we chose to use a knockout mouse dataset [20, 21]. We proposed that a successful algorithm would recover, for each of the 106 knockout genes, a bicluster containing the roughly 20 samples where the gene was knocked out and enriched for genes that share a pathway with the knocked-out gene. Thus this dataset allows us to have some sense of its true bicluster structure.

#### 2.4.1 IMPC dataset

We use the RNA-seq dataset available on ArrayExpress under accession number E-MTAB-5131, part of the International Mouse Phenotyping Consortium (IMPC) [20, 21]. It consists of 106 knockout genotypes, from each of which are available roughly 3 replicates in each of up to 7 tissues. There are also samples from wild-type mice.

To make the study feasible for multiple algorithms in terms of computational time, we chose to restrict to a subset of genes. We restricted to the 4444 genes which share a Reactome pathway with at least one of the 106 knockout genes, found by searching the Reactome pathways [22, 23] using Mouse Mine [24]. We apply three different normalisation methods to the data: (1) library size adjustment using DESeq’s median of ratios normalisation method, (2) the log transform *x* → log (*x* + 1), which is commonly used in analysis of gene expression data and (3) Gaussian quantile normalisation so that each gene has approximate *N*(0,1) distribution.

#### 2.4.2 Tensor structure

The *Tensor* algorithms require the dataset to have three dimensions i.e. *m* individuals, *t* tissues, *p* genes rather than just *n* = *m* × *t* samples and *p* genes. We chose the 3 tissues with the most samples (liver, lung and cardiac ventricle), and the 64 genotypes with at least one sample in each of these tissues. Unfortunately we were unable to find information detailing which samples came from which specific mouse replicate so couldn’t simply include a row for each individual, a column for each gene and a layer for each tissue. Instead we pooled the samples from each genotype for each tissue individually by taking the mean of the replicates. Thus we had *m* = 64, *t* = 3 with a total of *n* = 192 samples.

This type of dataset, which we call the *tensor* dataset, can be used by all the algorithms, whereas the *non-tensor* dataset, which simply uses all *n* = 1143 samples, can’t be used by the *Tensor* algorithms.

### 2.5 Evaluation metrics

We use a range of metrics to evaluate performance of the biclustering algorithms (Table 3), which are described fully in Section S3. In particular, we made use of an extensive study of biclustering accuracy metrics [8] to choose the *clustering error* (CE) metric [25, 8] to evaluate biclustering accuracy. This was shown to satisfy all but one of the desirable properties defined by Horta and Campello, which is a great improvement on the consensus score and recovery and relevance scores commonly used to evaluate biclustering similarity.

**Table 3:** Metrics used to evaluate the biclustering algorithms, which are described more fully in Section S3. For each metric we note what information about the dataset it requires, the best and worst scores theoretically possible and briefly describe what it measures.

| **Name** | **Requires** | **Best** | **Worst** | **Measures** |
| --- | --- | --- | --- | --- |
| Clustering Error (CE) | Entire bicluster structure | 1 | 0 | Biclustering accuracy |
| Normalised Reconstruction Error (NRE) | - | 0 | 1 | Reconstruction error |
| Mean Biclustering Redundancy (MBR) | - | 0 | 1 | Similarity between biclusters within a run |
| Similarity between runs | Multiple runs | 1 | 0 | Similarity between runs |
| Sample clustering | Sample clusters | 1 | 0 | Sample clustering |
| Pathway enrichment | Pathway database | 1 | 0 | Gene clustering |
| Relevant pathway enrichment | Known link between pathways and sample clusters | 1 | 0 | Biclustering |

Most metrics used by previous reviews, including the *clustering error* metric that we intend to use for evaluation of performance on simulated datasets, cannot be used on real datasets, as they require knowledge of the entire biclustering structure of the dataset. We introduce two metrics that can be used even when nothing is known about the structure of the dataset: Normalised Reconstruction Error (NRE) and Mean Biclustering Redundancy (MBR).

## 3 Results

### 3.1 Post-processing

After looking at the raw output, we decided that we would first need to apply some postprocessing steps in order to allow meaningful comparison of the algorithms. The *Tensor* algorithms, *NMF* algorithms and FABIA returned many biclusters containing all genes and all samples (Figure S11 and Figure S12). We found that removing elements in the matrices below a certain threshold, a process we call *thresholding*, helped to reveal the biclusters within the noisy raw output. The exact process is described in Section S4. Without thresholding, the biclusters returned by FABIA, SDA and the *NMF* algorithms were highly redundant but this redundancy was reduced by thresholding with a threshold of 0.01 (Figure 2). The optimal threshold is similar for most algorithms, both on simulated datasets (Figure S13) and real datasets (Figure S14), and is largely independent of the metric used to select the threshold. In particular, we note that we could have chosen a suitable threshold using only measures available for real datasets, such as Mean Bicluster Redundancy (Figure 2) and Normalised Reconstruction Error (Figure S15).

**Figure 2:**
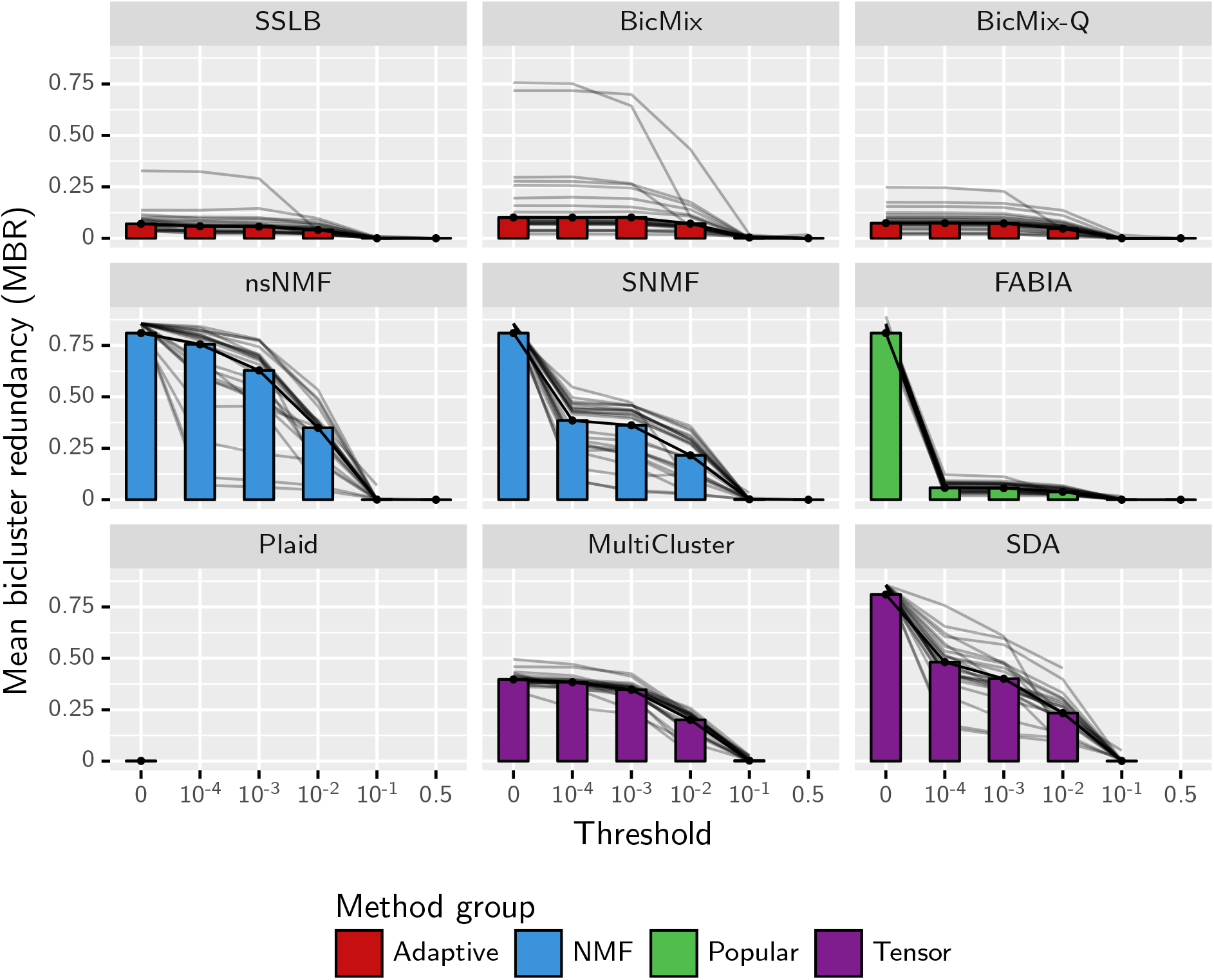
Mean bicluster redundancy (MBR) within a run plotted against the threshold for inclusion in bicluster, for simulated datasets. MBR (Section S3.3) is in the range [0, 1], and a lower value is preferred as it suggests that the algorithm is not returning biclusters which are very similar to each other. The thresholding process is described in Section S4. The median of this measure across all simulated datasets and all values of *K_init_* is shown by the bars. The grey lines show the median for each dataset type. Note that without thresholding (threshold 0) the biclusters within each run by FABIA, SDA and nsNMF are almost all identical.

It is worth highlighting that Plaid and the *Adaptive* algorithms did not require this postprocessing step, but in the interests of avoiding bias and unnecessary complications in the analysis we apply the same post-processing steps for every algorithm. The one exception is Plaid, whose implementation returns only the binary membership variables so thresholding cannot be applied. The fact that these algorithms perform well without need for post-processing is a key advantage in terms of ease of use.

### 3.2 Choice of *K_init_*

Overall, algorithms were poor at accurately recovering the right number of biclusters (Figure S19), with only FABIA and BicMix-Q showing any positive correlation between the true K and recovered K (BicMix-Q had correlation of 0.836 between true K and recovered K). Ideally we would simply use a large value of *K_init_* for all algorithms, as this is what we would do in practice on a real dataset with unknown structure. However, only Plaid and the *Adaptive* algorithms have shown that they would effectively learn the number of biclusters to include. The remaining algorithms consistently returned the same number of biclusters as they started with, so didn’t ‘learn’ K at all (Figure 3). The *Adaptive* algorithms achieve better performance when started with an overestimate of the number of biclusters (Figure S20). Thus, we use *K_init_* = *K*, the true number of biclusters for all algorithms, except for the *Adaptive* algorithms for which we use a slight overestimate of *K_init_* (*K_init_* = *K* +10, except for when *K* = 20, when we use *K_init_* = 25.). Note that our way of choosing *K_init_* is dependent on knowing the true number of biclusters, so it gives the algorithms an advantage they would not have on real datasets. However, it allows us to compare the ‘ideal’ behaviour of each algorithm. For the real datasets we use *K_init_* = 50,200 for all algorithms.

**Figure 3:**
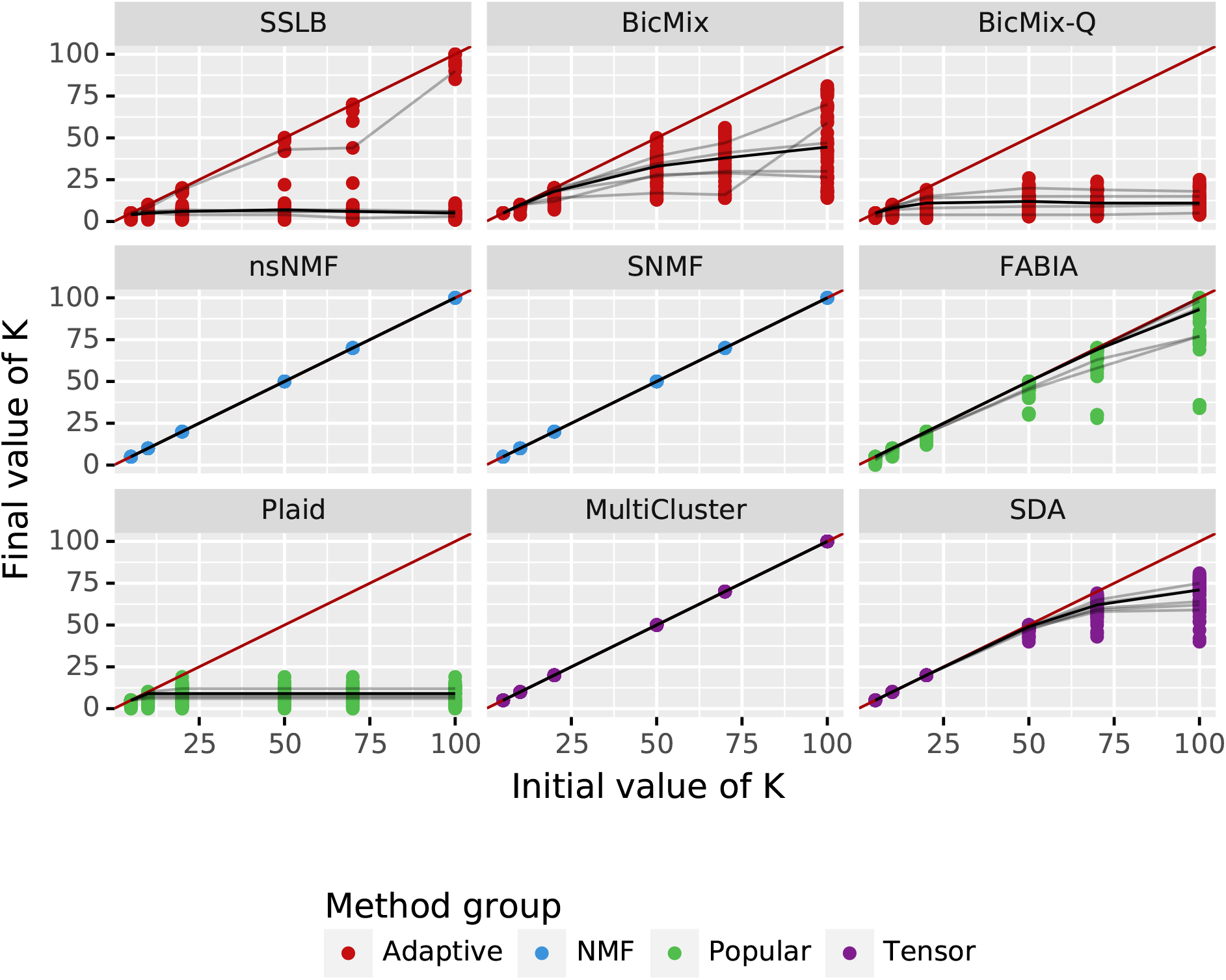
Robustness to choice of *K_init_*, shown by final value of K (after post-processing) plotted against initial value of K. There is a point for each of the 3 seeds for each dataset of type *K5, K10, base, K50* and *K70*. The red line shows the limit, where the final value of K is the same as the initial value of K. The black line joins the median 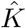 across all datasets, and the gray lines join the medians for each individual dataset. The ideal behaviour is for the line to be flat once *K_init_* exceeds the true *K*, showing that the algorithm converges to the same value of K regardless of the initial value given, as long as *K_init_* is sufficiently large. True K varies from *K* = 5 to *K* = 70. MultiCluster and *NMF* algorithms always returned the same number of factors as they started with.

### 3.3 Results on simulated datasets

The results are summarised in Table 4. Figure 4 shows the biclustering accuracy of the algorithms across all the simulated datasets. The *Adaptive* algorithms performed best, with SSLB having the best overall accuracy on simulated datasets (0.336). SNMF has the best accuracy of the non-Bayesian algorithms (0.239).

**Figure 4:**
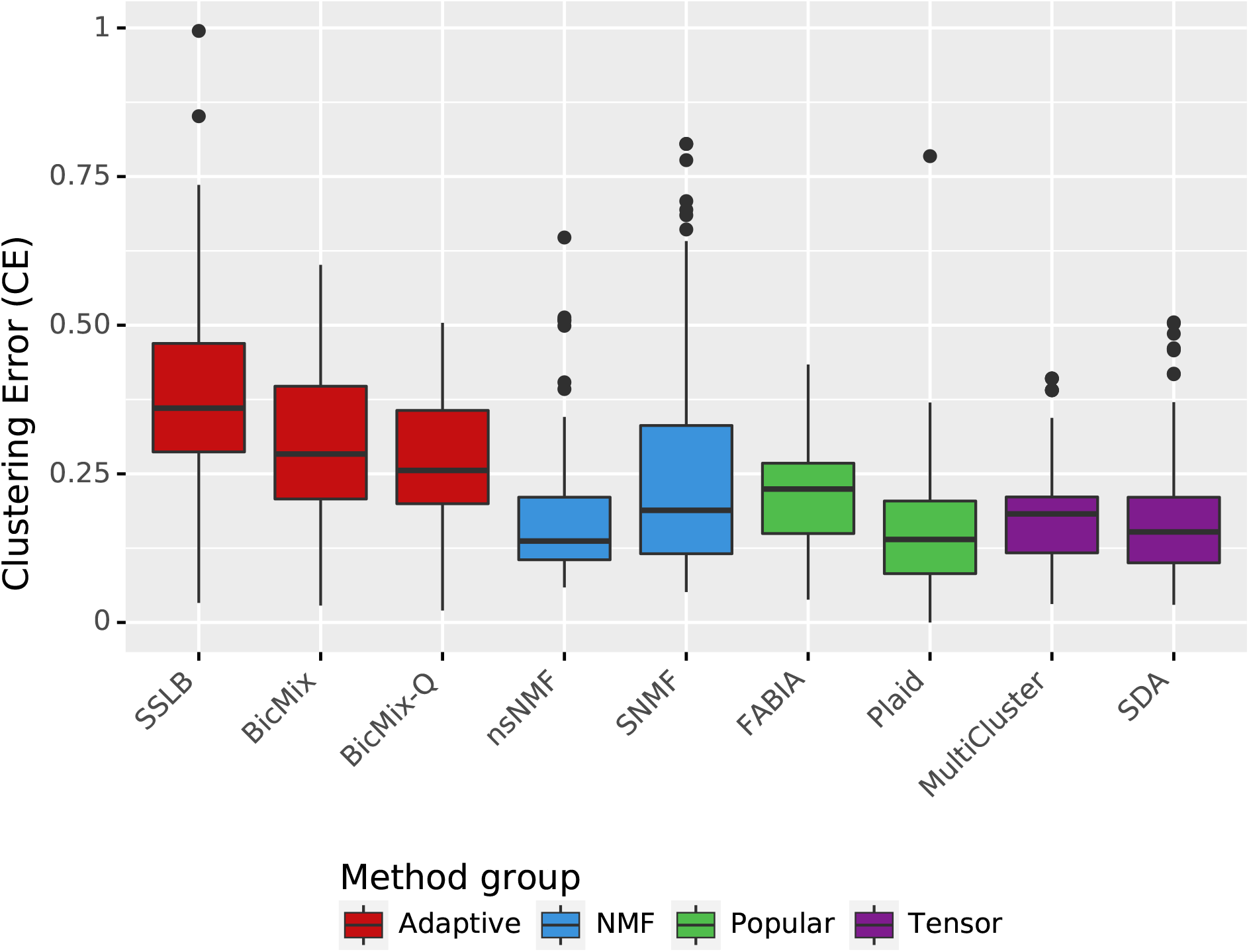
Clustering error (CE) across all simulated datasets. The score is in the range [0, 1] with larger values preferred. Datasets used are described in Table 2. *K_init_* is as described in Section 3.2 and standard thresholding has been applied. Runs that failed (Table S3) are discarded in the analysis.

**Table 4:**
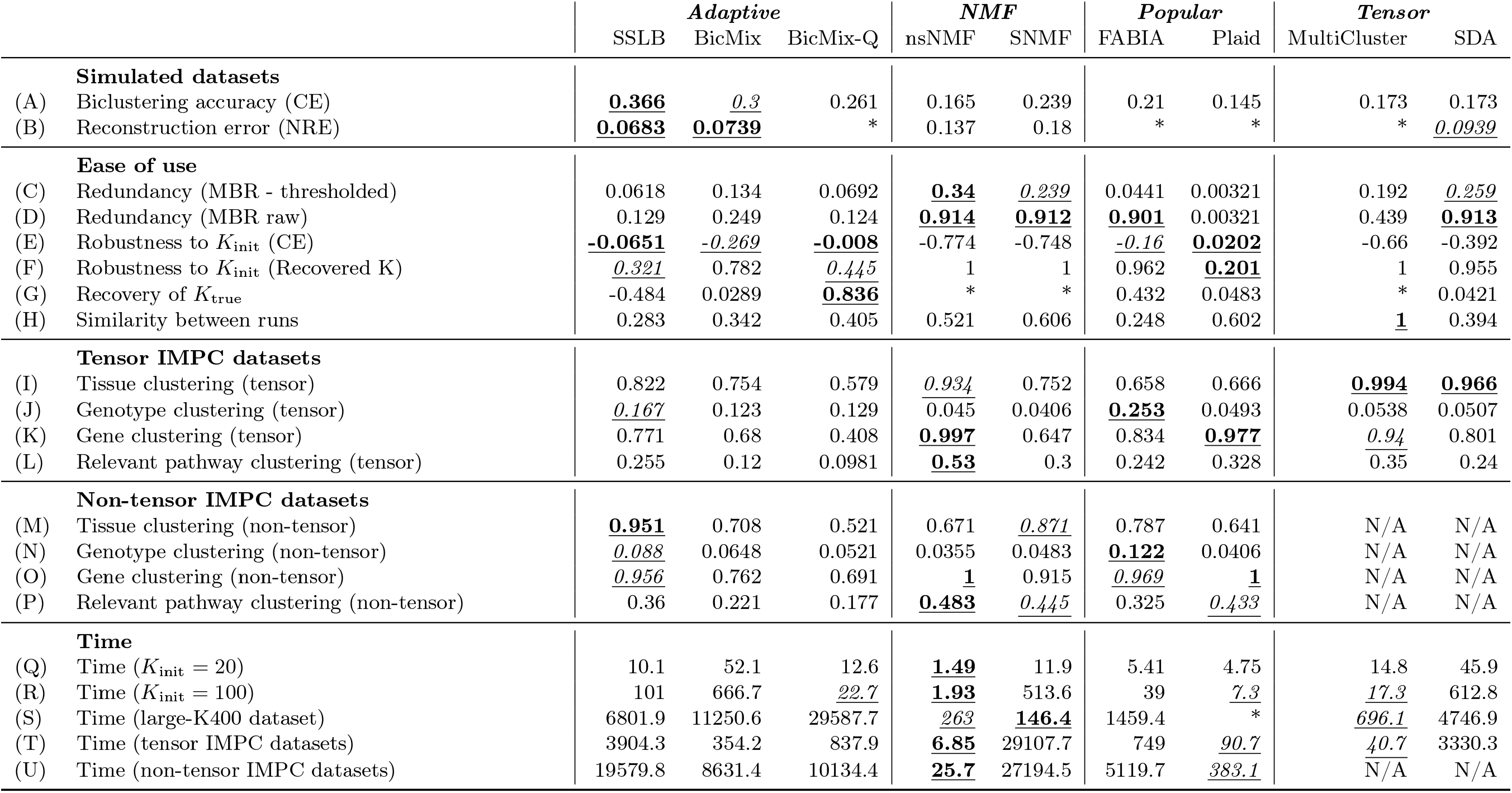
Summary of results, with the best score for each measure underlined and in bold and scores close to the best score underlined and in italics. Unless otherwise stated, the measures are given for all runs using *K_init_* as described in Section 3.2 and after the standard thresholding has been applied. Runs that failed (Table S3 and Table S4) are discarded in the analysis. *Tensor* algorithms could not be run on non-tensor datasets so these entries are marked ‘N/A’. (A) Clustering Error (CE) across all simulated datasets. (B) Reconstruction error (NRE) across all simulated datasets. Algorithms which did not return the raw values required to calculate this measure are marked with ‘*’. (C-D) Average similarity between recovered biclusters (MBR) before (raw) and after (*thresholded*) the standard thresholding has been applied. (E) Correlation between and CE, best score is 0, indicating that score was unaffected by (F) Correlation between *K_init_* and *K_recovered_*, best score is 0, indicating that the algorithm converged to the same number of biclusters regardless of *K_init_* (G) Correlation between *K*_recovered_ and *K_true_* when *K_init_* = 100, an overestimate of *K_true_*. The best score is 1. Entries marked ‘*’ correspond to algorithms that returned *K_recovered_* = *K_init_* for all runs, for which correlation could not be calculated. (H) Similarity between all ten runs on each IMPC dataset. (I-L) Performance on IMPC datasets with tensor structure with *K_init_* = 50, using measures described in Section S3. (M-P) Same as H-K but for IMPC datasets with non-tensor structure. Note that the *Tensor* algorithms were not run on the non-tensor datasets. (Q-R) Time to run in seconds on the datasets of type *K5, K10, base, K50* and *K70*, with = 20 and *K_init_* = 100 respectively. (S) Time to run in seconds for the largest simulated dataset *large-K400*, when the algorithms were run with a small number of biclusters (*K_init_* = 20 for all algorithms except SSLB, BicMix, BicMix-Q which used = 25). Plaid failed to complete any runs on this largest dataset and is marked as ‘*’. (T-U) for the largest IMPC datasets, where the algorithms used a large number of biclusters (*K_init_* = 200). Note that the *Tensor* algorithms were not run on the non-tensor datasets.

The accuracy of the algorithms generally decreases as the size of the dataset increases (Figure S21), and as the number of biclusters increases (Figure S22). We had expected algorithms to perform better when there were fewer biclusters, which is the case for SSLB and the *NMF* algorithms. However, FABIA, BicMix and BicMix-Q have poor accuracy on the datasets with small number of biclusters. For the very largest datasets (*large-K100* and *large-K400*) many algorithms took a long time to run and only MultiCluster and nsNMF completed runs within 12 hours when using *K_init_* close to 400 (Table S5 shows failure counts across all runs and Table S3 shows failure counts restricted to the value of *K_init_* chosen for analysis).

Changing the sparsity of the biclusters in the simulated datasets had a large effect on accuracy. We had expected that on the datasets with only very sparse biclusters, the *Adaptive* algorithms would have best accuracy as they have the strongest sparsity constraints but in fact the *NMF* algorithms performed best on these datasets (Figure S23). We looked at how the recovery of true biclusters was affected by the sparsity of the bicluster and found that most algorithms achieved better recovery scores for denser biclusters (Figure 5).

**Figure 5:**
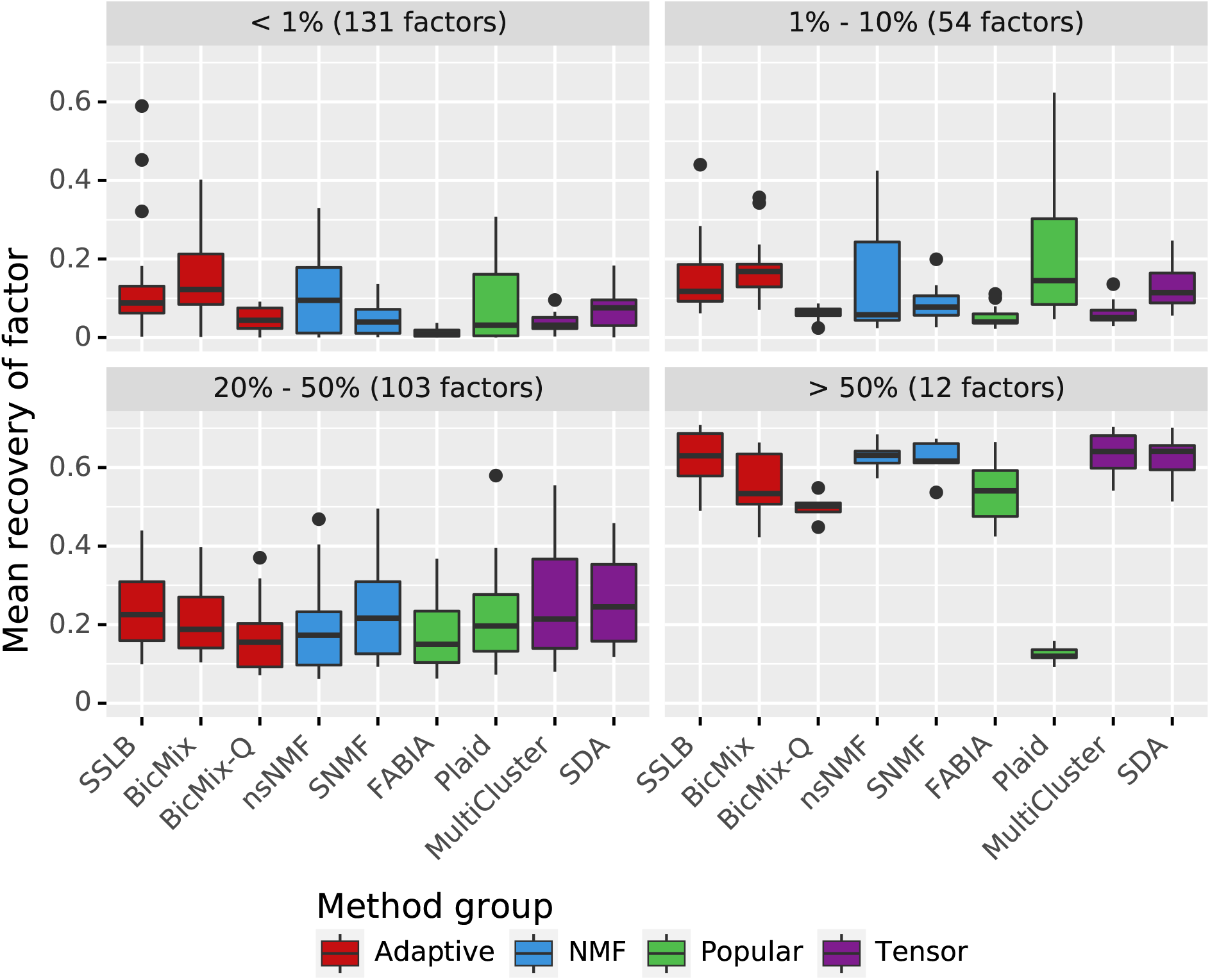
Mean recovery of true biclusters, grouped by size of true biclusters (fraction of total matrix area taken up by true bicluster). We restrict to the datasets base, *sparse, dense, sparse-square, dense-square*. For each true bicluster in these datasets and for each algorithm we find the recovered bicluster achieving maximum Jaccard index with the true bicluster. We call this the recovery score for that true bicluster and that algorithm, which is a measure of how well the algorithm has recovered a particular true bicluster. This plot shows the spread of recovery scores for each algorithm, grouped by the proportion of the total area of the dataset taken up by the true bicluster. Recovery scores are generally better for denser biclusters, though Plaid has notably lower recovery scores for the densest biclusters compared to other algorithms.

The algorithms were in general, however, fairly robust to noise, with little difference in performance between datasets using Negative Binomial noise, Gaussian noise and no noise and only Plaid showing significant decrease in accuracy as noise was increased (Figure S24).

### 3.4 Results on real datasets

The algorithms performed well at finding biclusters corresponding to tissues, with many achieving near perfect performance (Table 4, Figure S25). The algorithms were less effective at finding biclusters corresponding to genotypes, with FABIA and SSLB the top two algorithms (Figure S26). This poor clustering of samples from the same genotype might be due to the fact that the algorithms did not return many biclusters containing samples from multiple tissue types (Figure S27), suggesting that between-tissue differences are dominating over between-genotype differences.

Many algorithms also achieved good clustering of genes, as measured by enrichment of biclusters for Reactome pathways (Figure 6 and Figure S28), with FABIA, SSLB, nsNMF, MultiCluster and Plaid achieving near perfect scores on multiple versions of the dataset. However, this performance should be considered alongside the fact that Plaid returned on average only 4 biclusters and that nsNMF returned factors with high similarity to each other (Figure S14) and thus many of nsNMF’s factors may be enriched for the same small set of Reactome pathways. For example, in one run 145 of the 200 factors recovered by nsNMF were enriched for the ‘Metabolism’ pathway (*q* < 0.05) compared to only 39 of the 188 factors recovered by an SSLB run on the same version of the IMPC dataset.

**Figure 6:**
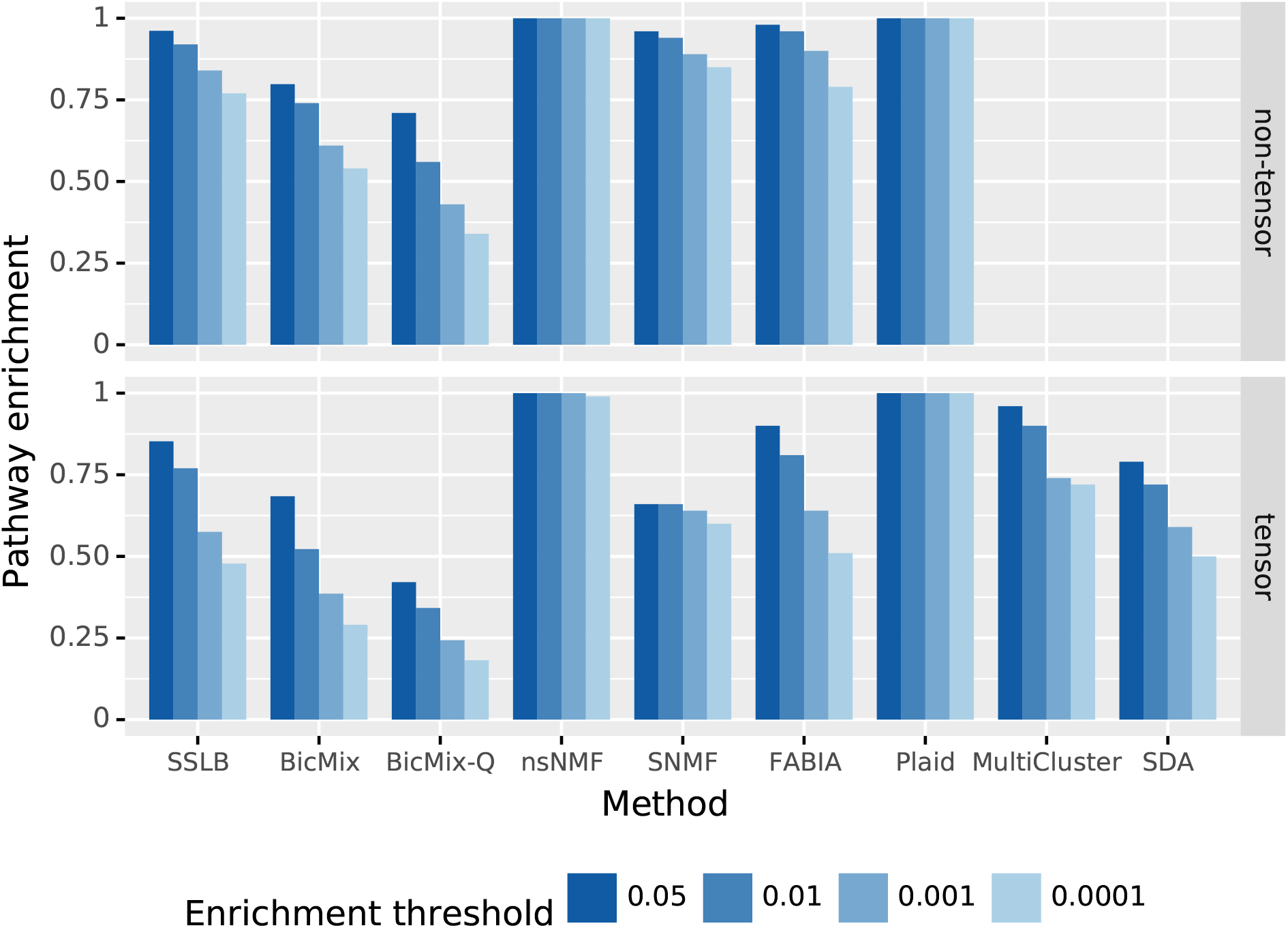
Gene clustering ability, measured by the mean proportion of recovered biclusters which are enriched for at least one pathway. Enrichment is measured using the one-tailed hypergeometric test adjusted for multiple testing using the Benjamini-Yekutieli correction, using a range of thresholds. The median of this measure is shown for each algorithm, split into tensor and non-tensor datasets. Runs that failed (Table S4) are discarded in the analysis.

The unifying test on IMPC data is the biclustering ability, measured as the proportion of knockout genotypes for which the bicluster best recovering the samples is enriched for at least one pathway containing the knocked-out gene (Figure 7). To achieve a high score, an algorithm needs to (1) cluster samples well by genotype, (2) cluster genes well by pathway and (3) return biclusters where there is a link between the samples selected and the genes selected. The *NMF* algorithms, Plaid and SSLB did best according to this metric, achieving enrichment of relevant pathways for approximately 45% of the knockout genotypes in multiple versions of the IMPC dataset. Most algorithms had worse performance when using *K_init_* = 200 (Figure S29), particularly SNMF which failed on 34 of the 40 runs using *K_init_* = 200 (Table S6). Plaid was the only method to fail on a higher percentage of all runs (54 out of 120) but had similar failure rates when using *K_init_* = 50 and *K_init_* = 200.

**Figure 7:**
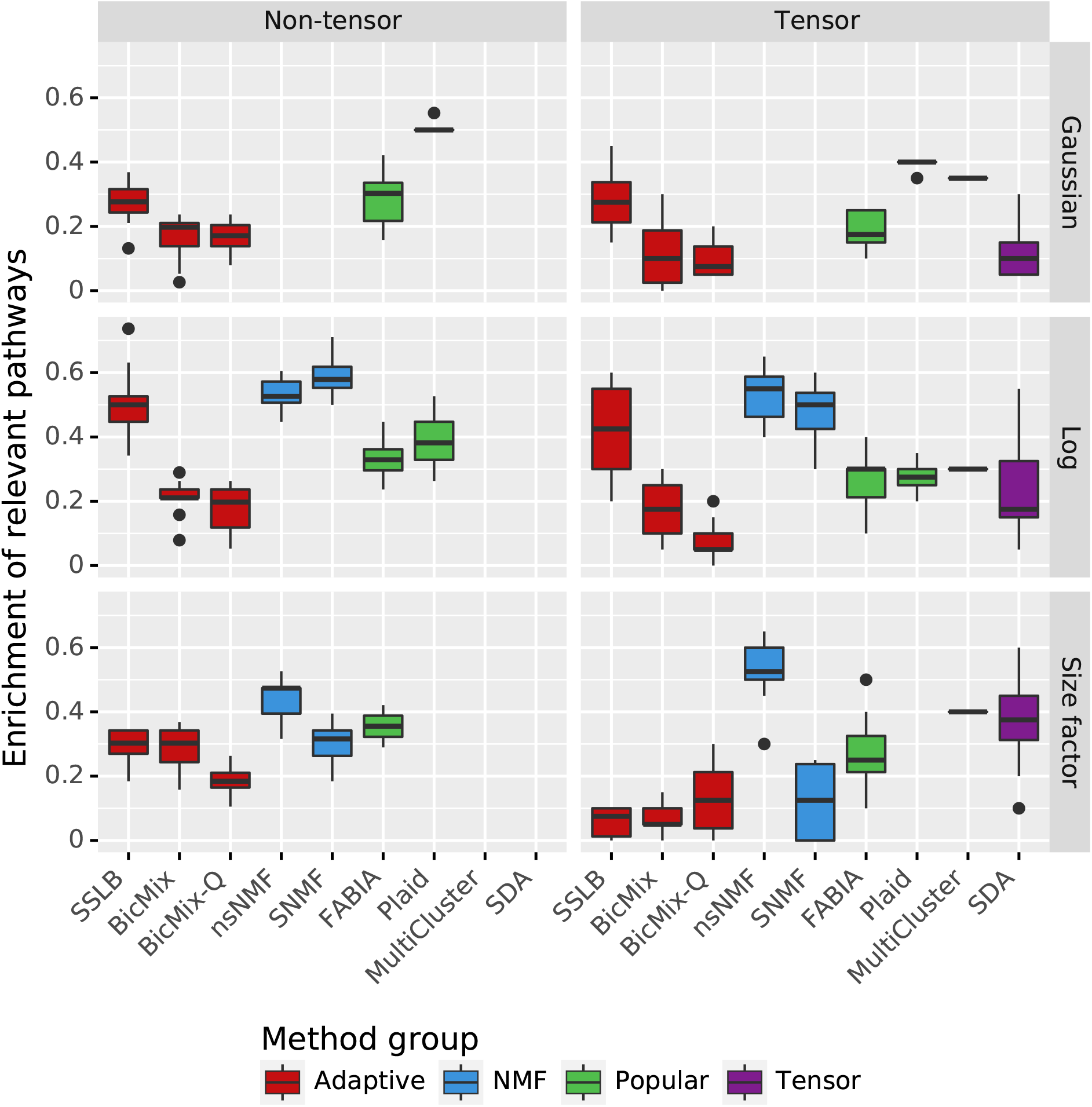
Biclustering ability on IMPC datasets, measured by the mean proportion of knocked-out genes for which the bicluster best matching the samples where the gene was knocked out is enriched for at least one pathway containing the knocked-out gene. Enrichment is measured using the one-tailed hypergeometric test adjusted for multiple testing using the Benjamini-Yekutieli correction, using a threshold for significance of 0.05. Standard thresholding is applied and *K_init_* = 50. Results for *K_init_* = 200 are in S29. Note that *Tensor* algorithms couldn’t be run on the non-tensor datasets, *NMF* algorithms couldn’t run on datasets which used quantile normalisation and Plaid failed to run on the dataset which used DESeq’s size factor normalisation (Table S4).

Strikingly, the reconstruction error (NRE) on the IMPC datasets is much worse (higher) than on the simulated datasets (Figure S30), except for nsNMF. This demonstrates the additional complexity in the real datasets compared to the simulated datasets.

### 3.5 Robustness

With the exception of MultiCluster, the algorithms compared here are stochastic and thus may produce different results each time they are run. If a similar set of biclusters is recovered by repeated runs of an algorithm, this can give confidence that the bicluster decomposition is meaningful. For each algorithm in turn we considered pairs of runs on the same dataset which used the same value of *K_init_* and calculated the similarity between each pair using CE (Clustering Error). As MultiCluster is deterministic, it achieves a perfect score of 1 in this test. Of the remaining algorithms, Plaid and the *NMF* algorithms are the only ones to have a median similarity score between runs of over 0.5 (Figure 8).

**Figure 8:**
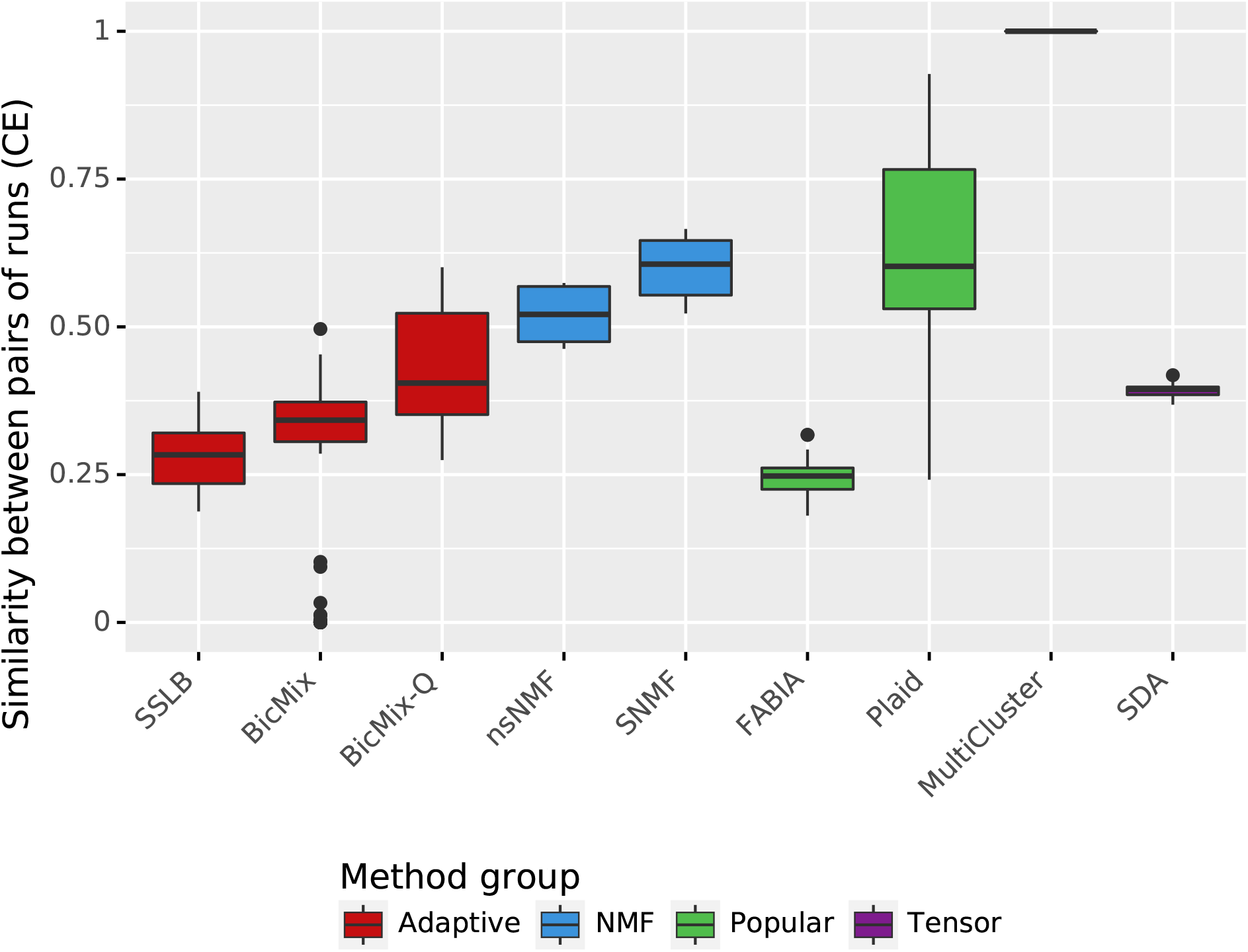
Robustness of biclusters recovered by the algorithms. For each IMPC dataset we calculate similarity scores using clustering error between each pair of runs from a given algorithm, restricted to runs which used the same *K_init_*. This boxplot shows the spread of similarity scores achieved by each algorithm. Higher scores are preferred, as they indicate that the algorithm recovers similar biclusters on each re-run. MultiCluster is deterministic, so achieves maximal score of 1. Plaid and the *NMF* algorithms all achieve relatively high scores.

### 3.6 Computational time

It is important to evalute the time taken for the algorithms to run, and how this scales with the size and complexity of the dataset, as this can restrict the datasets that an algorithm is able to process. The slowest algorithm on the IMPC datasets was SNMF, which took 8 hours to run on the tensor log-transformed dataset, compared to the 7 seconds taken by nsNMF (Table 4). Figure S31 shows runtime with small value of *K_init_*, Figure S32 and Figure S33 show, for simulated datasets and IMPC datasets respectively, that runtime for the *Adaptive* algorithms, SDA, SNMF and FABIA changed drastically with *K_init_*.

## 4 Discussion

On simulated datasets *Adaptive* algorithms had the best overall performance. We investigated the limitations of biclustering algorithms by varying dataset complexity. All algorithms were relatively unaffected by increasing noise in simulated datasets, but performance decreased when dataset size and number of biclusters were increased. Dense biclusters were generally recovered better than sparse biclusters. From a biological perspective, we expect the dense biclusters to correspond to confounding variables such as sex and age and the sparse biclusters to be more biologically interesting, so this behaviour is not ideal.

Like previous studies, we found that algorithms achieved good enrichment of biclusters for gene pathways in real datasets. However, all algorithms struggled to cluster samples from different tissues, highlighting the difficulty of borrowing information across tissue types. We carefully chose a knockout mouse dataset to allow evaluation of biclustering on real datasets, a task which has eluded previous studies, and found that *NMF* algorithms, SSLB and Plaid performed best at recovering biclusters.

In terms of ease of use, *Adaptive* algorithms and Plaid are the only algorithms well suited to use without tuning of *K_init_*, and also didn’t require post-processing. *NMF* algorithms and Plaid had the most robust results, with different runs having on average a similarity of 0.5, as measured by clustering error. Plaid and nsNMF were the fastest, with nsNMF running on the largest IMPC dataset in 7 seconds, compared to the 8 hours taken by the slowest method (SNMF). We found that most algorithms performed well with their default parameters, with few parameter values that showed consistent and significant improvement over the default values during our parameter sweep.

Overall, many algorithms performed better than the *Popular* algorithms which had performed best in previous reviews, showing the need for continued comparison studies as biclustering algorithms develop further. *NMF* algorithms had poor raw output but nsNMF was one of the top-ranking methods after using the sparsity-inducing thresholding procedure we introduce. *Tensor* algorithms did not perform better than other algorithm types, despite both real and simulated datasets having tensor structure. *Adaptive* algorithms performed particularly well on the simulated datasets, and SSLB also had good performance on the real datasets.

## Key points

- We introduce a promising thresholding procedure to enhance sparsity of the returned biclusters, essential for FABIA, SDA, and *NMF* algorithms which otherwise returned only biclusters containing every gene and every sample.
- We introduce the MBR metric for redundancy within a run, and NRE metric for measuring reconstruction error, which can be used even when the true structure of the dataset is not known.
- We have shown the potential for re-purposing of *NMF* algorithms to the task of biclustering. The nsNMF algorithm was orders of magnitude faster than the more complex algorithms, and had good performance, particularly on the real datasets.
- For datasets with unknown structure we recommend SSLB. If a fast algorithm is needed and the number of biclusters is known, or if metrics are available to aid the choice of *K_init_*, then we recommend nsNMF.
- Normalisation method used for real datasets had a large impact on the performance of algorithms. The algorithms performed best on log-transformed data.

