## Supplementary Information for "Comparison of sparse biclustering algorithms for gene expression datasets"

### S1 Algorithms compared

Most algorithms included in this comparison use the same form, differing primarily in whether they explicitly allow sharing of information across cell types and in which approach they use to induce sparsity in the biclusters. We define a matrix $Y \in \mathbb{R}^{n \times p}$ where entry $Y_{ij}$ gives the expression of gene $j$ in sample $i$. This can either be raw read counts from an RNA-seq experiment, or normalised counts which have been adjusted for sample-specific effects such as library size, or gene-specific effects such as mean expression level. The typical approach to biclustering is to factorise this matrix as a product of two matrices $X \in \mathbb{R}^{n \times K}$, which we call the sample loadings matrix, and $B \in \mathbb{R}^{p \times K}$, which we call the gene loadings matrix, with error matrix $\varepsilon$:

$$Y = XB^T + \varepsilon \tag{1}$$

#### S1.1 Plaid

The general Plaid model is:

$$Y_{ij} = \mu_0 + \sum_{k=1}^{K} (\mu_k + \alpha_{jk} + \beta_{ik}) \rho_{jk} \kappa_{ik} + \varepsilon_{ij} \tag{2}$$

The gene expression matrix is written as a sum of $K$ layers, each of which is a sum of a layer effect $\mu_k$, gene-specific effects $\alpha_{jk}$ for each gene and sample-specific effects $\beta_{ik}$ for each sample. The membership of genes and samples to each layer is given by binary variables $\rho_{jk}$ and $\kappa_{ik}$ respectively. Finally, there is a background layer with constant value $\mu_0$. The error for each entry is given by $\varepsilon_{ij}$. Layers are added one-at-a-time and parameters are chosen by alternating ordinary least squares.

#### S1.2 FABIA

Factor Analysis for Bicluster Acquisition (FABIA) [1] is a sparse factor analysis model using the model given in Equation 1. FABIA induces sparsity by using a Laplacian prior on all entries of $X$ and $B$. The same prior is used for all entries in the matrix.

In the parameter sweep conducted before full analysis, we found that using higher sparsity by setting spz = 1.5 greatly improved FABIA's performance.

#### S1.3 BicMix

BicMix [2, 3] uses the model given in Equation 1 and allows each bicluster to be either *'sparse'* or *'dense'*. On all 'sparse' biclusters, BicMix encourages sparsity through three layers of shrinkage, each of which uses a three parameter beta (TPB) prior [4].

One parameter that drastically affected performance was whether the data was quantile-normalised before BicMix ran. Since the effect was substantial but inconsistent, we ran BicMix twice on each dataset: once with quantile-normalisation (BicMix-Q) and once without (BicMix).

#### S1.4 SSLB

Spike-and-Slab Lasso Biclustering (SSLB) [5] is another Bayesian biclustering algorithm using the model given in Equation 1. SSLB uses the Spike-and-Slab Lasso prior [6] on both the factor matrix X and the loadings matrix B. The Spike-and-Slab Lasso prior is a mixture of two Laplacians, one of which has weak regularisation (the *slab*) and one has strong regularisation (the *spike*). When fitting a model, stronger regularisation will result in a sparser result but may come at the expense of accuracy. Using this prior allows much stronger regularisation on the coefficients that are near-zero (those in the spike) in order to achieve sparsity, whilst having weaker regularisation on the larger coefficients (those in the slab) to avoid loss of accuracy. Moran et al. claim that their algorithm produces sparser matrices than FABIA and BicMix.

Another advantage of the SSLB prior compared to FABIA and BicMix is that it allows different levels of sparsity for each bicluster. FABIA uses the same prior for all biclusters, and BicMix only allows for two different levels of sparsity ('sparse' or 'dense'). SSLB allows each bicluster to have a different sparsity parameter.

#### S1.5 SDA

In contrast to the matrix algorithms seen so far, Sparse Decomposition of Arrays (SDA) [7] rearranges the gene expression matrix to be a 3D tensor with dimensions $m \times p \times t$ where the total number of samples $n = mt$ is the product of the number of individuals $m$ and number of tissues $t$. It decomposes the gene expression matrix $Y$ as a tensor product of three matrices $A, B, Z$ corresponding to individual, gene and tissue components respectively:

$$Y = \sum_k a_k \otimes b_k \otimes z_k + \varepsilon \tag{3}$$

This enables information to be shared between different tissues, which the other algorithms are unable to do. This decomposition can be written as a sum of contributions from each bicluster:

$$Y_{ijl} = \sum_k A_{ik} B_{jk} Z_{lk} + \varepsilon_{ijl} \tag{4}$$

SDA encourages sparsity in the gene components by using a prior similar to that of SSLB. It uses the Gaussian Spike-and-Slab prior described in [8], which consists of a point-mass at 0 (the *spike*) and a Gaussian distribution (the *slab*). It does not encourage sparsity in either sample components or tissue components.

The implementation of SDA that we use provides an alternative model for when $t = 1$, but we have chosen not to include it in this comparison as the tensor factorisation seems the most promising.

#### S1.6 SNMF

Sparse non-negative matrix factorisation (SNMF) [9] factorises the gene expression matrix $Y$ as a product of two non-negative matrices $X$ and $B$. Non-negativity makes the biclusters easier to interpret, and naturally encourages sparsity. SNMF uses an $L_1$ penalty on the elements of the gene loadings matrix $B$ to further encourage sparsity of this matrix. The paper also includes a algorithm that uses an $L_1$ penalty on the sample loadings matrix instead, but we chose to use the version with sparsity on the gene loadings matrix as one of the main aims is to find interesting groups of genes, and these are easier to interpret if they are sparser.

#### S1.7 nsNMF

Like SNMF, the non-smooth non-negative matrix factorisation (nsNMF) model [10] factorises the gene expression matrix as a product of non-negative matrices:

$$Y = XSB \tag{5}$$

The key difference is the inclusion of a third matrix, called the *smoothing matrix*:

$$\mathbf{S} = (1-\theta)\mathbf{I} + \frac{\theta}{K}\mathbf{1}\mathbf{1}^T \tag{6}$$

When $\theta = 0$, we have $S = I$ which makes nsNMF equivalent to normal non-negative matrix factorisation. However, for larger $\theta$, the matrix $S$ helps to increase sparsity without increasing error. Without it, increasing sparsity of samples loading matrix $X$ would encourage the genes loading matrix $B$ to be denser in order for the product $XB$ to be close to the original matrix $Y$. Thus if we want sparsity in both $X$ and $B$, we will suffer an increase in error. With nsNMF, $Y = (XS)B$ so in order to allow $B$ to be sparse, we must make sure that $XS$ is not too sparse. When a matrix is multiplied by $S$, it becomes less sparse. Thus we can encourage $X$ to be sparse, and $XS$ will not be that sparse, meaning that $B$ will not be forced to be denser. Thus, with nsNMF we can apply sparsity constraints to $B$ as well as to $X$ without a large increase in the error.

#### S1.8 MultiCluster

MultiCluster [11], like SDA, rearranges the gene expression matrix into a 3D tensor and decomposes it as:

$$Y = \sum_k a_k \otimes b_k \otimes z_k + \varepsilon \tag{7}$$

To aid interpretability and encourage sparsity, the tissue components $z_k$ are constrained to be non-negative. MultiCluster uses a deterministic algorithm to find optimal components $a_k, b_k, z_k$ for each $k$ in turn.

Table S1: Datasets used in the parameter sweep.

| Name | Individuals | Tissues | Genes | Bicluster sizes | K | Noise |
|---|---|---|---|---|---|---|
| negbin_K20 | 10 | 10 | 1000 | mixed | 20 | NegBin |
| gaussian_K50 | 10 | 10 | 1000 | small | 50 | Gaussian |
| no_noise_K10 | 10 | 10 | 1000 | large | 10 | No noise |

### S2 Parameter sweep

Complex algorithms such as those included in this review can be very sensitive to choice of parameters. This has two consequences for this review: firstly we need to decide how to set the parameters during the review, and secondly we should consider the robustness of algorithms to parameter choice in itself as a criterion by which to judge the algorithms.

Many previous reviews of biclustering algorithms have used a single set of parameters throughout the study [12, 13]. By taking this approach, the comparison study naturally favours the algorithms which have good default parameters, or which do not vary much depending on the parameter choice. Another approach is to try many different combinations of parameters for each algorithm on each dataset. In a review of module detection algorithms, Saelens et al. performed a grid search for each algorithm on each dataset to find an optimal set of parameters [14] for each dataset. This approach of finding optimal parameter sets can be computationally expensive and also does not extend well to real datasets, where calculating a score for results is more difficult, as the true structure of the dataset is unknown.

We conducted a investigation of the effect of changing key parameters for each algorithm, looking for parameter values that consistently improved performance over the default parameters on three distinct simulated datasets. Overall we found that most parameters had little effect on performance as measured by clustering error (CE) and for most parameters, there was either no clear optimal parameter, or the default parameter was clearly optimal. The results are summarised in Table S2.

We simulated three very differently structured datasets (Table S1), and ran each algorithm with five random seeds using $K_{init} = 20$ and five seeds using $K_{init} = 50$ for each set of parameter values on each dataset. For each parameter (except $K$) we chose approximately five different values, including the default value. We then ran the algorithm changing each parameter in turn, keeping the other parameters at their default value, running 5 times with $K = 20$ and 5 times with $K = 50$. Below we discuss for each algorithm how we chose the parameters to change, and show the effect of changing each parameter whilst keeping the other parameters at their default value.

#### S2.1 BicMix

##### S2.1.1 Prior

BicMix uses a Three Parameter Beta prior to induce sparsity in the factor values. There are two key parameters that control this prior, with default values $a = b = 0.5$. This choice of $a, b$ is called the Horseshoe Prior. [4] discusses hyper-parameters for the Three Parameter Beta distribution in sparse models, and note that it is necessary to have $a < 1$ in order to

Table S2: Effect of parameters on Clustering Error (CE). For each parameter we consider if changing it had an effect on performance and if so whether the effect was consistent and substantial. Finally we note the action we decided to take, leaving this column blank if the default value is used.

| Method | Parameter | Effect? | Consistent? | Substantial? | Action |
|---|---|---|---|---|---|
| BicMix | a | **Yes** | No | No | |
| | b | **Yes** | No | No | |
| | qnorm | **Yes** | No | **Yes** | Run with both |
| FABIA | alpha | No | | | |
| | spz | **Yes** | **Yes** | **Yes** | Use spz=1.5 |
| | spl | No | | | |
| | eps | **Yes** | **Yes** | **Yes** | None, fixed with spz=1.5 |
| | rescale_l | **Yes** (failure) | | | |
| Plaid | row_release | **Yes** | No | **Yes** | |
| | col_release | **Yes** | No | No | |
| SDA | num_dense | No | | | |
| | step_size | **Yes** | **Yes** | No | |
| | conv_crit | No | | | |
| nsNMF | $\theta$ | **Yes** | **Yes** | No | |
| SNMF | $\beta$ | **Yes** | No | No | |
| SSLB | d | No | | | |
| | IBP | No | | | |
| | alpha | No | | | |
| | a | No | | | |
| | b | No | | | |

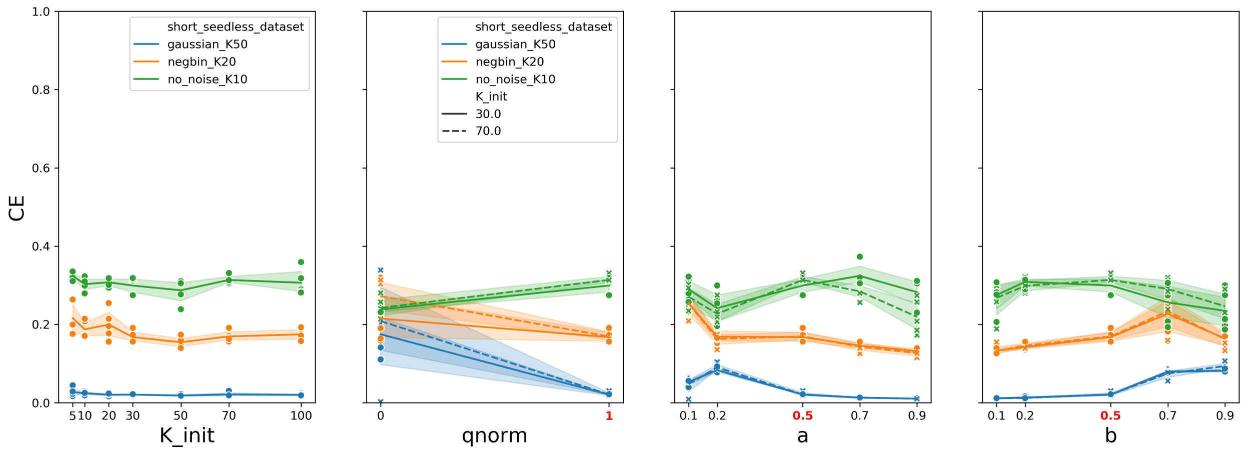


Figure S1: Parameter sweep for BicMix. Each plot shows the effect on the clustering error measure (CE) when a single parameter is varied, while the other parameters are kept at their default values. The default value for a parameter is highlighted in red on the x-axis. Higher clustering error values are better.

achieve sparse solutions, and suggest using $b < 1$ too. We used this to guide our choice of values to try for $a, b$.

The performance is affected by these parameters, but the optimal parameter choice isn't the same for all datasets (Figure S1).

##### S2.1.2 Pre-processing

The other important parameter is *qnorm*, which determines whether or not a Gaussian rank transformation is applied to each gene in the data matrix before BicMix runs.

Gaussian rank transformation transforms each variable (gene) so that the resulting distribution is exactly Gaussian. The values from the original variable are ranked with ties broken at random, and the value at position $k$ in the list of $n$ values will be mapped to the $\frac{k}{n+1}$ quantile of the standard Gaussian $N(0, 1)$. A key consequence of this is that the many 0s will get mapped to *different* values, so this introduces noise to the data.

Changing *qnorm* has a dramatic effect on performance (Figure S1). Most notably, the performance on the dataset *gaussian_K50* is near 0 when the data are quantile-normalised.

Due to the large changes in performance with quantile-normalisation, we ran two versions of BicMix: one with quantile-normalisation and one without. To check that the default prior parameters are sufficient when quantile-normalisation is not used, we performed another parameter sweep with $qnorm = 0$ and found that there is no parameter choice that shows consistent improvement over the default choice $a = b = 0.5$ (Figure S2).

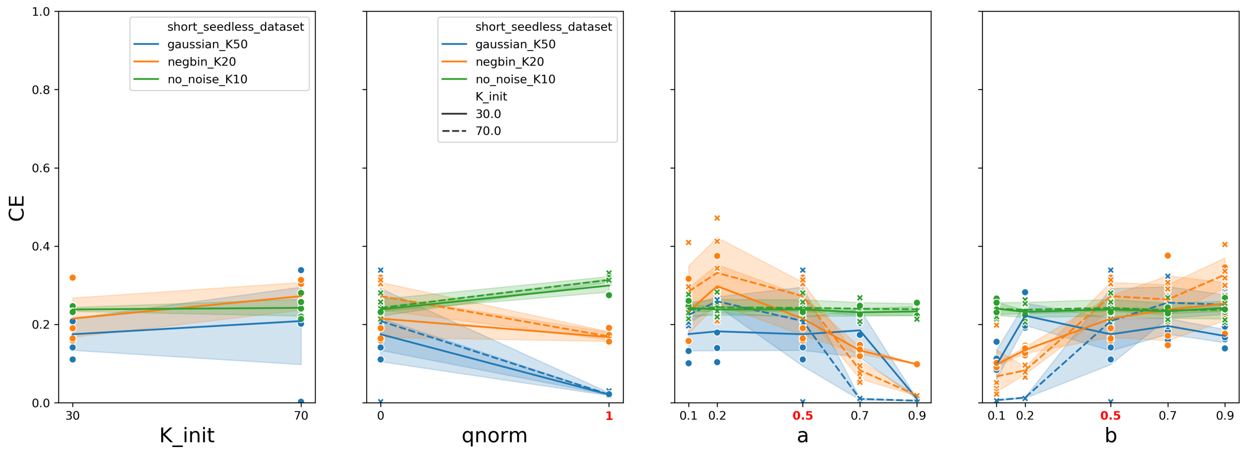


Figure S2: Parameter sweep for BicMix without qnorm. Each plot shows the effect on the clustering error measure (CE) when a single parameter is varied, while the other parameters are kept at their default values (with the exception of qnorm, which is kept at 0 for the parameter sweeps for all other parameters). The default value for a parameter is highlighted in red on the x-axis. Higher clustering error values are better.

#### S2.2 FABIA

##### S2.2.1 Sparsity

FABIA controls the sparsity of the factor matrix $X$ using *spz*. The sparsity of the loadings matrix $B$ is controlled using $\alpha$ and *spl*. For all these parameters, higher values increase sparsity. The parameter *eps* determines a threshold under which entries in either $X$ or $B$ will be treated as 0, so higher values increase sparsity of both $X$ and $B$. The *pyfabia* implementation gives ranges for each value, which we used to guide our choice of values.

The two that seem to have the biggest effect are *eps* and *spz* (Figure S3). The best performance was achieved with $spz = 1.5$ for all datasets. Since the change is substantial and consistent, we chose to use this value of the parameter for the rest of the study. We investigated the effect of changing *eps* when $spz = 1.5$ and found that it still had an effect, but it was not as great as when $spz = 0.5$ (Figure S4).

##### S2.2.2 Rescaling during fitting

If *rescale_l* is *True* then FABIA rescales each $B_k$ to have unit variance each iteration. This is not the default behaviour, but the *pyfabia* documentation claims this can help convergence, but we found that many runs using this rescaling actually failed to return a result.

##### S2.2.3 Bicluster membership

After fitting the model, both the *pyfabia* and R FABIA implementations provide functions to determine which rows and columns belong to each bicluster. Values under the threshold are set to 0, essentially removing this element from the bicluster. The threshold is *thresZ* for $X$ and *thresL* for $B$. These are the only two parameters varied in [14], a review of

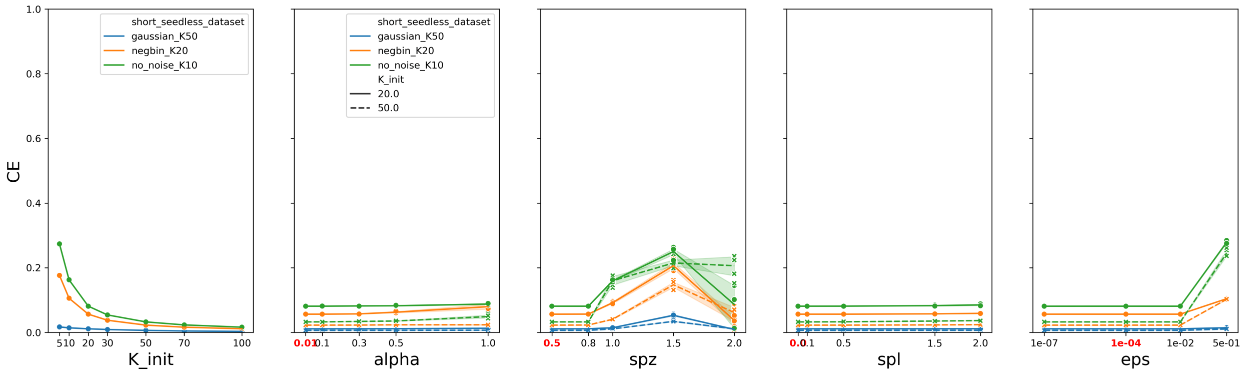


Figure S3: Parameter sweep for FABIA. Each plot shows the effect on the clustering error measure (CE) when a single parameter is varied, while the other parameters are kept at their default values. The default value for a parameter is highlighted in red on the x-axis. Higher clustering error values are better.

module detection algorithms. This thresholding is a post-processing step which we apply to all algorithms, so we ignored this parameter.

#### S2.3 Plaid

In a Bacherlor's thesis focusing on the Plaid algorithm, Pfundstein notes that the two key paramters to tune are *row_release* and *col_release*, which determine when a row or column is removed from a bicluster [15]. These are also the only two parameters altered in the module detection review [14]. A small recommended range of $[0.5, 0.7]$ is given in the documentation of the *biclust* package that contains the Plaid implementation we used.

Changing these parameters has some effect, but the effect differs for each dataset (Figure S5). Since we are primarily using Plaid as a benchmark to allow results in this study to be related to other reviews of biclustering algorithms, and since these reviews mostly used the default parameters, we did the same.

#### S2.4 MultiCluster

The implementation of MultiCluster that we use doesn't have any parameters that can be changed.

#### S2.5 SDA

##### S2.5.1 Prior

The parameters determining the priors are set up so that the priors are uninformative, so we didn't alter them.

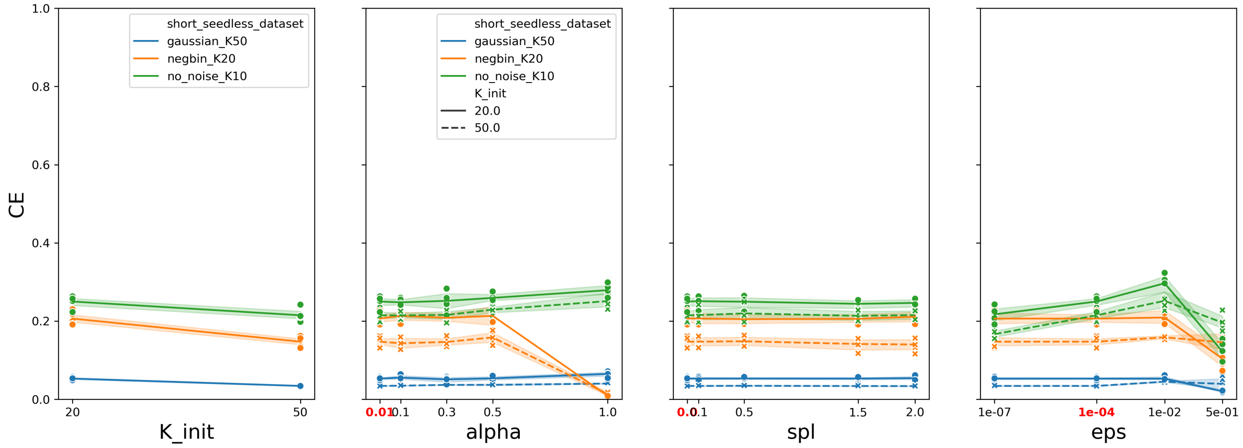


Figure S4: Parameter sweep for FABIA with $spz = 1.5$. Each plot shows the effect on the clustering error measure (CE) when a single parameter is varied, while the other parameters are kept at their default values. The default value for a parameter is highlighted in red on the x-axis. Higher clustering error values are better.

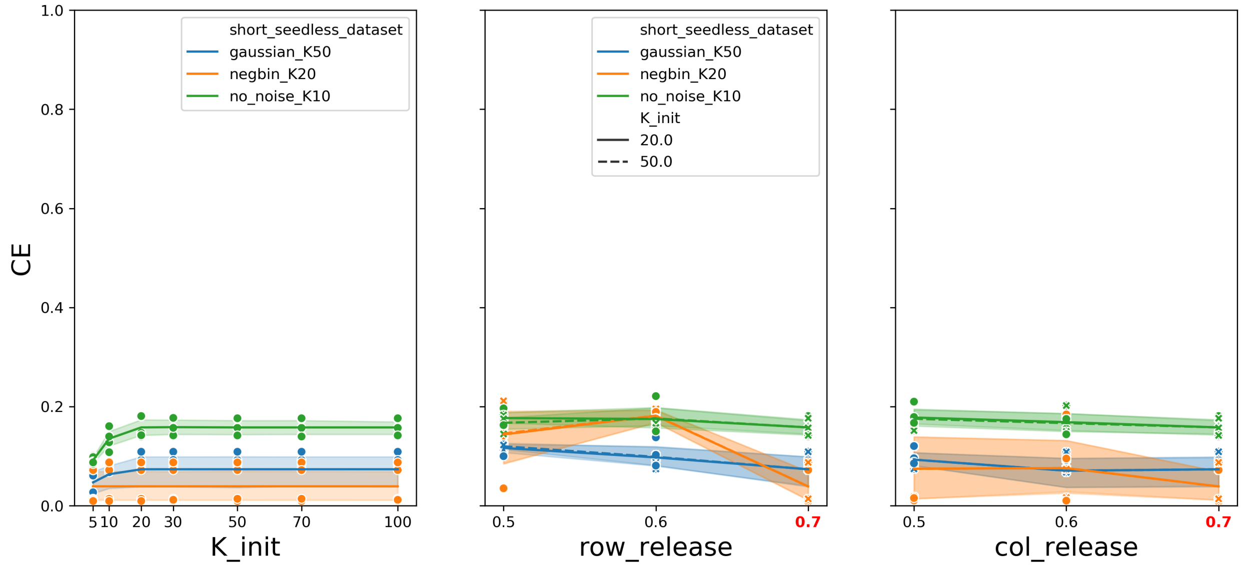


Figure S5: Parameter sweep for Plaid. Each plot shows the effect on the clustering error measure (CE) when a single parameter is varied, while the other parameters are kept at their default values. The default value for a parameter is highlighted in red on the x-axis. Higher clustering error values are better.

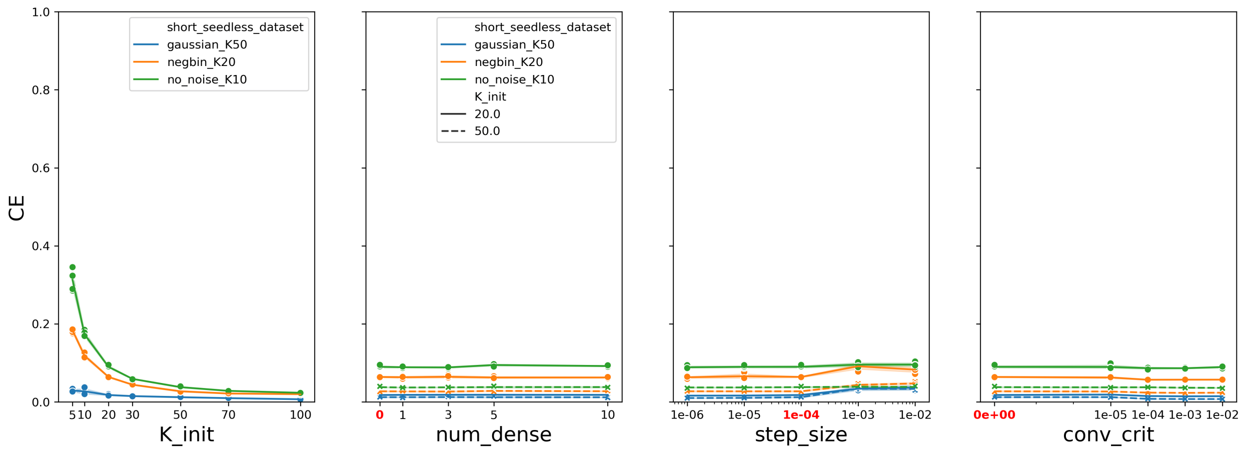


Figure S6: Parameter sweep for SDA. Each plot shows the effect on the clustering error measure (CE) when a single parameter is varied, while the other parameters are kept at their default values. The default value for a parameter is highlighted in red on the x-axis. Higher clustering error values are better.

##### S2.5.2 Convergence

SDA is one of the slower algorithms. We suspected this was partly due to its strict convergence criteria. The parameter *conv_crit* determines the largest allowed change in free energy at convergence, thus larger values make it easier for the algorithm to converge. The default value is 0. In Figure S6 we see there is no difference when *conv_crit* is $1 \times 10^{-5}$, but marginal differences for larger values. Using $1 \times 10^{-5}$ seems to have little effect on performance, but decreases the median running time by 50% (Figure S7). We recommend users consider changing from the default value of *conv_crit* in this way to save computational time. We chose not to make the change in this study as it is valuable for us to show performance using the default values, since this is what most users will do.

#### S2.6 nsNMF

The only crucial parameter for nsNMF is $\theta$ which controls sparsity. The higher $\theta$, the sparser the model. Surprisingly, we found that changing $\theta$ had little impact (Figure S8). The dataset that suffered worst performance from using a high value of $\theta$ (thus a sparser model) was the *gaussian_K50* dataset which had the smallest biclusters.

#### S2.7 SNMF

As with nsNMF, the important parameter is $\beta$, which controls sparsity. Larger $\beta$ makes the model sparser, but we found that changing the sparsity had little impact on the performance (Figure S9).

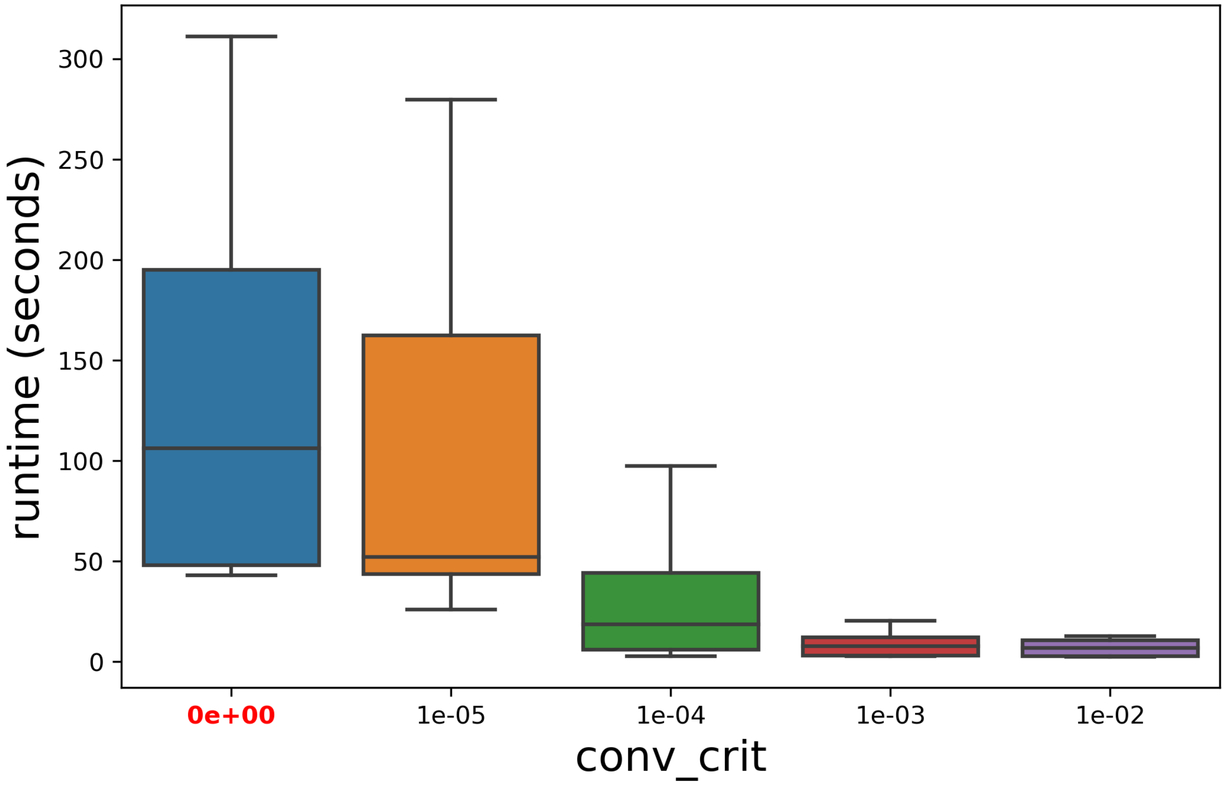


Figure S7: Running time of SDA with different *conv_crit* values.

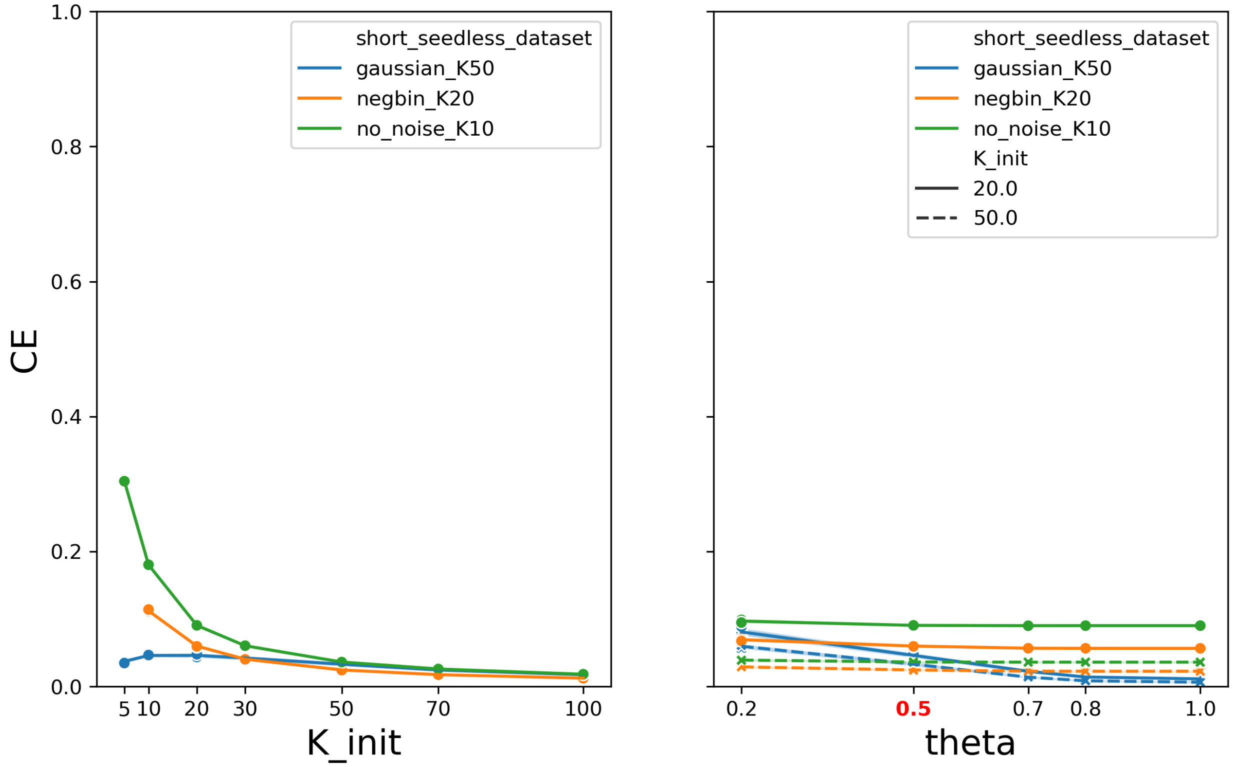


Figure S8: Parameter sweep for nsNMF. Each plot shows the effect on the clustering error measure (CE) when a single parameter is varied, while the other parameters are kept at their default values. The default value for a parameter is highlighted in red on the x-axis. Higher clustering error values are better.

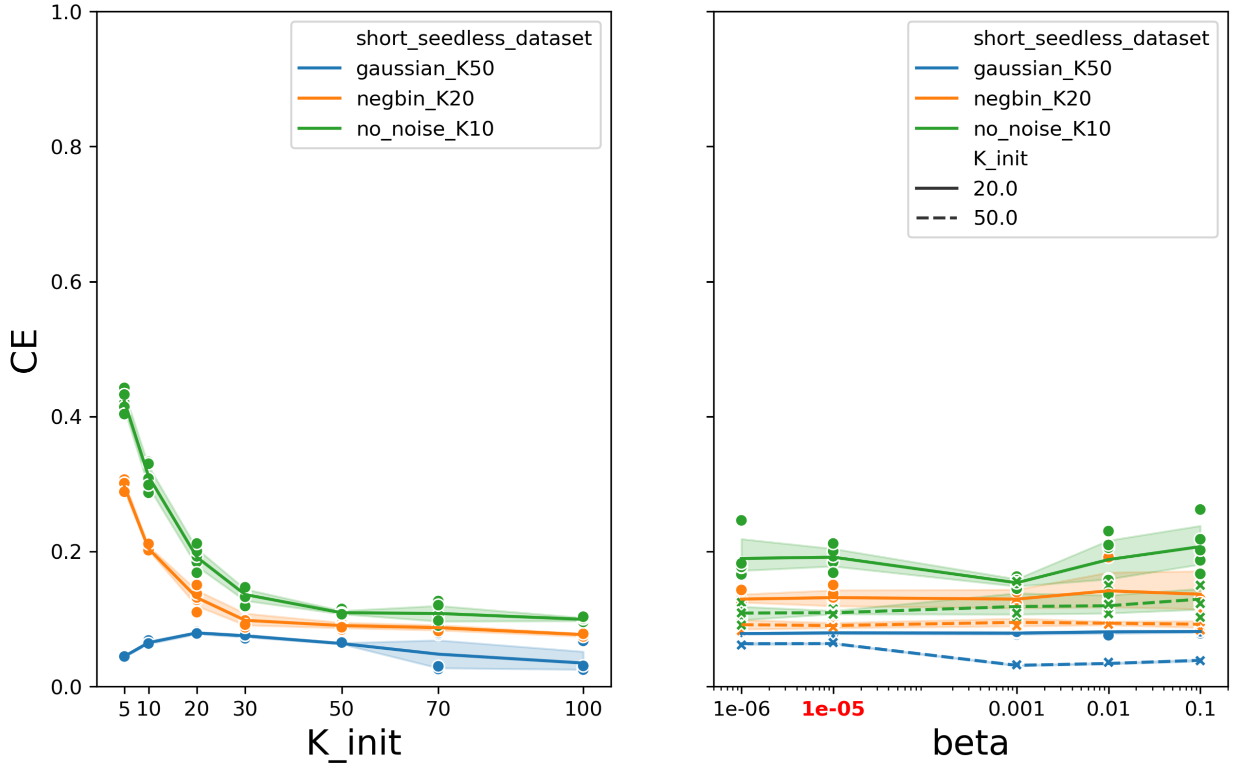


Figure S9: Parameter sweep for SNMF. Each plot shows the effect on the clustering error measure (CE) when a single parameter is varied, while the other parameters are kept at their default values. The default value for a parameter is highlighted in red on the x-axis. Higher clustering error values are better.

#### S2.8 SSLB

The parameter likely to have the biggest effect is the choice of prior for the values in $B$ and $X$.

##### S2.8.1 Prior for $B$

The prior on $B$ is a Beta-Binomial, with parameters $a, b$. Changing these seems to have little effect on the performance of SSLB (Figure S10).

##### S2.8.2 Prior for $X$

There are three different priors that can be used for $X$. The prior choice is controlled by the parameters *IBP* and $d$. If IBP = 0 then the simple Beta-Binomial prior is used; if IBP = 1 and $d = 0$ then a normal IBP is used; if IBP = 1 and $d \neq 0$ then a Pitman-Yor extension of the IBP is used.

The simplest is the same Beta-Binomial used for $B$, which has parameters $\tilde{a}, \tilde{b}$. We chose to experiment with using this Beta-Binomial prior, but, since it is not the default and generally performed worse than the other priors in the original paper [5], we decided not to investigate optimal parameters for this prior. Figure S10 shows that using the Beta-Binomial prior (i.e. IBP = 0) slightly degrades the perforance.

The prior used by SSLB by default is the IBP prior, which uses the parameter $\alpha$. This has default value $\alpha = \frac{1}{N}$, i.e. it is dependent on the number of samples. All the datasets shown in Figure S10 have the same number of samples: $N \times T = 10 \times 10 = 100$, so the default value was 1/100 for all datasets.

Finally, the Pitman-Yor extension to the IBP prior uses both $\alpha$ and $d$. We only investigated the effect of changing $d$. $d = 0$ gives the standard IBP prior, and we did not find a significant difference when the Pitman-Yor extension was used, for any values of $d$ (Figure S10).

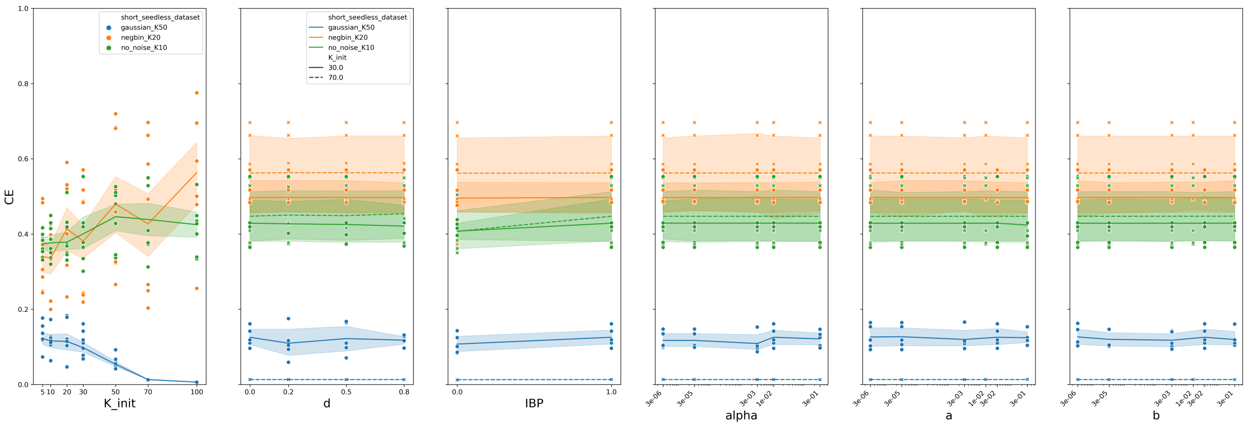


Figure S10: Parameter sweep for SSLB. Each plot shows the effect on the clustering error measure (CE) when a single parameter is varied, while the other parameters are kept at their default values. Default values were: $d = 0, \text{IBP} = 1$ (i.e. using the standard IBP prior), $\alpha = \frac{1}{N} = \frac{1}{100}$, $a = b = \frac{1}{K}$. Higher clustering error values are better.

### S3 Evaluation Metrics

#### S3.1 Accuracy on simulated datasets

Common choices for metric to compare biclusterings, and thus to evaluate biclustering algorithms on simulated datasets, are the *consensus score* [1, 5] and the *recovery* and *relevance* scores [12, 5, 2, 3]. Horta and Campello defined eight desirable properties of similarity measures for comparing biclusterings, including for example whether the measure penalises biclusterings with duplicated biclusters [16]. The consensus score satisfies only four of these desirable properties, and the recovery and relevance scores satisfy only one.

We chose to use the *clustering error* (CE) metric [17, 16] to evaluate biclustering accuracy, which was shown to satisfy all but one of the desirable properties defined by Horta and Campello, which is a great improvement on the consensus score and recovery and relevance scores commonly used to evaluate biclustering similarity.

##### S3.1.1 Metrics comparing two biclusterings

To be able to define metrics comparing two biclusterings, we first write biclusters as sets of elements. Suppose we have two factorisations $Y_{ij} = \sum_k X_{ik} B_{jk}$ and $\hat{Y}_{ij} = \sum_l \hat{X}_{il} \hat{B}_{jl}$. We write each bicluster as a set of the matrix elements that it contains: $A_k = \{(i,j) : X_{ik} \neq 0, B_{jk} \neq 0\}$ and $\hat{A}_l = \left\{(i,j) : \hat{X}_{il} \neq 0, \hat{B}_{jl} \neq 0\right\}$.

The Jaccard index [18] measures how closely two sets match, comparing their intersection to their union. Using the description of biclusters as sets, we can use the Jaccard index to measure how closely two biclusters match:

$$J(A_k, B_l) = \frac{\left|A_k \cap \hat{B}_l\right|}{\left|A_k \cap \hat{B}_l\right|} \tag{8}$$

The recovery and relevance scores [12] are defined as:

$$\begin{aligned} \text{Relevance} &:= \frac{1}{K} \sum_{k=1}^{K} \max_l \; J\left(A_k, \hat{A}_l\right), \\ \text{Recovery} &:= \frac{1}{L} \sum_{l=1}^{L} \max_k \; J\left(A_k, \hat{A}_l\right) \end{aligned} \tag{9}$$

The consensus score also makes use of the Jaccard index, but first calculates the best way to pair biclusters from each set. We can write a pairing of biclusters as $\left\{\left(A_{k_i}, \hat{A}_{l_i}\right)\right\}_{i=1}^{\min\{K,L\}}$ where each element of this set gives a pair: a bicluster $A_{k_i}$ from $A$ and the bicluster $\hat{A}_{l_i}$ from $\hat{A}$ that it is paired with. Note that if one biclustering contains more biclusters than the other, there will be some unpaired biclusters. We seek the pairing which maximises the following sum, the sum of the Jaccard indices of pairs divided by the number of biclusters in the larger biclustering:

$$\text{Consensus} := \frac{1}{\max K, L} \sum_{i=1}^{\min K,L} \left| A_{k_i} \cap \hat{A}_{l_i} \right| \tag{10}$$

In practice the optimal pairing is found using the Hungarian algorithm for the assignment problem [19].

##### S3.1.2 Clustering error

The clustering error (CE) is defined as:

$$\text{CE}(A, \hat{A}) := \frac{d_{\max}}{|U|} \tag{11}$$

where $d_{max}$ measures how much the biclusterings intersect and $|U|$ measures the total space collectively covered by the biclusterings, taking overlaps into account. Despite its name, CE is a measure of similarity between biclusters rather than dissimilarity.

We often use this measure to compare a biclustering $\hat{A} = \left\{\hat{A}_1, \dots \hat{A}_L\right\}$ recovered by a algorithm to a biclustering $A = \{A_1, \dots, A_K\}$ which we treat as the *true* biclustering. However, this measure is symmetric and can also be used when there is no 'truth', such as to compare two biclusterings returned by different runs of the same algorithm.

In a similar way to the consensus score, we use the Hungarian algorithm to find a pairing of biclusters from the two sets to maximise a sum, this time the sum of the intersections of pairs:

$$d := \sum_{i=1}^{\min K,L} \left| A_{k_i} \cap \hat{A}_{l_i} \right| \tag{12}$$

The maximum value this sum attains is called $d_{\max}$ and measures how much the biclusterings intersect.

The *size counting overlapping* of a union of two sets, called $|U|$, is

$$|U| := \sum_{i,j} \max \left\{ \mathrm{N}_{i,j}, \hat{\mathrm{N}}_{i,j} \right\} \tag{13}$$

where $N_{i,j} := |\{k : (i,j) \in A_k\}|$ gives the number of times that the matrix element $(i,j)$ is included in biclusters in $A$ and $\hat{N}_{i,j} := \left|\left\{l : (i,j) \in \hat{A}_l\right\}\right|$ is the same for $\hat{A}$.

#### S3.2 Normalised reconstruction error

Most metrics used by previous reviews, including the *clustering error* metric that we intend to use, can only be used on simulated datasets. We introduce two metrics that can be used even when nothing is known about the structure of the dataset. The first uses the error matrix $\varepsilon = Y - \hat{X}\hat{B}^T = Y - \hat{Y}$ to see how similar the recovered factorisation is to the original matrix.

Given original matrix $Y$, and factorisation $\hat{Y} = \hat{X}\hat{B}^T$ returned by the algorithm, we define the normalised reconstruction error (NRE) as:

$$\text{NRE}(Y, \hat{Y}) := \frac{\|Y - \hat{Y}\|_F}{\|Y\|_F + \|\hat{Y}\|_F} \tag{14}$$

where $\|A\|_F$ denotes the Frobenius norm $\sqrt{\sum_{ij} A_{ij}^2}$.

A score of 0 indicates perfect reconstruction. The maximum score is 1, indicating a large error relative to the true matrix $Y$ and the recovered matrix $\hat{Y}$. One big advantage of this metric is that it can be used on real datasets too, since all that is needed is the original matrix $Y$.

#### S3.3 Mean Bicluster Redundancy (MBR)

The second metric we introduce that can be used without knowledge of the structure of the dataset is the Mean Biclustering Redundancy (MBR), which measures how similar the biclusters returned in a single run are to each other. When running biclustering algorithms on large datasets, it can be difficult to interpret the results if the algorithms return many copies of the same bicluster. The perfect score of 0 indicates that the biclusters do not overlap at all, and the worst score of 1 indicates that all biclusters are identical.

We first construct a matrix $J$ using the Jaccard index between each pair of biclusters:

$$J_{kl} := \frac{|A_k \cap A_l|}{|A_k \cup A_l|} \tag{15}$$

and then take the mean of the off-diagonal entries:

$$\text{MBR}(A) := \frac{2}{K(K-1)} \sum_{k=1}^{K-1} \sum_{l=k+1}^{K} J_{kl} \tag{16}$$

#### S3.4 Sample clustering in real datasets

The samples in the IMPC dataset can be clustered in two main ways: by tissue or by genotype. We refer to these as *sample traits*. We chose to measure how well a bicluster matches a given sample trait using the $F_1$ score, which balances reward for containing *only* elements with the sample trait (*precision*) and reward for containing *all* the elements with that sample trait (*recall*). It is defined as:

$$F_1 \text{ score} = 2 \times \frac{\text{Precision} \times \text{Recall}}{\text{Precision} + \text{Recall}} \tag{17}$$

where precision and recall are defined in terms of the set $S$ of samples with the trait and set $F$ of samples contained in the bicluster:

$$\begin{aligned} \text{Precision} &= \frac{|S \cap F|}{|F|} \\ \text{Recall} &= \frac{|S \cap F|}{|S|} \end{aligned} \tag{18}$$

For each trait we find the best $F_1$ score across all biclusters, and then take the mean of these maximum $F_1$ scores across sample traits. We call the mean of this maximum across all traits the sample clustering ability, the mean across only tissue traits we call the tissue clustering ability and the mean across only genotype traits we call the genotype clustering ability.

#### S3.5 Gene clustering in real datasets

The typical approach to evaluate gene clustering in real datasets is to look at what proportion of biclusters are enriched for at least one pathway in some pathway database such as GO, Reactome or KEGG [12, 1, 13, 20–22]. We look at what proportion of the biclusters returned by each algorithm are enriched for at least one Reactome pathway [23, 24] (restricted to Reactome pathways containing at least one of the 106 genes knocked out in this experiment), measured by Fisher's one-tailed hypergeometric test, with p-values adjusted for multiple testing by the Benjamini-Yekutieli adjustment [25].

#### S3.6 Biclustering in real datasets

We identify, for each knocked-out gene, the bicluster which a algorithm has recovered which most closely matches the samples from that knockout genotype, using the $F_1$ score. Then we look at whether this bicluster is also enriched for genes that share a pathway with the knocked-out gene, using Fisher's hypergeometric test. This evaluation of simultaneous sample clustering and gene clustering is one aspect of our study that is unique among comparisons of biclustering algorithms, and we think it is a very important inclusion.

### S4 Post-processing

FABIA, SDA, SNMF and nsNMF return many biclusters containing all genes and all samples, as shown in Figure S11 and Figure S12. Without post-processing, these algorithms would thus receive very poor scores. We hypothesised that many values in the matrices defining these biclusters were close to 0, and by setting these small values to 0 we would reveal the underlying structure of the biclusters and thus enable meaningful evaluation.

We explored the effect of setting elements of $X$ and $B$ to zero if they were below a certain threshold, with various choices of threshold $\alpha$. Some algorithms, most notably BicMix, return sample loadings $X_{ik}$ and gene loadings $B_{jk}$ on very different scales. To avoid having to account for this imbalance for each algorithm individually, we scaled the vectors $x_k$ and $b_k$ for each bicluster to have $L_2$ norm equal to 1 before removing any elements. Thus, when using a threshold $\alpha$, we set any elements satisfying the following inequality to be 0:

$$\frac{X_{ik}}{\sqrt{\sum_l |X_{lk}|^2}} < \alpha \tag{19}$$

and similarly for any elements of B satisfying:

$$\frac{B_{jk}}{\sqrt{\sum_l |B_{lk}|^2}} < \alpha \tag{20}$$

We found that a threshold of 0.01 was the best choice for all algorithms for simulated datasets (Figure S13), and was also sensible for real datasets (Figure S16 and Figure S14).

It is worth noting that, since thresholding appears to affect different metrics in similar ways, it would have been possible to choose a sensible threshold using only metrics that we would have from a standard real dataset, such as redundancy of recovered biclusters and reconstruction error. Also, although we believe the threshold chosen is suitable for all algorithms, there is potential for further fine-tuning for individual algorithms.

After thresholding, the distribution of factor sizes is less extreme. Figure S18 and S17 show the distribution of bicluster sizes in terms of samples and genes respectively. Interestingly some algorithms such as BicMix-Q and FABIA have a similar distribution regardless of dataset type, whereas others such as BicMix and SNMF seem to adapt to the distribution of bicluster sizes.

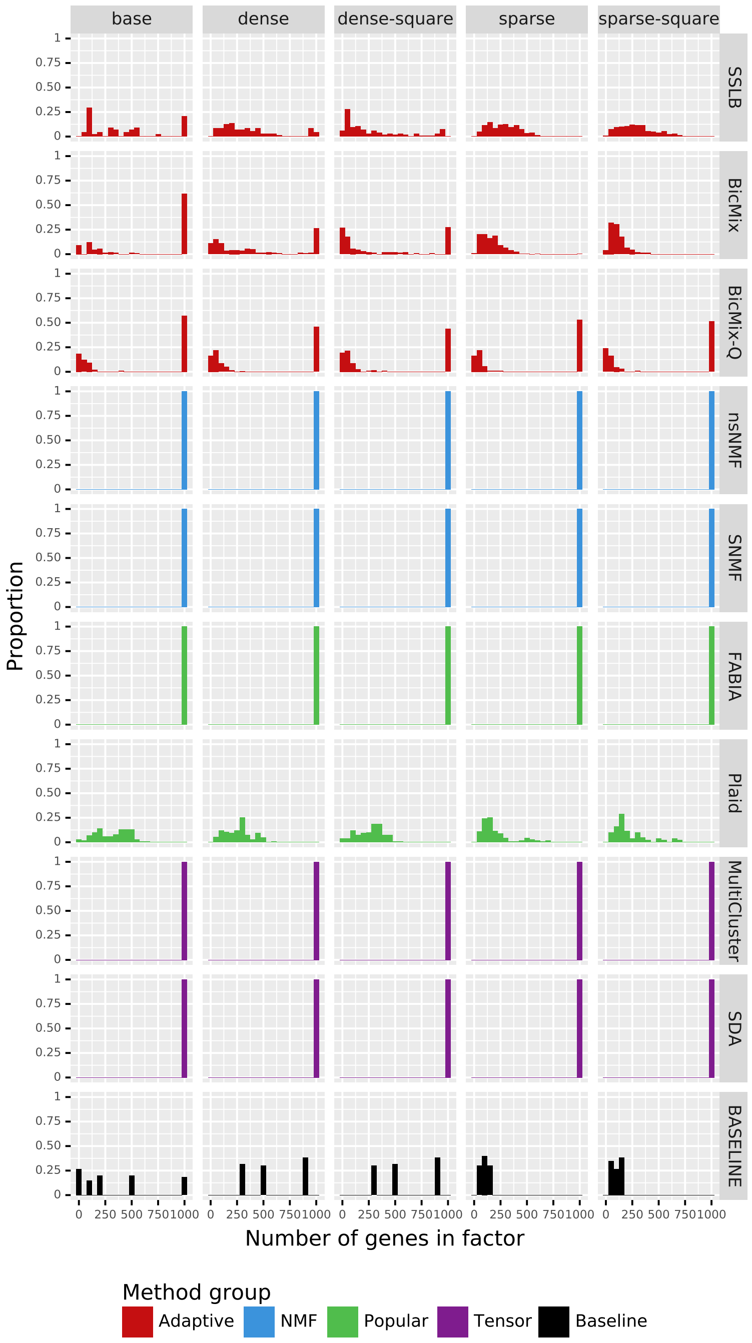


Figure S11: Number of genes contained in biclusters, with no thresholding applied. Each column shows a dataset with different distribution of bicluster sizes. Each graph in the grid shows the relative frequency of the number of genes contained in biclusters. The last row (BASELINE) shows the distribution of number of genes in biclusters in the datasets themselves, and the other rows show the distribution for each algorithm in turn. Note that 100% of the biclusters recovered by FABIA, MultiCluster, SDA, SNMF and nsNMF contain all 1000 genes. BicMix, BicMix-Q and SSLB used $K_{init} = 40$ and the remaining algorithms used $K_{init} = 20$.

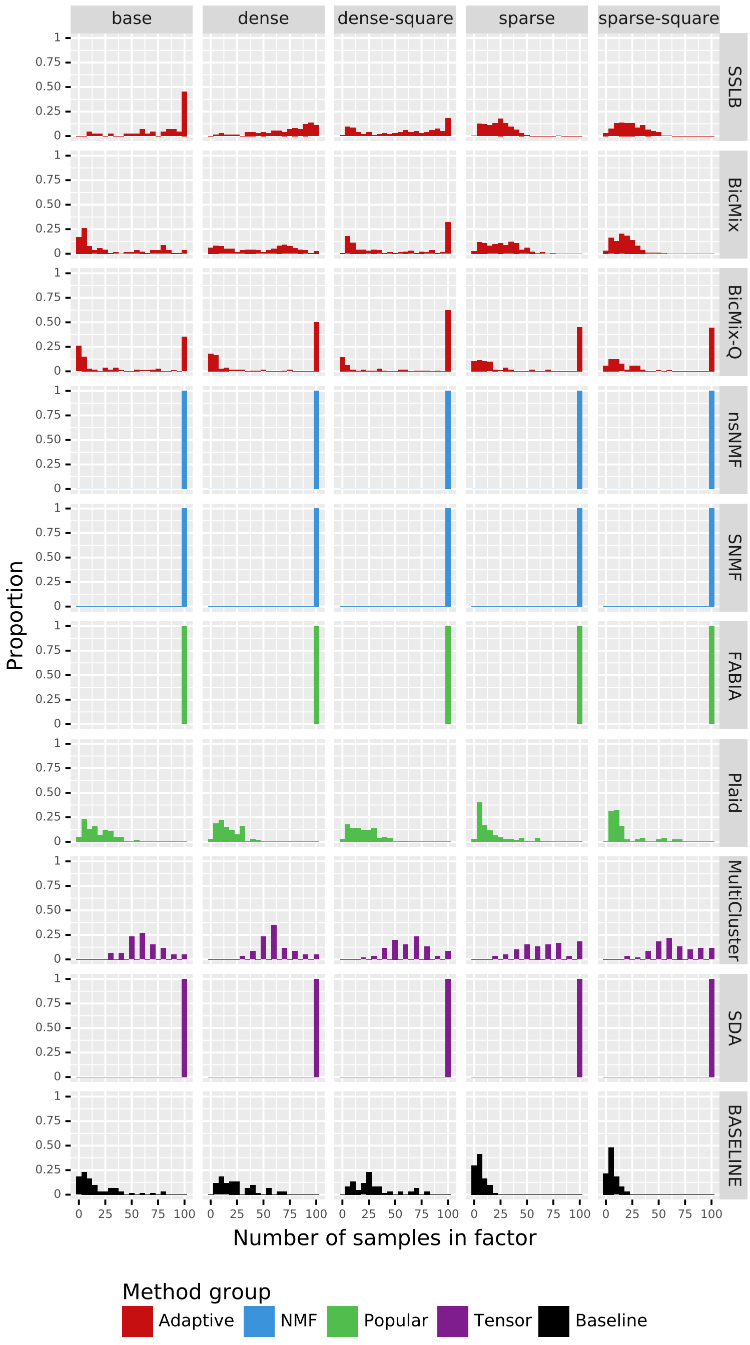


Figure S12: Number of samples contained in biclusters, with no thresholding applied. Each column shows a dataset with different distribution of bicluster sizes. Each graph in the grid shows the relative frequency of the number of samples contained in biclusters. The last row (BASELINE) shows the distribution of number of samples in biclusters in the datasets themselves, and the other rows show the distribution for each algorithm in turn. Note that 100% of the biclusters recovered by FABIA, SDA, SNMF and nsNMF contain all 100 samples. BicMix, BicMix-Q and SSLB used $K_{init} = 40$ and the remaining algorithms used $K_{init} = 20$.

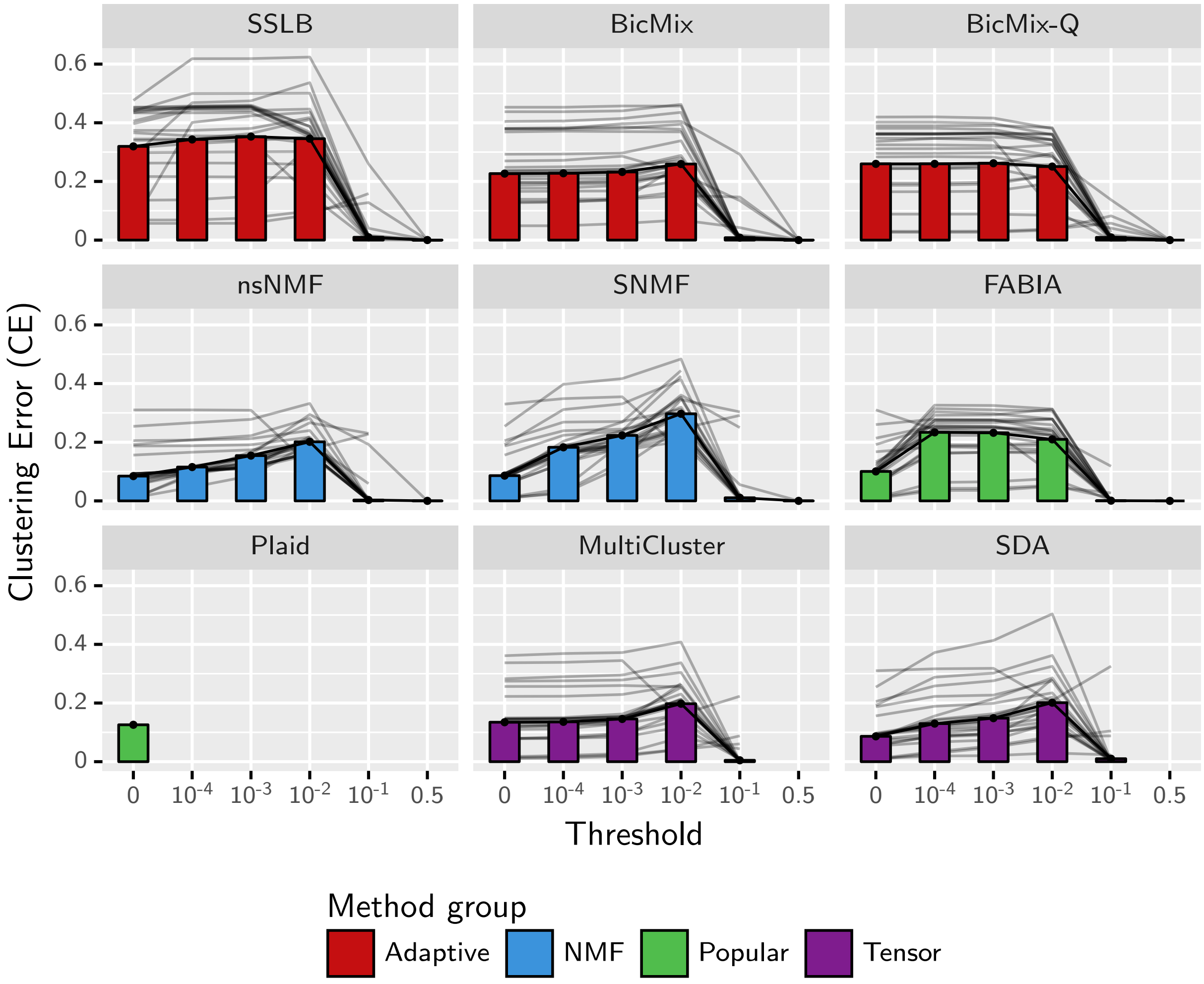


Figure S13: Biclustering accuracy (CE) against the threshold for inclusion in bicluster. Higher values of CE are better. The thresholding process is described in Supplementary Section S4. The median CE across all simulated datasets and all values of $K_{init}$ is shown by the bars. The grey lines show the median for each dataset type. FABIA, MultiCluster, SDA, SNMF and nsNMF show significant improvement in CE as the thresholding becomes stricter and more genes and samples are moved from the recovered biclusters. BicMix and SSLB are relatively unaffected by the thresholding process, until the threshold reaches 0.1. Thresholding could not be applied to Plaid.

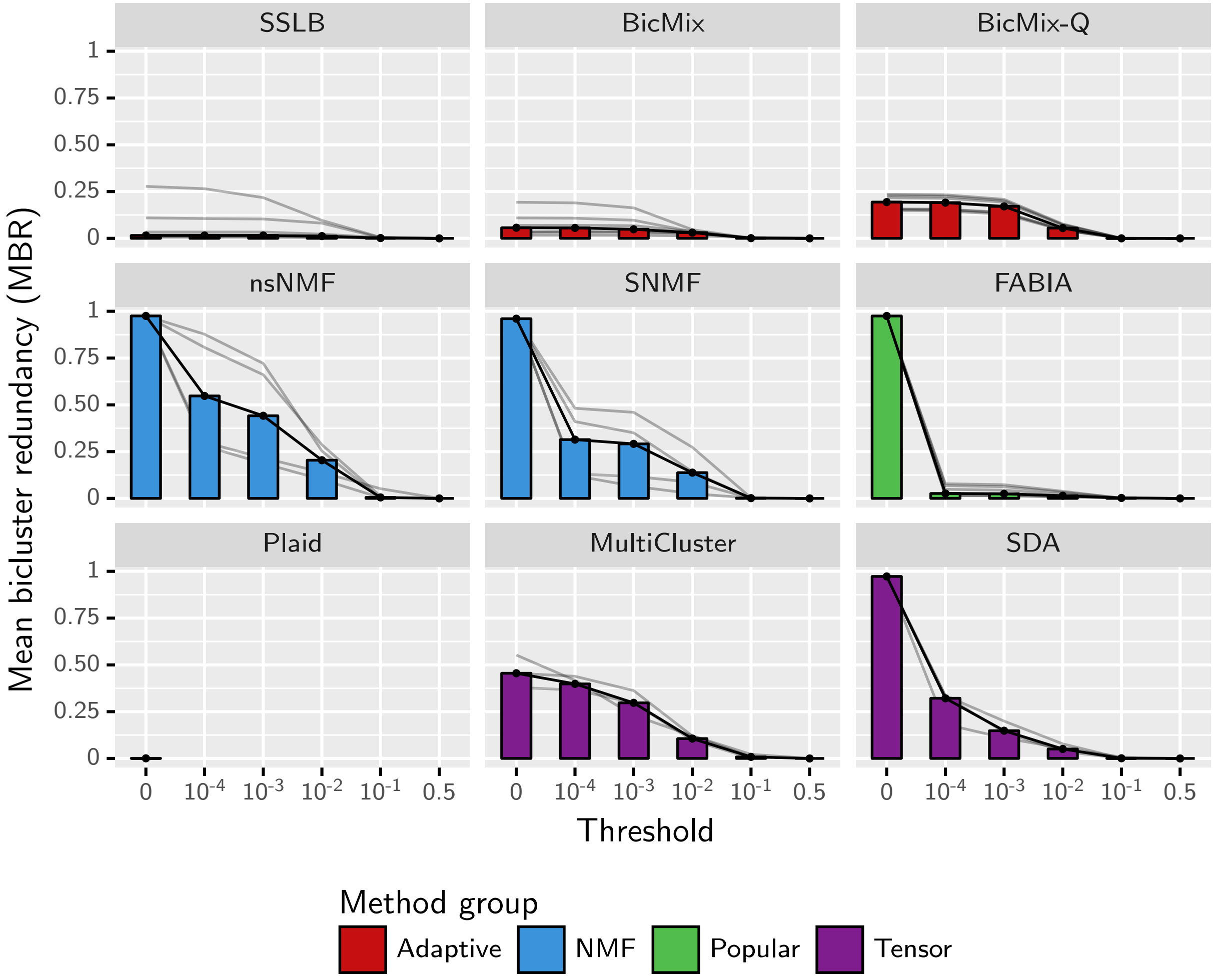


Figure S14: Average similarity of biclusters (MBR) within a run plotted against the threshold for inclusion in bicluster, for IMPC datasets. The redundancy measure is described in detail in Section S3.3. MBR is in the range $[0, 1]$, and a lower value is preferred as a high value suggests that the algorithm is returning many copies of the same bicluster. The thresholding process is described in Section S4. The median of the MBR measure across all IMPC datasets and all values of $K_{init}$ is shown by the bars. The grey lines show the median for each dataset type. Note that without thresholding (threshold 0) the biclusters within each run by FABIA, SDA and nsNMF are almost all identical.

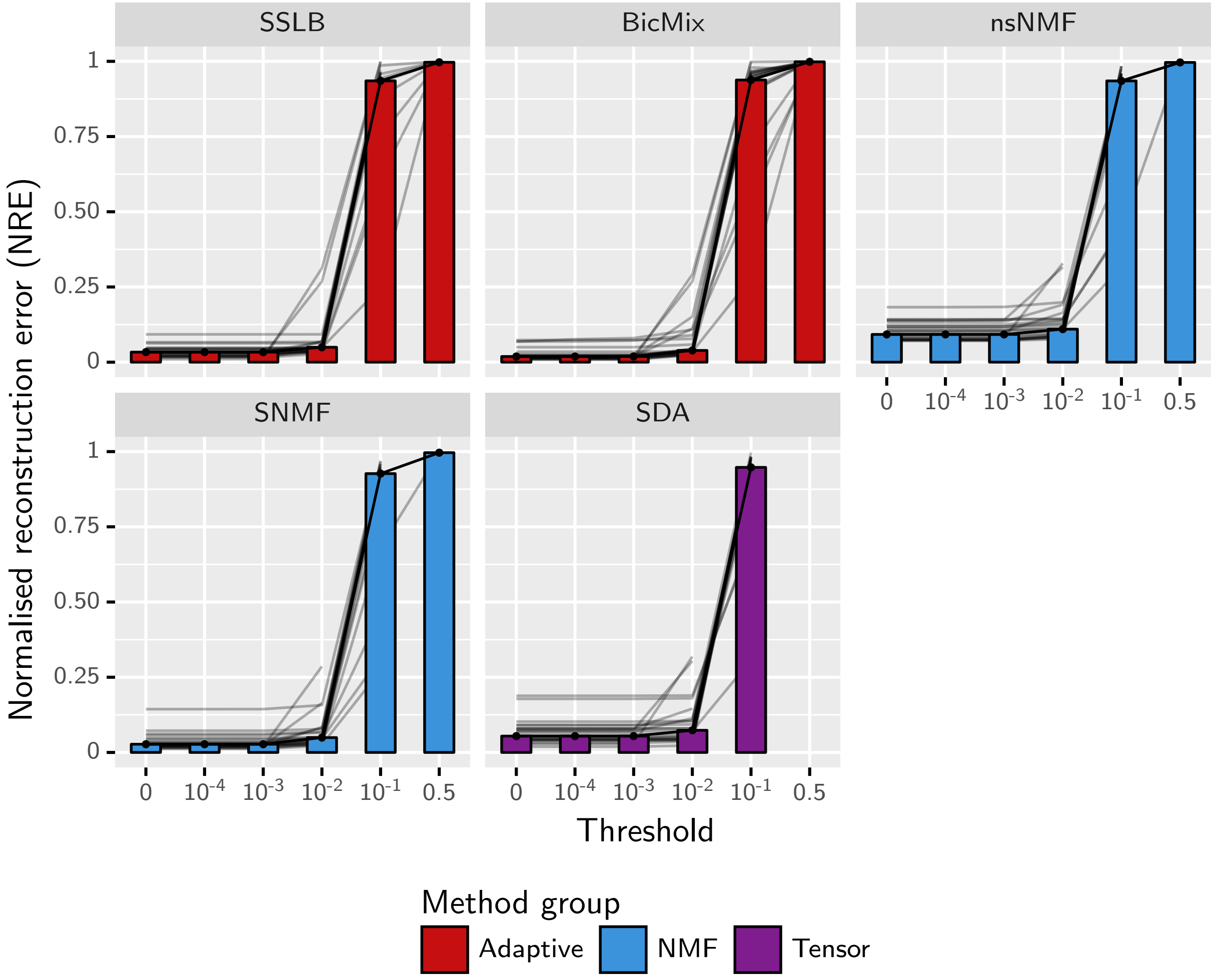


Figure S15: Normalised reconstruction error on simulated datasets, plotted against the threshold for inclusion in bicluster. The thresholding process is described in Section S4. The median of this measure across all simulated datasets and all values of $K_{init}$ is shown by the bars. The grey lines show the median for each dataset type. Note that without thresholding (threshold 0) the biclusters within each run by FABIA, SDA and nsNMF are almost all identical.

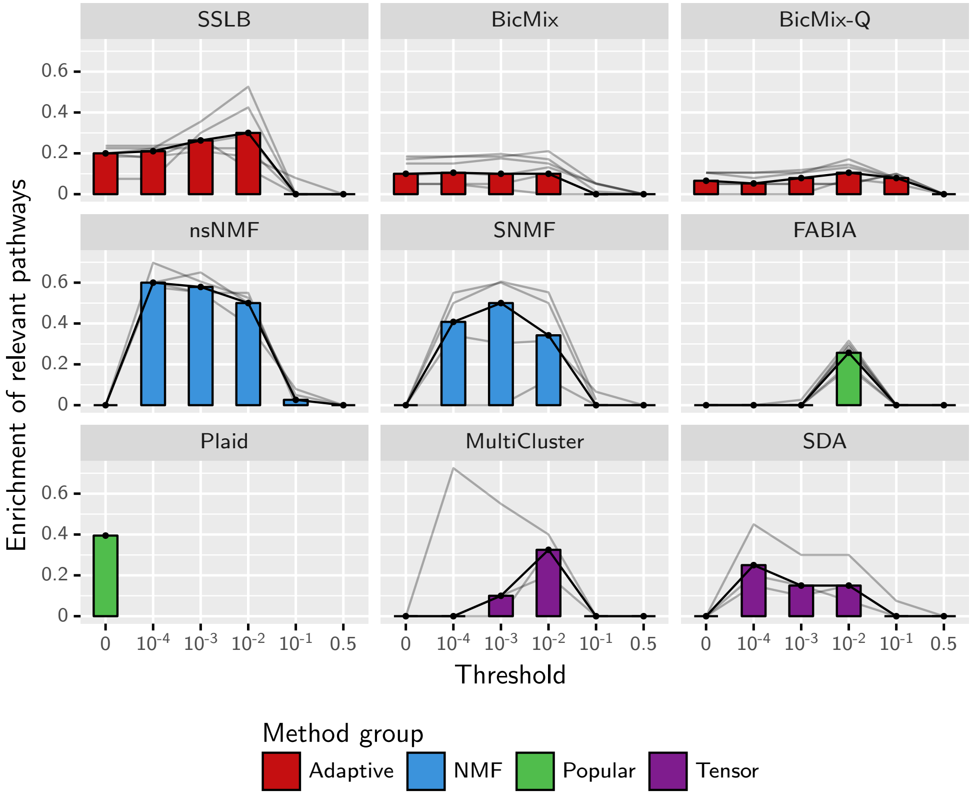


Figure S16: Biclustering performance on IMPC datasets, measured by the mean proportion of knocked-out genes for which the biclusters best matching the samples where the gene was knocked out is enriched for at least one pathway containing the knocked-out gene. A higher score is preferred. Enrichment is measured using the one-tailed hypergeometric test adjusted for multiple testing using the Benjamini-Yekutieli correction, using a threshold for significance of 0.05. The thresholding process is described in Supplementary Section S4. The median of this measure across all IMPC datasets and all values of $K_{init}$ is shown by the bars. The grey lines show the median for each dataset type. As with simulated datasets, the performance of FABIA, MultiCluster, SDA, SNMF and nsNMF show significant improvement as the thresholding becomes stricter.

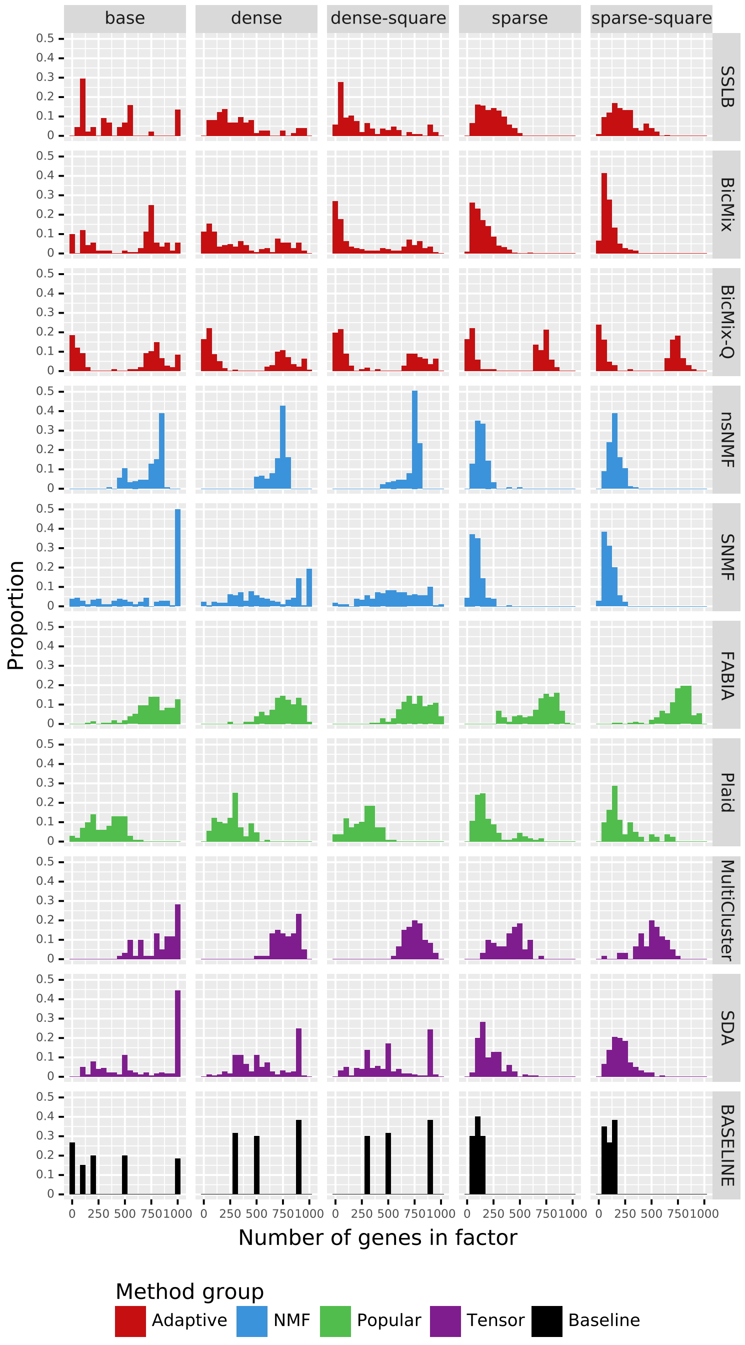


Figure S17: Number of genes contained in biclusters, with thresholding applied as specified in Section S4. Each column shows a dataset with different distribution of bicluster sizes. Each graph in the grid shows the relative frequency of the number of genes contained in biclusters. The last row shows the distribution of number of genes in biclusters in the datasets themselves, and the other rows show the distribution for each algorithm in turn. BicMix, BicMix-Q and SSLB used $K_{init} = 40$ and the remaining algorithms used $K_{init} = 20$.

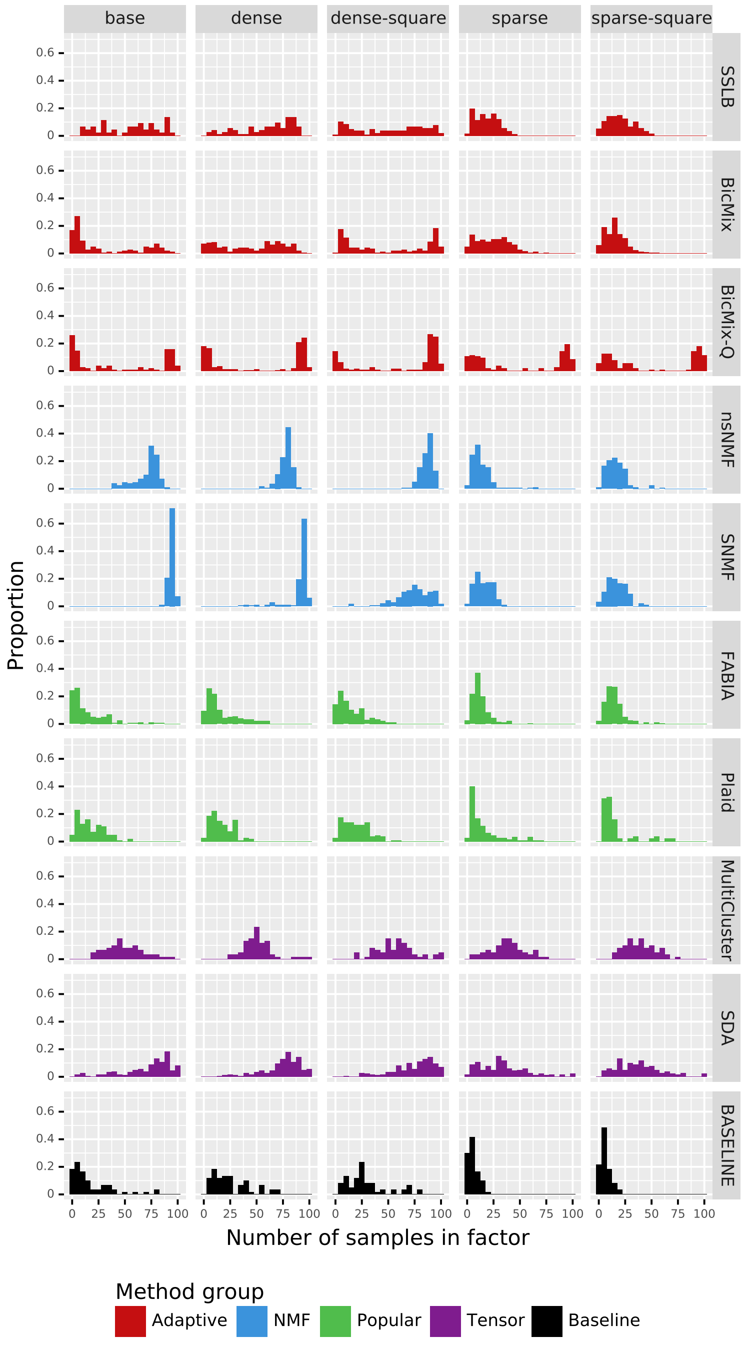


Figure S18: Number of samples contained in biclusters, with thresholding applied as specified in Section S4. Each column shows a dataset with different distribution of bicluster sizes. Each graph in the grid shows the relative frequency of the number of samples contained in biclusters. The last row shows the distribution of number of samples in biclusters in the datasets themselves, and the other rows show the distribution for each algorithm in turn. BicMix, BicMix-Q and SSLB used $K_{init} = 40$ and the remaining algorithms used $K_{init} = 20$.

Table S3: Number of failed runs by each method on each simulated dataset, restricted to runs with the value of $K_{init}$ chosen for analysis. $K_{init} = K$ for all methods except the *Adaptive* methods. Since they are designed to be run with an overestimate of the number of factors, *Adaptive* methods use $K_{init} = 25$, except for the datasets with varying $K$ (the last section of the table), for which they use $K_{init} = K + 10$. Failures are counted as runs that crashed, exceeded the time limit of 12 hours, exceeded the memory limit of 6GB or that failed to find any biclusters. Each cell displays the number of failures as a fraction of total number of attempted runs: *Failures/Attempts*. When all attempts failed, the counts are shown in bold.

| | ***Adaptive*** | | | ***NMF*** | | ***Popular*** | | ***Tensor*** | |
|---|---|---|---|---|---|---|---|---|---|
| **Dataset** | SSLB | BicMix | BicMix-Q | nsNMF | SNMF | FABIA | Plaid | MultiCluster | SDA |
| base | 0 / 9 | 0 / 9 | 0 / 9 | 0 / 9 | 0 / 9 | 0 / 9 | 0 / 9 | 0 / 9 | 0 / 9 |
| N50-T2 | 0 / 9 | 0 / 9 | 0 / 9 | 0 / 9 | 0 / 9 | 0 / 9 | 0 / 9 | 0 / 9 | 0 / 9 |
| N10-T20 | 0 / 9 | 2 / 9 | 0 / 9 | 0 / 9 | 0 / 9 | 0 / 9 | 0 / 9 | 0 / 9 | 0 / 9 |
| N100-T10 | 0 / 9 | 0 / 9 | 0 / 9 | 0 / 9 | 0 / 9 | 0 / 9 | 0 / 9 | 0 / 9 | 0 / 9 |
| N500-T10 | 0 / 9 | 0 / 9 | 0 / 9 | 0 / 9 | 0 / 9 | 0 / 9 | 0 / 9 | 0 / 9 | 0 / 9 |
| G100 | 0 / 9 | 0 / 9 | 0 / 9 | 0 / 9 | 0 / 9 | 0 / 9 | 0 / 9 | 0 / 9 | 0 / 9 |
| G5000 | 0 / 9 | 3 / 9 | 1 / 9 | 0 / 9 | 0 / 9 | 0 / 9 | 0 / 9 | 0 / 9 | 0 / 9 |
| large-K20 | 0 / 9 | 1 / 9 | 0 / 9 | 0 / 9 | 0 / 9 | 0 / 9 | **9 / 9** | 0 / 9 | 0 / 9 |
| Negbin-medium | 0 / 9 | 0 / 9 | 0 / 9 | 0 / 9 | 0 / 9 | 0 / 9 | 0 / 9 | 0 / 9 | 0 / 9 |
| Negbin-high | 0 / 9 | 0 / 9 | 0 / 9 | 0 / 9 | 0 / 9 | 0 / 9 | 0 / 9 | 0 / 9 | 0 / 9 |
| Gaussian | 0 / 9 | 3 / 9 | 0 / 9 | 0 / 9 | 0 / 9 | 0 / 9 | 0 / 9 | 0 / 9 | 0 / 9 |
| Gaussian-medium | 0 / 9 | 2 / 9 | 0 / 9 | 0 / 9 | 0 / 9 | 0 / 9 | 0 / 9 | 0 / 9 | 0 / 9 |
| Gaussian-high | 0 / 9 | 0 / 9 | 0 / 9 | 0 / 9 | 0 / 9 | 0 / 9 | 0 / 9 | 0 / 9 | 0 / 9 |
| noiseless | 0 / 9 | 0 / 9 | 0 / 9 | 0 / 9 | 0 / 9 | 0 / 9 | 0 / 9 | 0 / 9 | 0 / 9 |
| sparse | 0 / 9 | 0 / 9 | 0 / 9 | 0 / 9 | 0 / 9 | 0 / 9 | 0 / 9 | 0 / 9 | 0 / 9 |
| dense | 0 / 9 | 0 / 9 | 0 / 9 | 0 / 9 | 0 / 9 | 0 / 9 | 0 / 9 | 0 / 9 | 0 / 9 |
| sparse-square | 0 / 9 | 0 / 9 | 0 / 9 | 0 / 9 | 0 / 9 | 0 / 9 | 0 / 9 | 0 / 9 | 0 / 9 |
| dense-square | 0 / 9 | 1 / 9 | 0 / 9 | 0 / 9 | 0 / 9 | 0 / 9 | 0 / 9 | 0 / 9 | 0 / 9 |
| K5 | 0 / 9 | 0 / 9 | 0 / 9 | 0 / 9 | 0 / 9 | 0 / 9 | 0 / 9 | 0 / 9 | 0 / 9 |
| K10 | 0 / 9 | 0 / 9 | 0 / 9 | 0 / 9 | 0 / 9 | 0 / 9 | 0 / 9 | 0 / 9 | 0 / 9 |
| K50 | 0 / 9 | 1 / 9 | 0 / 9 | 0 / 9 | 0 / 9 | 0 / 9 | 0 / 9 | 0 / 9 | 0 / 9 |
| K70 | 0 / 9 | 0 / 9 | 0 / 9 | 0 / 9 | 0 / 9 | 0 / 9 | 0 / 9 | 0 / 9 | 0 / 9 |
| large-K100 | 4 / 9 | 7 / 9 | **9 / 9** | 0 / 9 | **9 / 9** | 0 / 9 | **9 / 9** | 0 / 9 | 0 / 9 |
| large-K400 | **9 / 9** | **9 / 9** | **9 / 9** | 0 / 9 | **9 / 9** | **9 / 9** | **9 / 9** | 7 / 9 | **9 / 9** |
| **Total** | 13 / 216 | 29 / 216 | 19 / 216 | 0 / 216 | 18 / 216 | 9 / 216 | 27 / 216 | 7 / 216 | 9 / 216 |

Table S4: Number of failed runs by each method on each IMPC dataset, restricted to runs with $K_{init} = 50$. Failures are counted as runs that crashed, exceeded the time limit of 12 hours, exceeded the memory limit of 6GB or that failed to find any biclusters. Each cell displays the number of failures as a fraction of total number of attempted runs: *Failures/Attempts*. When all attempts failed, the counts are shown in bold.

| | ***Adaptive*** | | | ***NMF*** | | ***Popular*** | | ***Tensor*** | |
|---|---|---|---|---|---|---|---|---|---|
| **Dataset** | SSLB | BicMix | BicMix-Q | nsNMF | SNMF | FABIA | Plaid | MultiCluster | SDA |
| Size factor | 0 / 10 | 0 / 10 | 0 / 10 | 1 / 10 | 0 / 10 | 0 / 10 | **10 / 10** | 0 / 0 | 0 / 0 |
| Log | 0 / 10 | 1 / 10 | 0 / 10 | 0 / 10 | 0 / 10 | 0 / 10 | 0 / 10 | 0 / 0 | 0 / 0 |
| Gaussian | 0 / 10 | 0 / 10 | 0 / 10 | 0 / 0 | 0 / 0 | 0 / 10 | 5 / 10 | 0 / 0 | 0 / 0 |
| Size factor (Tensor) | 0 / 10 | 0 / 10 | 2 / 10 | 0 / 10 | 0 / 10 | 0 / 10 | **10 / 10** | 0 / 10 | 0 / 10 |
| Log (Tensor) | 0 / 10 | 0 / 10 | 1 / 10 | 0 / 10 | 0 / 10 | 0 / 10 | 0 / 10 | 0 / 10 | 0 / 10 |
| Gaussian (Tensor) | 0 / 10 | 0 / 10 | 0 / 10 | 0 / 0 | 0 / 0 | 0 / 10 | 2 / 10 | 0 / 10 | 0 / 10 |
| **Total** | 0 / 60 | 1 / 60 | 3 / 60 | 1 / 40 | 0 / 40 | 0 / 60 | 27 / 60 | 0 / 30 | 0 / 30 |

Table S5: Number of failed runs by each method on each simulated dataset. Failures are counted as runs that crashed, exceeded the time limit of 12 hours, exceeded the memory limit of 6GB or that failed to find any biclusters. Each cell displays the number of failures as a fraction of total number of attempted runs: *Failures/Attempts*. When all attempts failed, the counts are shown in bold.

| | ***Adaptive*** | | | ***NMF*** | | ***Popular*** | | ***Tensor*** | |
|---|---|---|---|---|---|---|---|---|---|
| **Dataset** | SSLB | BicMix | BicMix-Q | nsNMF | SNMF | FABIA | Plaid | MultiCluster | SDA |
| base | 0 / 45 | 2 / 45 | 0 / 45 | 0 / 27 | 0 / 27 | 9 / 27 | 0 / 27 | 0 / 27 | 0 / 27 |
| N50-T2 | 0 / 27 | 0 / 27 | 0 / 27 | 0 / 18 | 0 / 18 | 0 / 18 | 0 / 18 | 0 / 18 | 0 / 18 |
| N10-T20 | 0 / 27 | 3 / 27 | 0 / 27 | 0 / 18 | 0 / 18 | 0 / 18 | 0 / 18 | 0 / 18 | 0 / 18 |
| N100-T10 | 0 / 27 | 0 / 27 | 0 / 27 | 0 / 18 | 0 / 18 | 0 / 18 | 0 / 18 | 0 / 18 | 0 / 18 |
| N500-T10 | 0 / 27 | 0 / 27 | 1 / 27 | 0 / 18 | 0 / 18 | 0 / 18 | 0 / 18 | 0 / 18 | 0 / 18 |
| G100 | 0 / 27 | 0 / 27 | 0 / 27 | 0 / 18 | 0 / 18 | 0 / 18 | 0 / 18 | 0 / 18 | 0 / 18 |
| G5000 | 0 / 27 | 6 / 27 | 2 / 27 | 0 / 18 | 0 / 18 | 0 / 18 | 0 / 18 | 0 / 18 | 0 / 18 |
| large-K20 | 0 / 27 | 2 / 27 | 0 / 27 | 0 / 18 | 0 / 18 | 0 / 18 | **18 / 18** | 0 / 18 | 0 / 18 |
| Negbin-medium | 0 / 27 | 0 / 27 | 0 / 27 | 0 / 18 | 0 / 18 | 0 / 18 | 0 / 18 | 1 / 18 | 0 / 18 |
| Negbin-high | 0 / 27 | 0 / 27 | 0 / 27 | 0 / 18 | 0 / 18 | 0 / 18 | 0 / 18 | 0 / 18 | 0 / 18 |
| Gaussian | 0 / 27 | 3 / 27 | 0 / 27 | 0 / 18 | 0 / 18 | 0 / 18 | 0 / 18 | 0 / 18 | 0 / 18 |
| Gaussian-medium | 0 / 27 | 2 / 27 | 0 / 27 | 0 / 18 | 0 / 18 | 0 / 18 | 0 / 18 | 0 / 18 | 0 / 18 |
| Gaussian-high | 0 / 27 | 0 / 27 | 0 / 27 | 0 / 18 | 0 / 18 | 0 / 18 | 0 / 18 | 0 / 18 | 0 / 18 |
| noiseless | 0 / 27 | 0 / 27 | 0 / 27 | 0 / 18 | 0 / 18 | 0 / 18 | 0 / 18 | 0 / 18 | 0 / 18 |
| sparse | 0 / 27 | 0 / 27 | 0 / 27 | 0 / 18 | 0 / 18 | 0 / 18 | 0 / 18 | 0 / 18 | 0 / 18 |
| dense | 0 / 27 | 5 / 27 | 0 / 27 | 0 / 18 | 0 / 18 | 0 / 18 | 0 / 18 | 0 / 18 | 0 / 18 |
| sparse-square | 0 / 27 | 0 / 27 | 0 / 27 | 0 / 18 | 0 / 18 | 0 / 18 | 0 / 18 | 0 / 18 | 0 / 18 |
| dense-square | 0 / 27 | 1 / 27 | 0 / 27 | 0 / 18 | 0 / 18 | 0 / 18 | 0 / 18 | 0 / 18 | 0 / 18 |
| K5 | 0 / 45 | 0 / 45 | 0 / 45 | 0 / 27 | 0 / 27 | 0 / 27 | 0 / 27 | 0 / 27 | 0 / 27 |
| K10 | 0 / 45 | 1 / 45 | 0 / 45 | 0 / 27 | 0 / 27 | 0 / 27 | 0 / 27 | 0 / 27 | 0 / 27 |
| K50 | 0 / 45 | 1 / 45 | 0 / 45 | 0 / 36 | 0 / 36 | 0 / 36 | 0 / 36 | 0 / 36 | 0 / 36 |
| K70 | 0 / 45 | 0 / 45 | 0 / 45 | 0 / 36 | 0 / 36 | 0 / 36 | 0 / 36 | 0 / 36 | 0 / 36 |
| large-K100 | 13 / 45 | 23 / 45 | 20 / 45 | 0 / 36 | 9 / 36 | 0 / 36 | **36 / 36** | 0 / 36 | 0 / 36 |
| large-K400 | 18 / 45 | 23 / 45 | 25 / 45 | 0 / 36 | 9 / 36 | 9 / 36 | **36 / 36** | 7 / 36 | 9 / 36 |
| **Total** | 31 / 774 | 72 / 774 | 48 / 774 | 0 / 531 | 18 / 531 | 18 / 531 | 90 / 531 | 8 / 531 | 9 / 531 |

Table S6: Number of failed runs by each method on each IMPC dataset. Failures are counted as runs that crashed, exceeded the time limit of 12 hours, exceeded the memory limit of 6GB or that failed to find any biclusters. Each cell displays the number of failures as a fraction of total number of attempted runs: *Failures/Attempts*. When all attempts failed, the counts are shown in bold.

| | ***Adaptive*** | | | ***NMF*** | | ***Popular*** | | ***Tensor*** | |
|---|---|---|---|---|---|---|---|---|---|
| **Dataset** | SSLB | BicMix | BicMix-Q | nsNMF | SNMF | FABIA | Plaid | MultiCluster | SDA |
| Size factor | 0 / 20 | 0 / 20 | 2 / 20 | 1 / 20 | 10 / 20 | 0 / 20 | **20 / 20** | 0 / 0 | 0 / 0 |
| Log | 0 / 20 | 4 / 20 | 4 / 20 | 0 / 20 | 7 / 20 | 0 / 20 | 0 / 20 | 0 / 0 | 0 / 0 |
| Gaussian | 0 / 20 | 0 / 20 | 3 / 20 | 0 / 0 | 0 / 0 | 0 / 20 | 10 / 20 | 0 / 0 | 0 / 0 |
| Size factor (Tensor) | 0 / 20 | 0 / 20 | 4 / 20 | 0 / 20 | 10 / 20 | 0 / 20 | **20 / 20** | 0 / 20 | 1 / 20 |
| Log (Tensor) | 0 / 20 | 0 / 20 | 3 / 20 | 0 / 20 | 7 / 20 | 0 / 20 | 0 / 20 | 0 / 20 | 0 / 20 |
| Gaussian (Tensor) | 0 / 20 | 0 / 20 | 3 / 20 | 0 / 0 | 0 / 0 | 0 / 20 | 4 / 20 | 0 / 20 | 0 / 20 |
| **Total** | 0 / 120 | 4 / 120 | 19 / 120 | 1 / 80 | 34 / 80 | 0 / 120 | 54 / 120 | 0 / 60 | 1 / 60 |

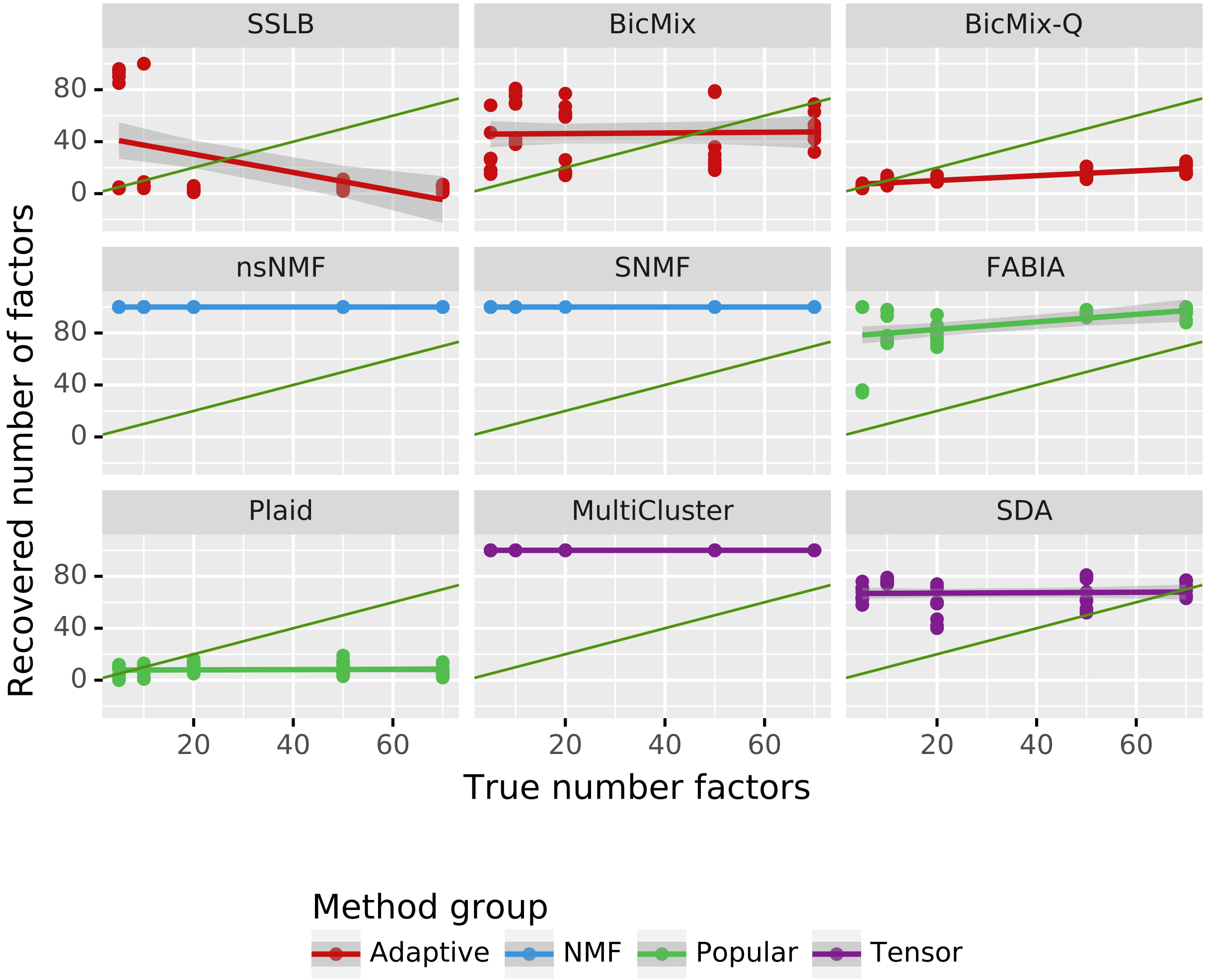


Figure S19: Recovery of K. Number of recovered biclusters plotted against the true number of biclusters when the algorithm was started with 100 biclusters and thresholding was applied as described in Section S4. The green line shows the ideal behaviour, with number of recovered biclusters matching the true number of biclusters exactly. A flatter line indicates poor sensitivity to the true number of biclusters.

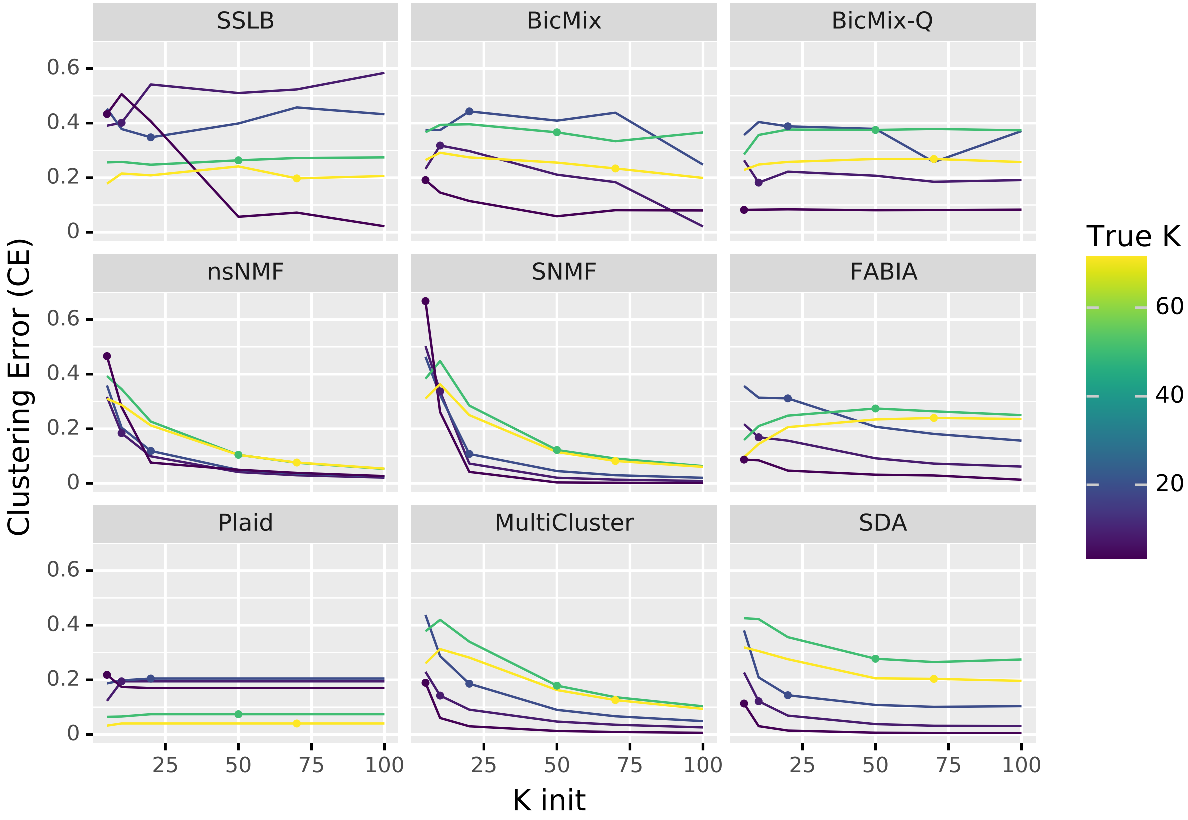


Figure S20: Biclustering accuracy (CE) of algorithms plotted against $K_{init}$. There is a line for each of dataset types *K5*, *K10*, *base*, *K50* and *K70*. For each dataset type there is a point at $K_{init} = K_{true}$ to draw attention to the performance of the algorithm at the true value of K. SNMF, nsNMF and MultiCluster have improved performance at low values of $K_{init}$, even when $K_{true}$ is 50 or 70.

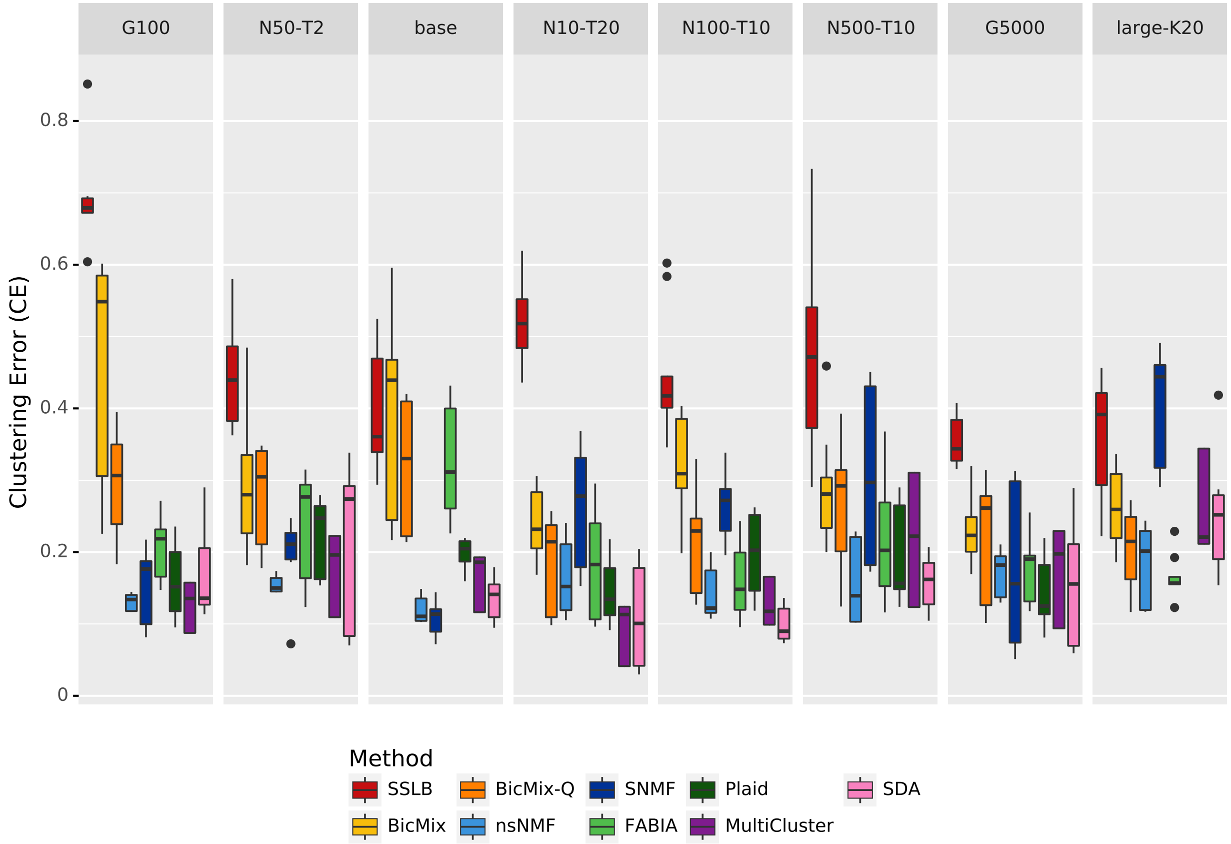


Figure S21: Biclustering accuracy (CE) of algorithms across datasets with a range of sizes. Note that Plaid failed to run on the *large-K20* dataset due to memory constraints. The bar shows median across all three runs on the thee random datasets of each type, with $K_{init} = K$ except for the *Adaptive* algorithms, and the standard thresholding applied.

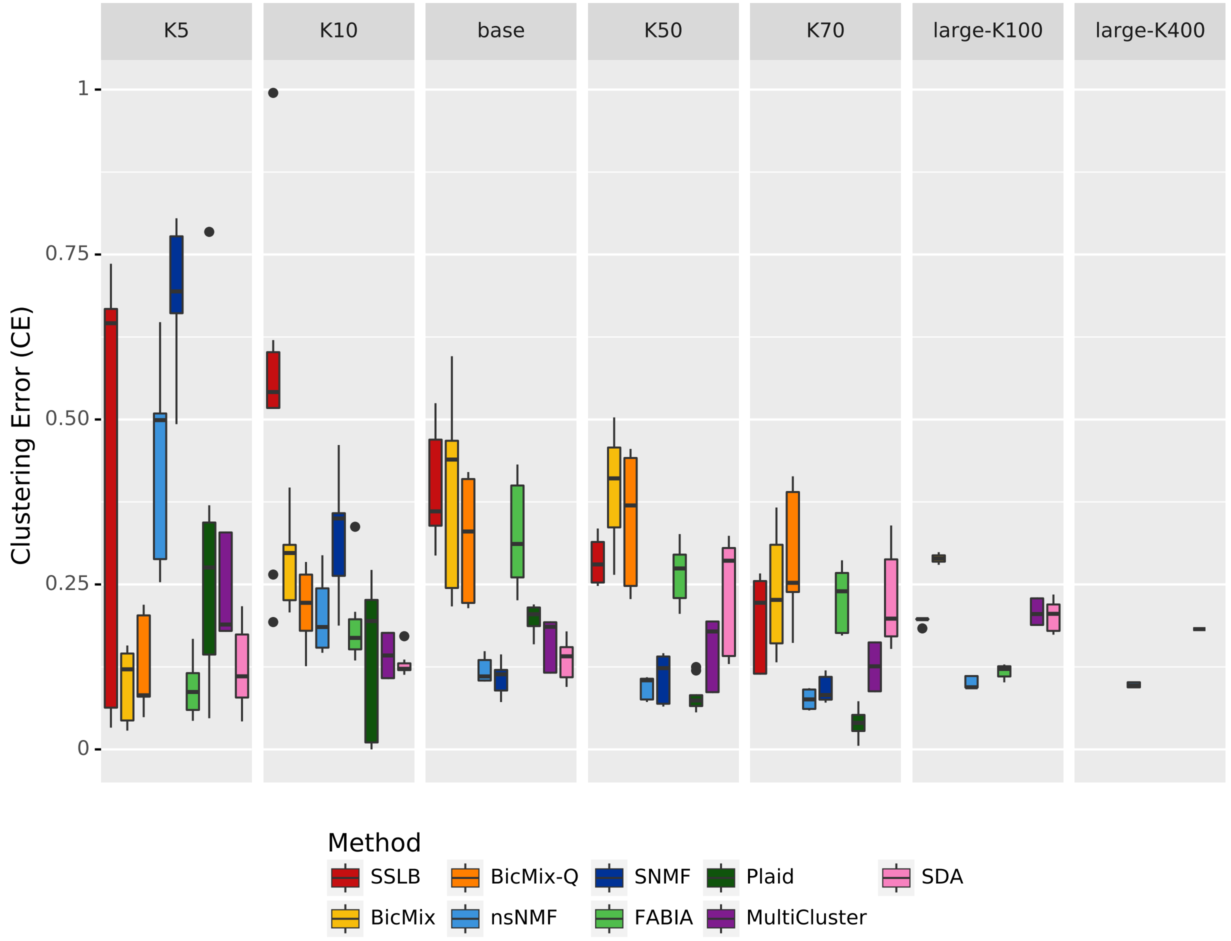


Figure S22: Biclustering accuracy (CE) of algorithms across datasets with different numbers of biclusters. The *base* dataset has 20 biclusters. The bar shows median across all runs on the random datasets of each type, with $K_{init} = K$ except for the *Adaptive* algorithms, and the standard thresholding applied.

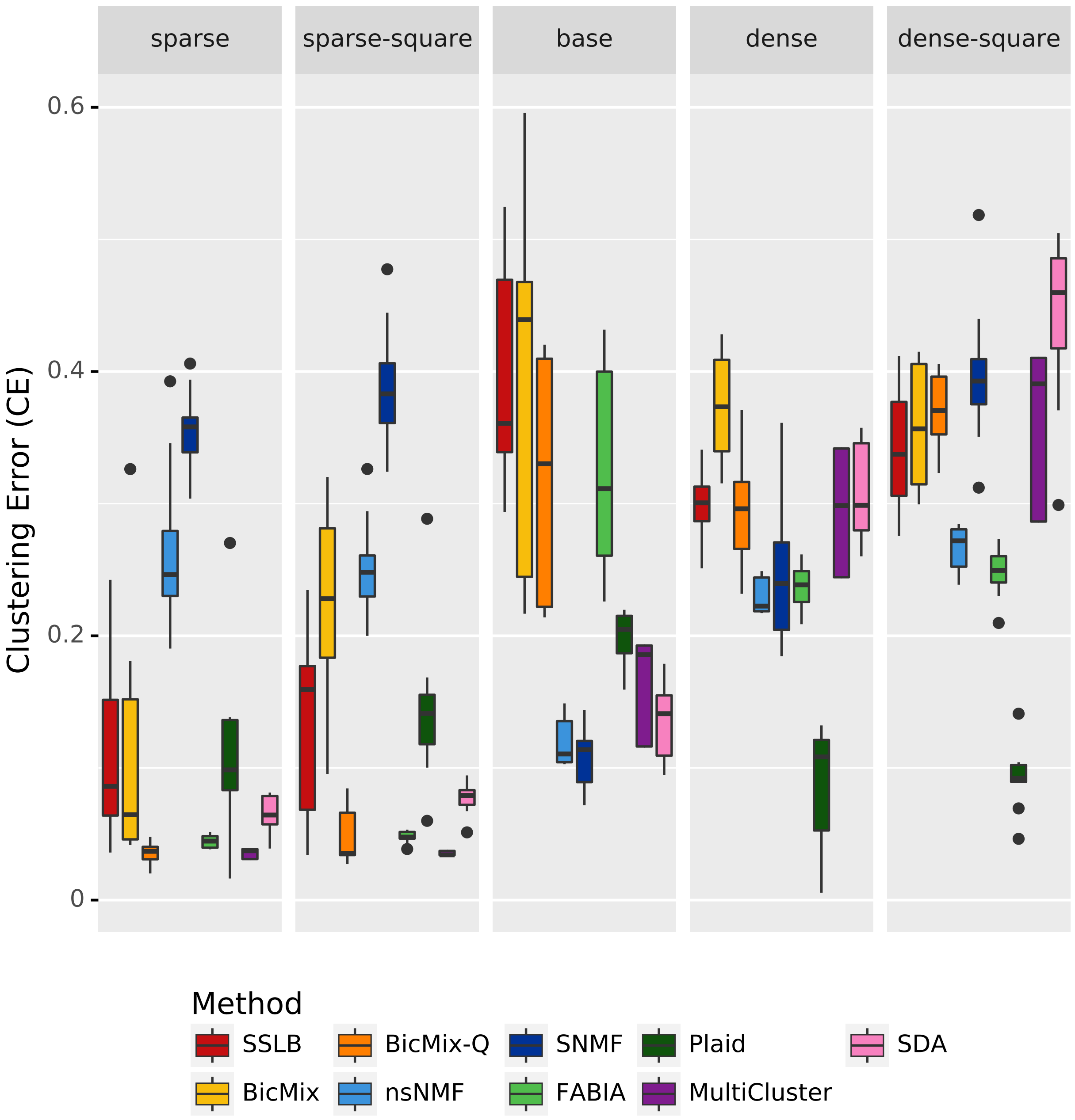


Figure S23: Biclustering accuracy (CE) of algorithms across datasets with different sparsity of biclusters. The distribution of bicluster sizes of the datasets is shown in the first row of Figures S17 and S18. The bar shows median across all runs on the random datasets of each type, with $K_{init} = K$ except for the *Adaptive* algorithms, and the standard thresholding applied.

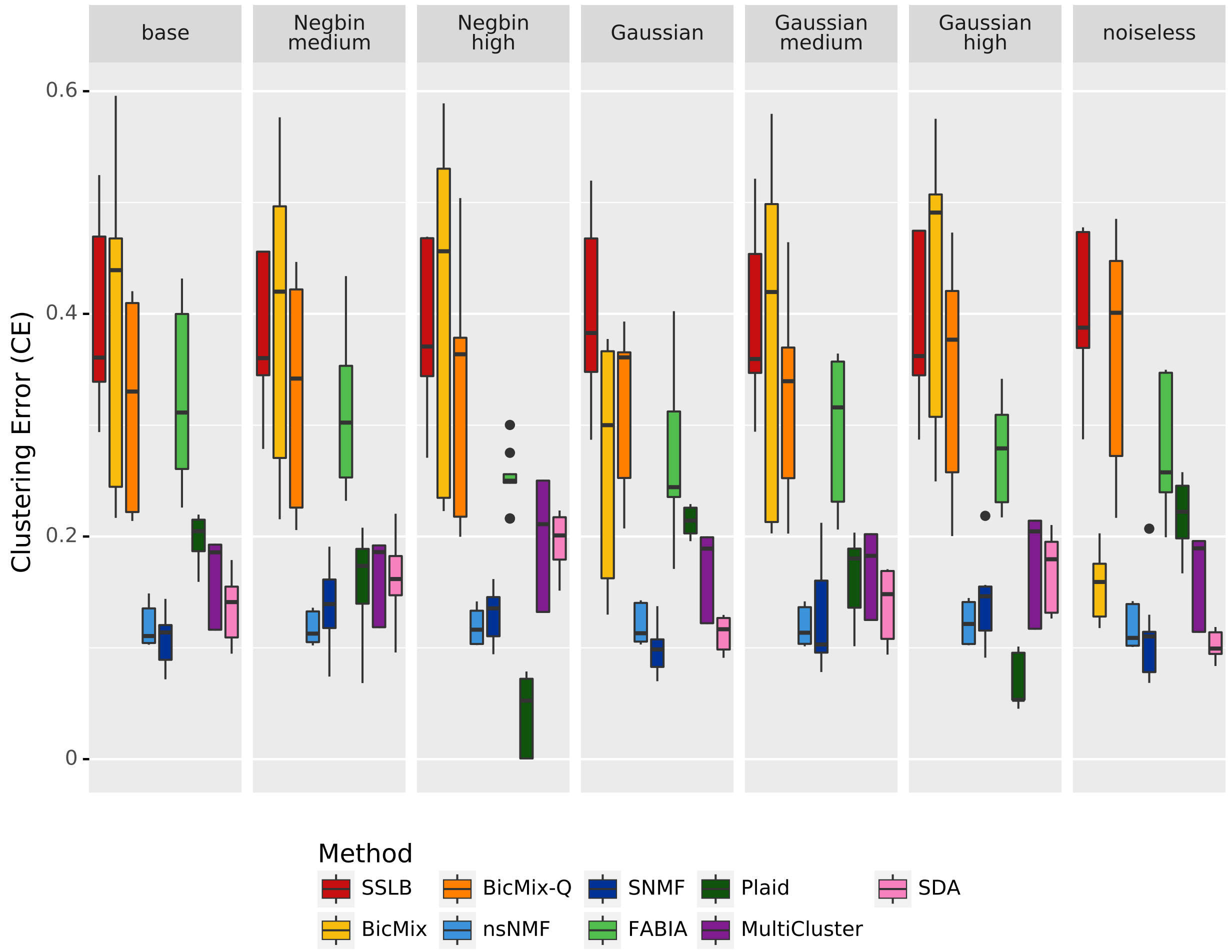


Figure S24: Biclustering accuracy (CE) of algorithms across datasets with different noise distribution. The *base* dataset uses Negative Binomial noise. The bar shows median across all runs on the random datasets of each type, with $K_{init} = K$ except for the *Adaptive* algorithms, and the standard thresholding applied.

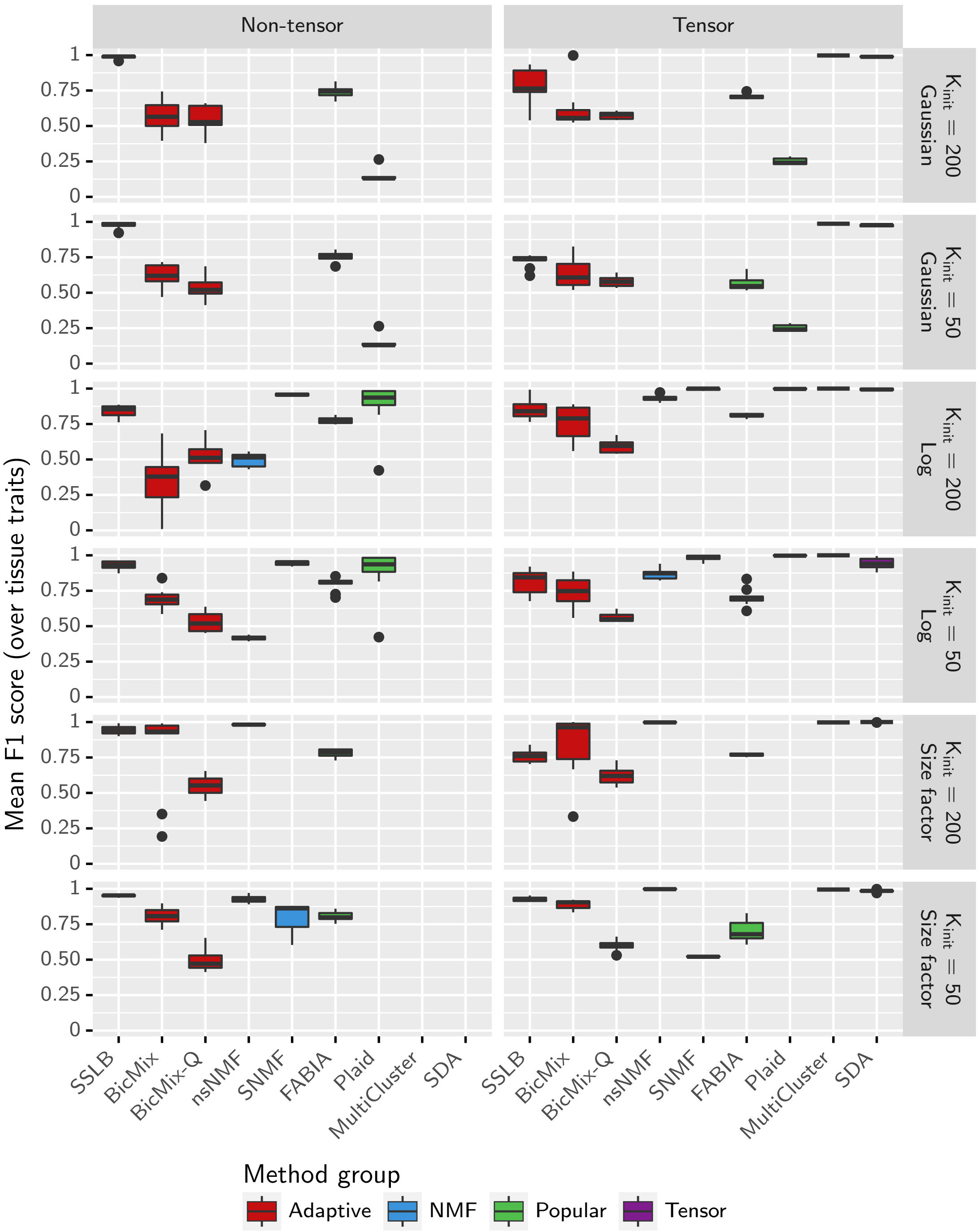


Figure S25: Sample clustering ability on IMPC datasets, measured by the mean over the tissue traits of the maximum $F_1$ score achieved by a bicluster. Standard thresholding is applied and results are shown both for $K_{init} = 50$ and $K_{init} = 200$. Note that MultiCluster and SDA couldn't be run on the non-tensor datasets, nsNMF and SNMF couldn't run on datasets using quantile normalisation and Plaid and SNMF failed to run on the DESeq dataset.

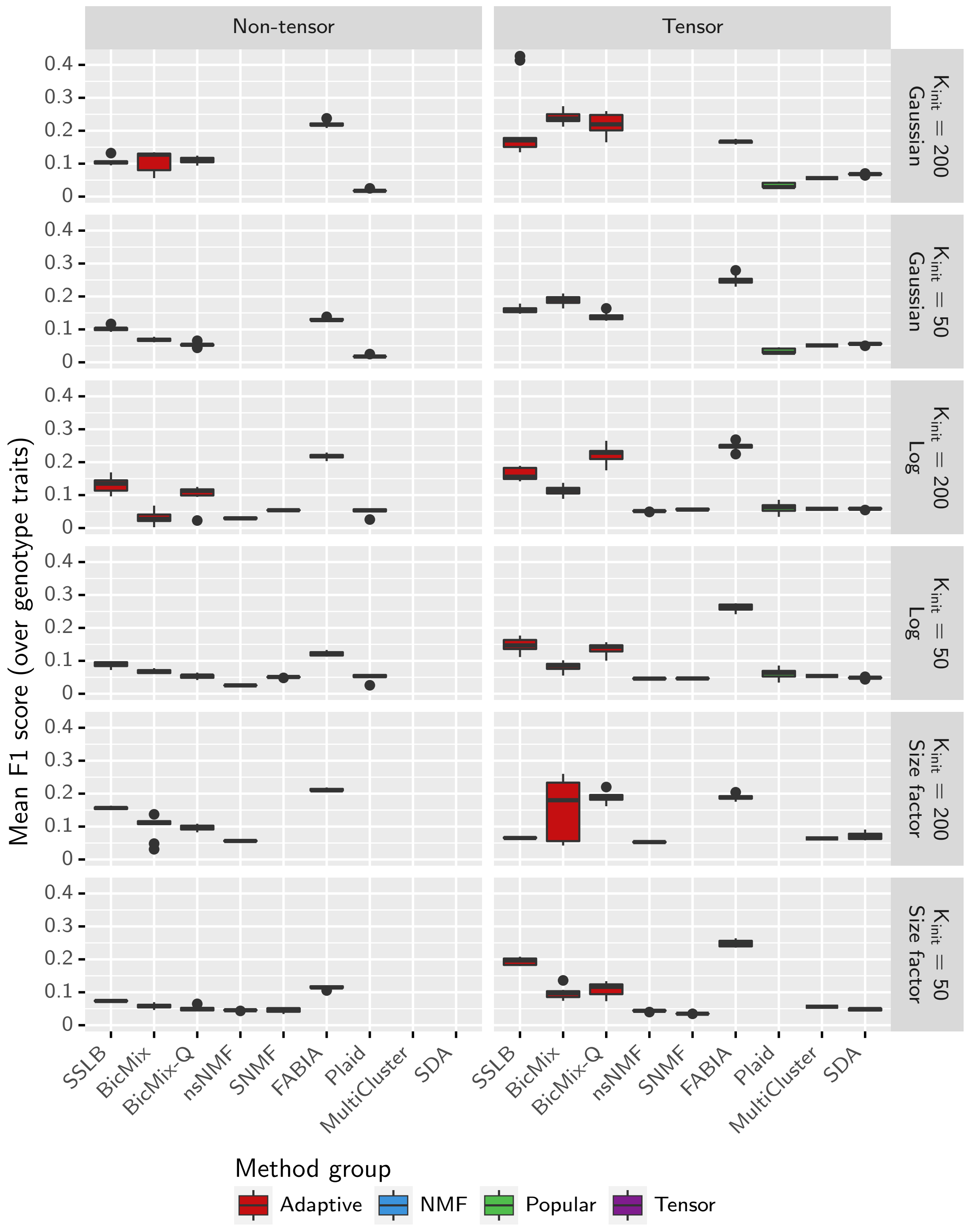


Figure S26: Sample clustering ability on IMPC datasets, measured by the mean over the genotype traits of the maximum $F_1$ score achieved by a bicluster. Standard thresholding is applied and results are shown both for $K_{init} = 50$ and $K_{init} = 200$. Note that MultiCluster and SDA couldn't be run on the non-tensor datasets, nsNMF and SNMF couldn't run on datasets using quantile normalisation and Plaid and SNMF failed to run on the DESeq dataset.

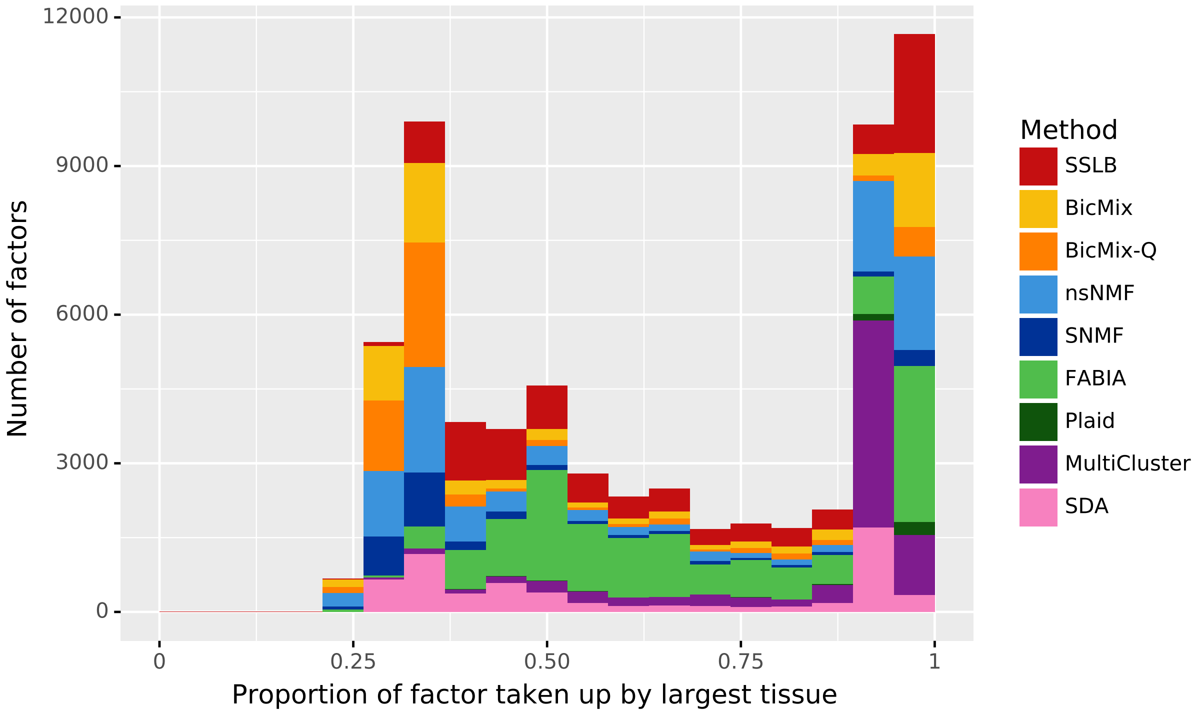


Figure S27: Proportion of bicluster taken up by the largest tissue contained in the bicluster, indicating how frequently the algorithms return biclusters containing multiple tissues. The proportion will be equal to 1 if a bicluster contains only samples from one tissue. The histogram shows that the majority of biclusters recovered contained samples mainly from a single tissue, rather than grouping together samples from different tissues. There is also a large spike around 0.29, which likely corresponds to biclusters containing every single sample as this is the proportion of samples that are from the liver, the most numerous tissue. All returned biclusters across all IMPC datasets are included in the count.

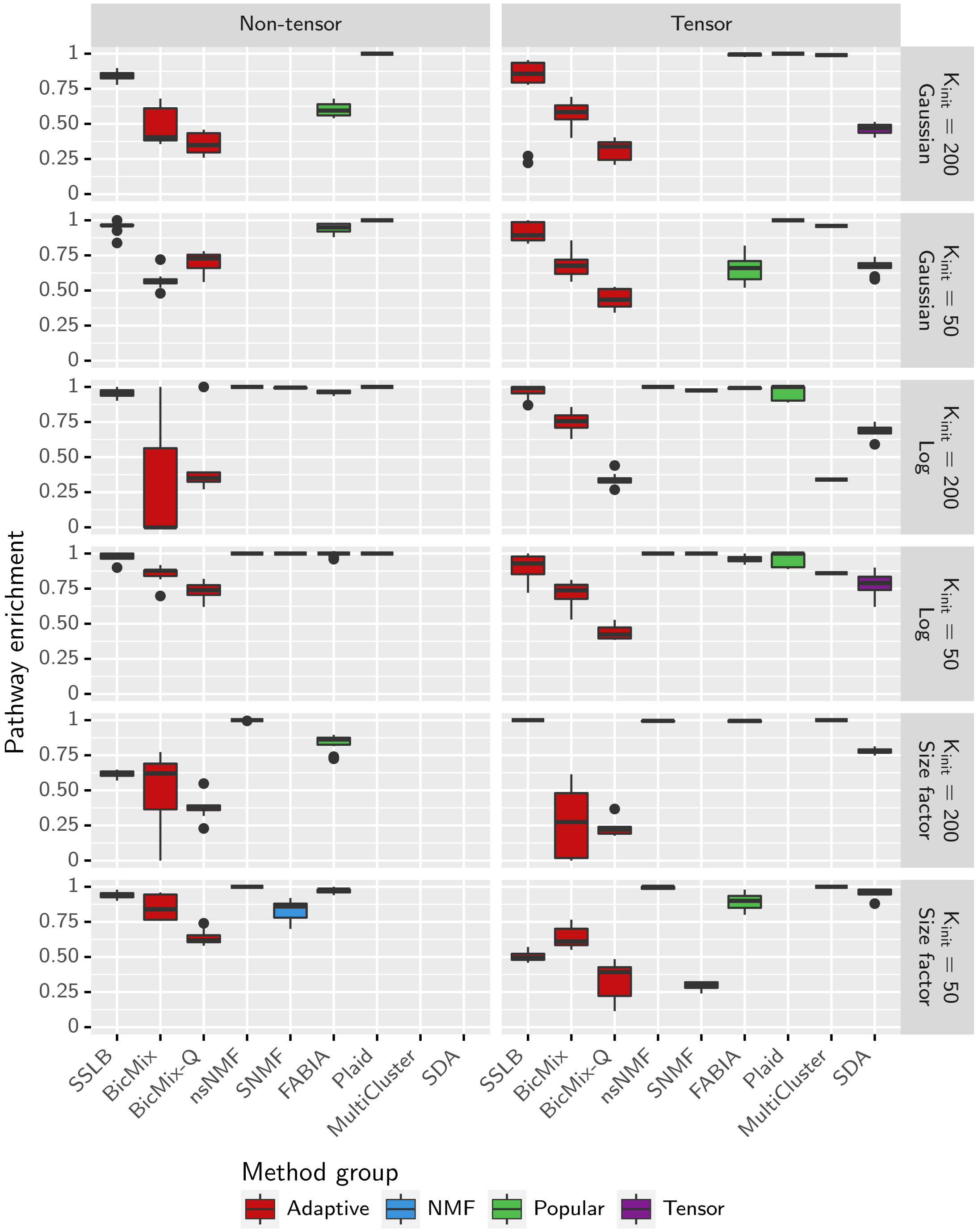


Figure S28: Gene clustering ability on IMPC datasets, measured by the mean proportion of recovered biclusters which are enriched for at least one pathway according to the one-tailed hypergeometric test adjusted for multiple testing using the Benjamini-Yekutieli correction, using a threshold for significance of 0.05. Standard thresholding is applied and results are shown both for $K_{init} = 50$ and $K_{init} = 200$. Note that MultiCluster and SDA couldn't be run on the non-tensor datasets, nsNMF and SNMF couldn't run on datasets using quantile normalisation and Plaid and SNMF failed to run on the DESeq dataset.

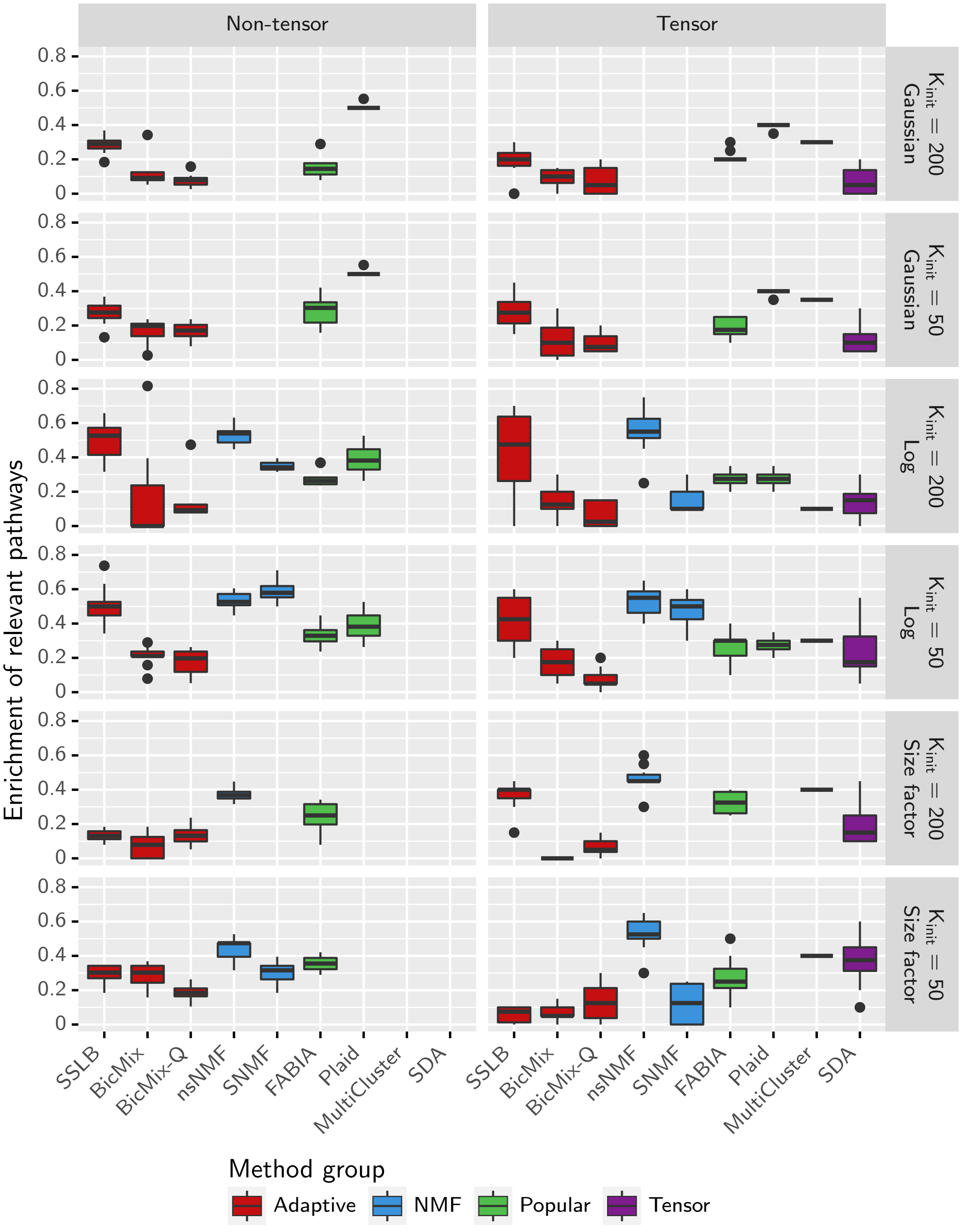


Figure S29: Biclustering ability on IMPC datasets, measured by the mean proportion of knocked-out genes for which the bicluster best matching the samples where the gene was knocked out is enriched for at least one pathway containing the knocked-out gene. Enrichment is measured using the one-tailed hypergeometric test adjusted for multiple testing using the Benjamini-Yekutieli correction, using a threshold for significance of 0.05. Standard thresholding is applied and results are shown both for $K_{init} = 50$ and $K_{init} = 200$. Note that *Tensor* algorithms couldn't be run on the non-tensor datasets, *NMF* algorithms couldn't run on datasets which used quantile normalisation and Plaid failed to run on the dataset which used DESeq's size factor normalisation.

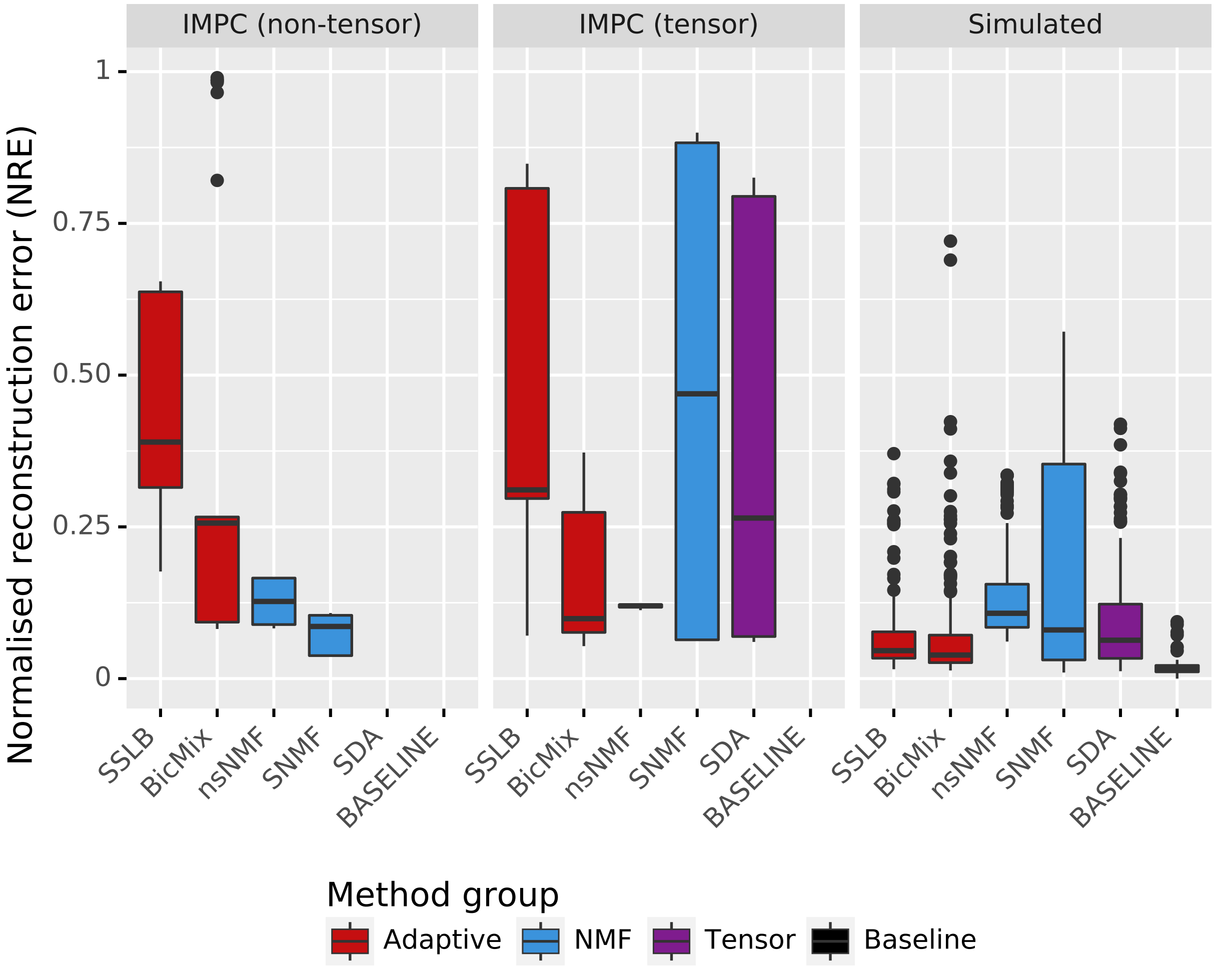


Figure S30: Normalised reconstruction error (NRE) across all datasets. Standard thresholding is applied. The best score is 0. For simulated datasets, BASELINE gives the NRE of the true factorisation $Y = XB^T + \varepsilon$, thus indicating the best performance possible.

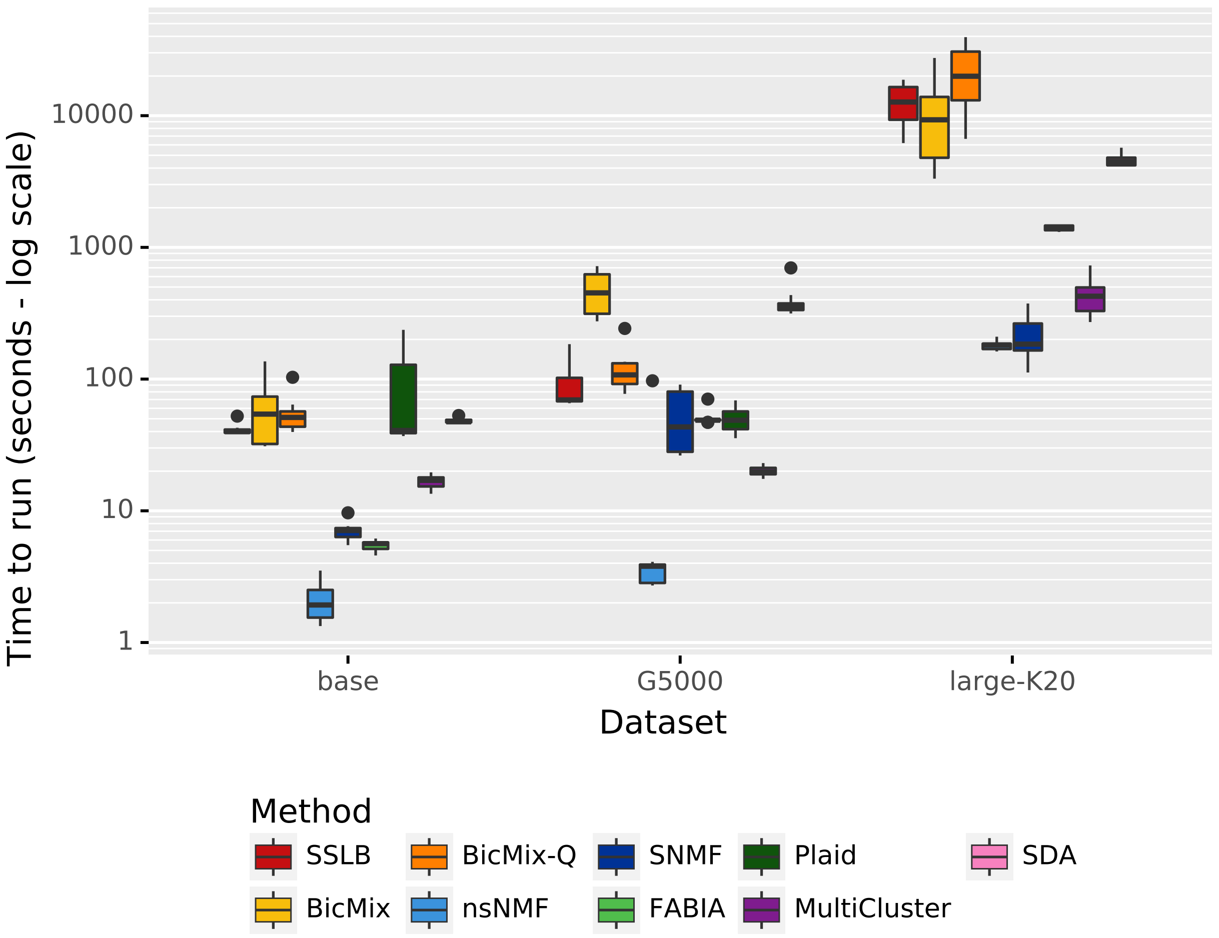


Figure S31: Mean run time across large datasets (log scale). All algorithms were run with $K_{init} = 20$, except for *Adaptive* algorithms which used $K_{init} = 25$. The *base* dataset has 1000 genes and 100 samples, the *G5000* dataset has 5000 genes and 100 samples and the *large-K20* dataset has 10000 genes and 6000 samples. All have 20 latent biclusters. Note that Plaid failed to run on the *large-K20* dataset due to memory constraints.

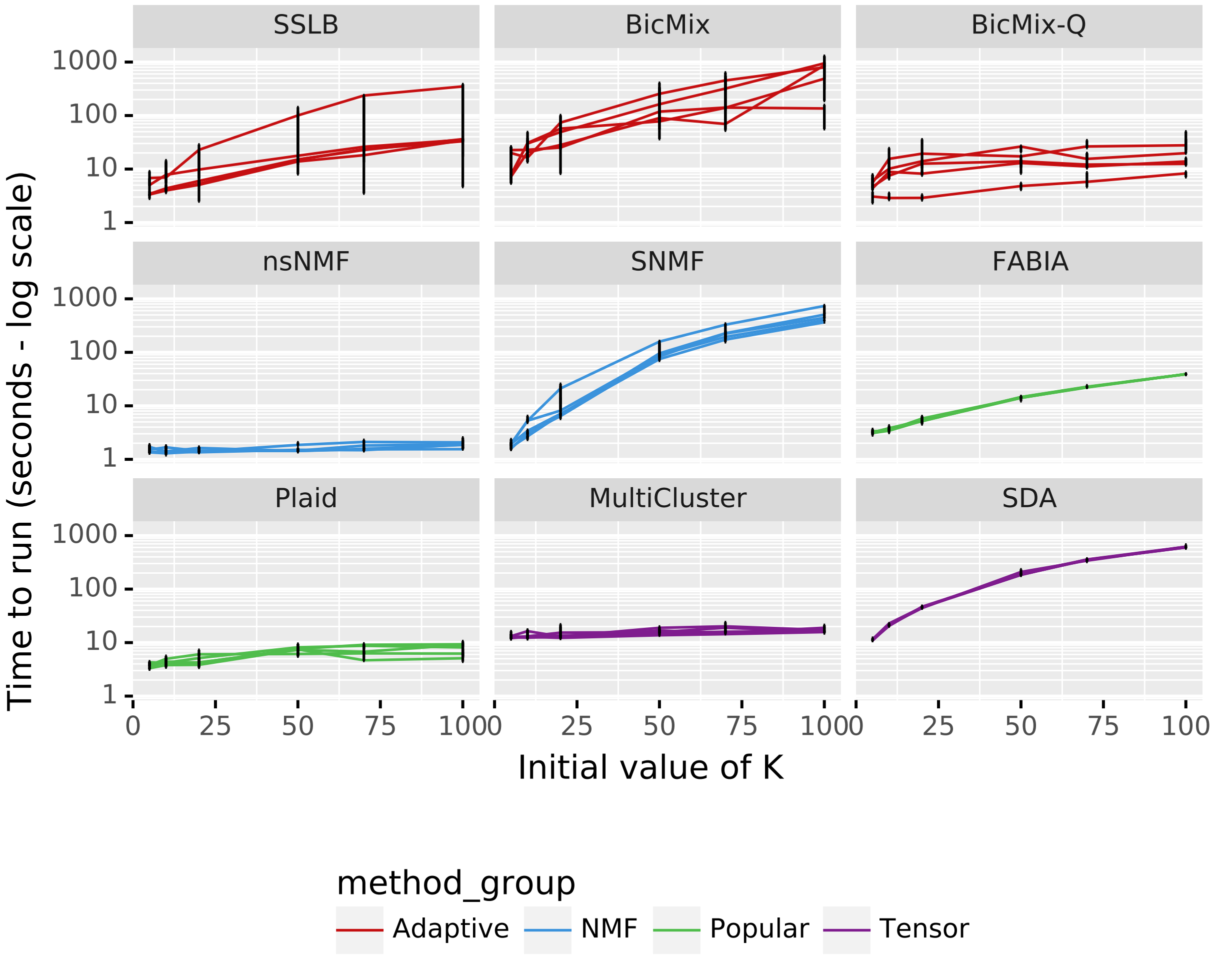


Figure S32: Effect of $K_{init}$ on run time. Each line shows median run time (log scale) for a different dataset type, which differ only in number of latent biclusters (*K5*, *K10*, *base*, *K50* and *K70*). The flatter the line, the less sensitive the runtime is to the choice of $K_{init}$.

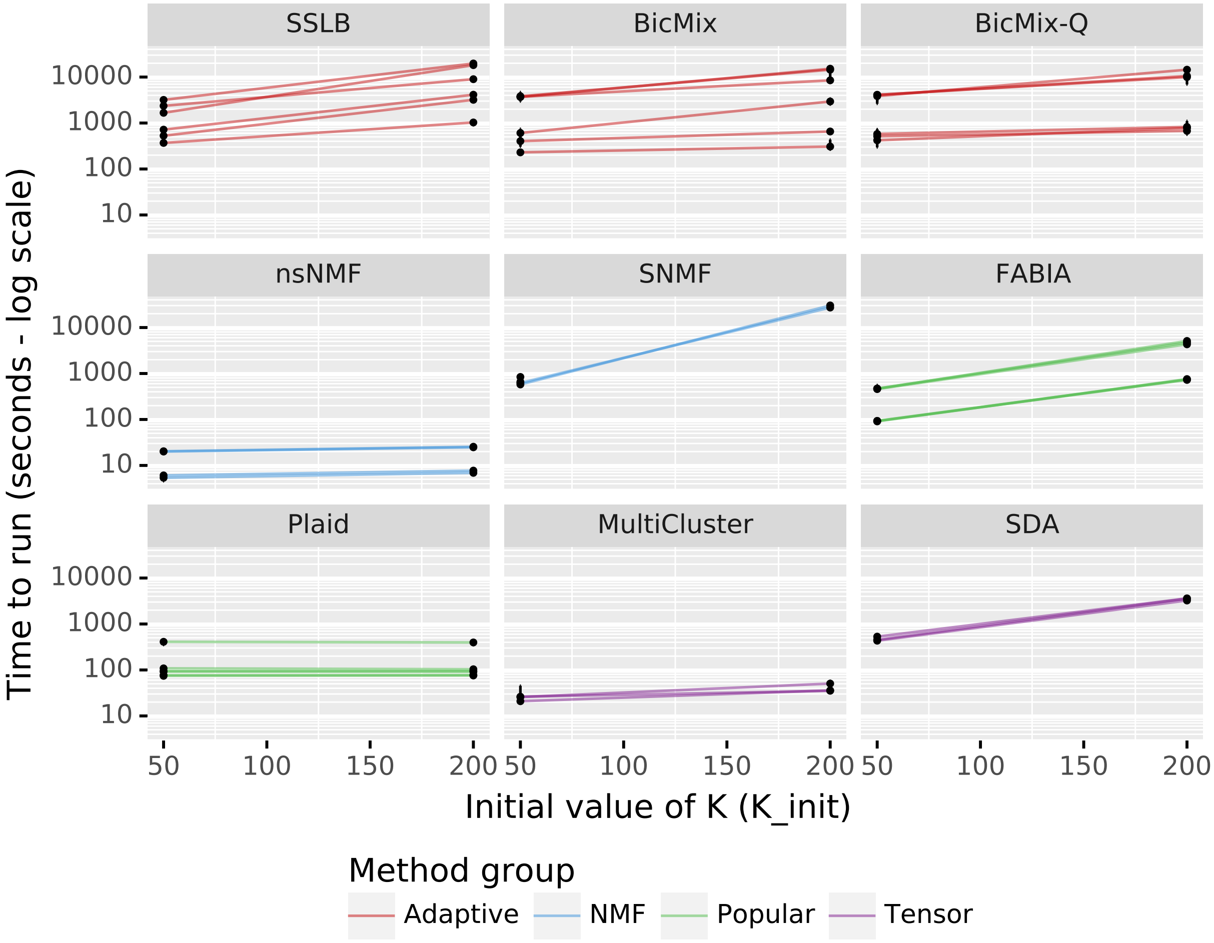


Figure S33: Effect of $K_{init}$ on run time. Each line shows median run time (log scale) for a different IMPC dataset. The flatter the line, the less sensitive the runtime is to the choice of $K_{init}$. Note that MultiCluster and SDA couldn't be run on the non-tensor datasets.

[3] Chuan Gao, Shiwen Zhao, Ian C. McDowell, Christopher D. Brown, and Barbara E. Engelhardt. Differential gene co-expression networks via Bayesian biclustering models. *arXiv:1411.1997 [q-bio, stat]*, November 2014.

[4] Artin Armagan, David B. Dunson, and Merlise Clyde. Generalized Beta Mixtures of Gaussians. *Advances in neural information processing systems*, 24:523–531, 2011. ISSN 1049-5258.

[5] Gemma E Moran, Veronika Ročková, and Edward I George. Spike-and-Slab Lasso Biclustering. page 43.

[15] George Pfundstein. *Ensemble Methods for Plaid Bicluster Algorithm*. Bachelor Thesis, University of München, 2010.

[16] Danilo Horta and Ricardo J.G.B. Campello. Similarity Measures for Comparing Biclusterings. *IEEE/ACM Transactions on Computational Biology and Bioinformatics*, 11(5):942–954, September 2014. ISSN 1545-5963. doi: 10.1109/TCBB.2014.2325016.

[17] Anne Patrikainen and Marina Meila. Comparing Subspace Clusterings. *IEEE Transactions on Knowledge and Data Engineering*, 18(7):902–916, July 2006. ISSN 1041-4347. doi: 10.1109/TKDE.2006.106.

[18] Paul Jaccard. *Nouvelles recherches sur la distribution florale*. Rouge, Lausanne, 1908.
